# Treatment-Resilient Lipid Remodeling Defines Temozolomide Resistance and Failure of Simvastatin Sensitization in Glioblastoma

**DOI:** 10.64898/2026.03.19.712341

**Authors:** Kianoosh Naghibzadeh, Hyunjin Cho, Amir Barzegar Behrooz, Mahboubeh Kavoosi, Marco Cordani, Marek J Los, Stevan Pecic, Rui Vitorino, Carla Vitorino, Amir Ravandi, Shahla Shojaei, Saeid Ghavami

## Abstract

Temozolomide (TMZ) resistance remains a major barrier to durable control of glioblastoma (GB). Our previous studies showed that simvastatin can enhance TMZ-induced cell death in non-resistant GB cells by disrupting autophagosome-lysosome fusion and engaging stress-response pathways, whereas established TMZ-resistant cells maintain impaired autophagy flux and remain refractory to TMZ, simvastatin, and their combination. Here, we asked whether this loss of therapeutic responsiveness is accompanied by a treatment-resilient lipid state. Targeted LC-MS quantified 304 lipid species across 25 analytical classes in non-resistant (NR) and TMZ-resistant (R) U251-mKate cells under control, simvastatin (ST), TMZ, and TMZ-ST conditions. Global heatmap, principal-component, volcano, and exploratory PLS-DA analyses demonstrated persistent NR/R lipidomic separation across all four treatment states. Resistant cells recurrently displayed enrichment of lysophospholipids, selected sphingolipids/glycosphingolipids, and cholesteryl esters, with depletion or redistribution of several glycerophospholipid and diacylglycerol pools. A family-resolved analysis across 19 lipid families showed that resistance status was most strongly associated with phosphatidylinositol (PI; PERMANOVA R²=0.52), phosphatidylglycerol (PG; R²=0.49), lysophosphatidylcholine (LPC; R²=0.46), lysophosphatidylethanolamine (LPE; R²=0.45), ether/plasmalogen phosphatidylcholine (R²=0.45), and phosphatidylcholine (R²=0.41). Structure-informed target prediction of discriminant lipids generated pathway hypotheses involving Rap1, PI3K-Akt, phospholipase D, calcium, PPAR, and lipid-metabolic signaling. Transmission electron microscopy showed persistent vesicle-rich, autophagosome-like architecture in resistant cells across treatment conditions, consistent with the previously established late-stage autophagy defect. These data extend the autophagy-cholesterol model of TMZ resistance to a broader membrane-remodeling program and indicate that failure of statin sensitization is associated with coordinated lipid-family remodeling rather than a single lipid species or pathway. The identified lipid families and cholesterol-storage phenotype provide testable vulnerabilities for future functional validation. An exploratory family-level linear SVM analysis provided an orthogonal proof-of-concept: several membrane and storage-lipid families retained complete NR/R separation when an entire treatment was withheld, but these internal results were interpreted as supplementary evidence rather than as a validated classifier.

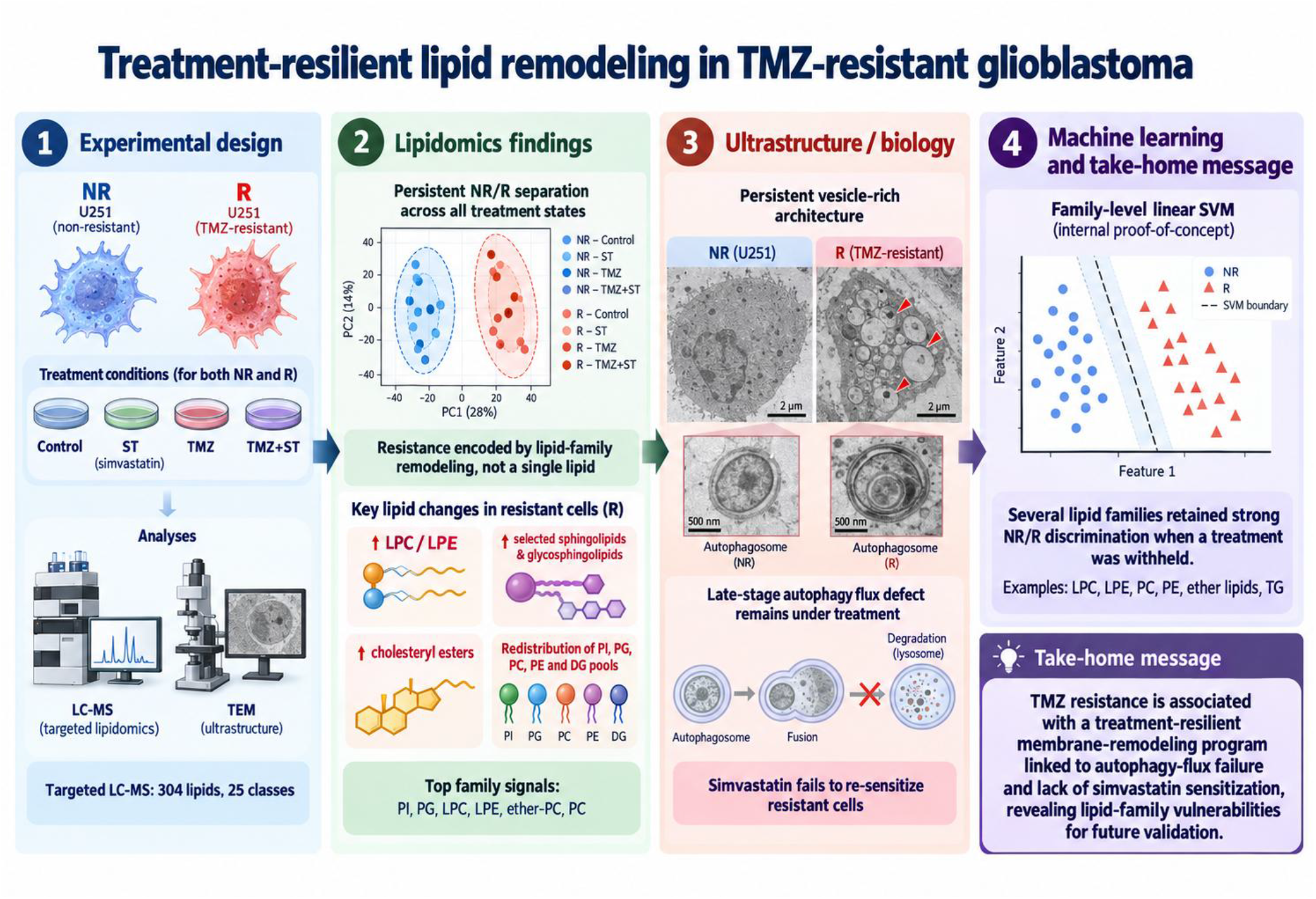

Graphical Abstract: pid analysis and machine learning found distinct fat patterns in TMZ-resistant brain cancer cells grown in the lab. These patterns persisted under treatment alongside faulty cell recycling, suggesting possible targets for further laboratory tests

## 1. Introduction

Glioblastoma (GB) is the most aggressive primary malignant brain tumor in adults and remains associated with poor long-term survival despite maximal surgical resection, radiotherapy, and temozolomide (TMZ)-based chemotherapy [1,2]. Acquired or intrinsic resistance to TMZ is a central cause of treatment failure and recurrence, reflecting not only canonical DNA-repair mechanisms but also broader stress-adaptive programs that preserve tumor-cell viability under continued therapy [3–5].

Drug repurposing has therefore been explored as a strategy to expose metabolic or survival dependencies that coexist with TMZ resistance. Statins are clinically established inhibitors of the mevalonate pathway and can affect tumor-cell survival through both cholesterol-dependent and pleiotropic mechanisms, including altered prenylation, membrane organization, mitochondrial stress, autophagy, and apoptosis [6,7]. In our previous work, simvastatin increased TMZ-induced death in GB models and blocked autophagosome-lysosome fusion, identifying late-stage autophagy as an actionable component of the TMZ response in non-resistant cells [8]. A subsequent study from our group showed that simvastatin-TMZ co-treatment also engages the unfolded protein response while inhibiting autophagy flux, further linking metabolic stress, proteostasis, and cell-death sensitivity [24,25].

The response changes markedly after TMZ resistance has been established. In our previously characterized resistant U251 model, autophagy flux is impaired, mitochondrial and stress-adaptive phenotypes are remodeled, and simvastatin no longer restores TMZ sensitivity [9–11]. The autophagy-cholesterol study further showed that resistant cells can accumulate selected cholesteryl esters despite suppression of de novo cholesterol synthesis, indicating that cholesterol storage, recycling, uptake, or turnover may be uncoupled from synthetic output in the resistant state [9]. In parallel, BCL2L13-dependent modulation of autophagy and ceramide metabolism altered lipid composition without restoring TMZ sensitivity, arguing that individual lipid pathways can be biologically relevant without being sufficient, by themselves, to explain the resistant phenotype [11]. Together, these observations suggested that resistance may be encoded by a broader lipid-network state rather than by one metabolite or one biosynthetic pathway.

This possibility is biologically plausible because autophagy is intrinsically membrane dependent. Autophagosome initiation, expansion, maturation, and lysosomal fusion require coordinated lipid supply and remodeling, including phosphatidylethanolamine-dependent ATG8/LC3 conjugation and phosphatidylinositol-3-phosphate-dependent recruitment of autophagy machinery [12–16]. TMZ-associated autophagy has long been linked to cytoprotective adaptation and to sensitivity to lysosomal inhibition in malignant glioma [19–21], while independent lipidomic studies show broad lipid remodeling in glioma models [22]. Lipid availability also influences mitochondrial membranes, lysosomal function, vesicle trafficking, oxidative stress, and cell-death susceptibility [13,28]. Thus, a cell that has adapted to chronic TMZ exposure may acquire a lipid composition that simultaneously buffers membrane stress and constrains the response to therapies that normally perturb autophagy.

The present study was designed to test that idea directly across treatment states. We hypothesized that acquired TMZ resistance establishes a treatment-resilient lipid architecture that is detectable before treatment, persists during TMZ or simvastatin exposure, and limits the dynamic membrane remodeling associated with therapeutic response in non-resistant cells. We further predicted that resistance would be represented by coordinated remodeling of lipid families rather than isolated species and that the lipidomic pattern would be accompanied by vesicular ultrastructure and pathway associations consistent with altered membrane trafficking and autophagy regulation. To address these questions, we integrated targeted lipidomics, treatment-stratified multivariate and univariate analyses, family-resolved dimensionality reduction with full-dimensional PERMANOVA, structure-informed target/pathway inference, and transmission electron microscopy. This design extends our earlier autophagy, cholesterol, ceramide, and statin studies by defining the lipidomic architecture that distinguishes resistant from non-resistant GB cells under the same therapeutic perturbations [8–11].

## 2. Materials and Methods

### 2.1 Study design and experimental model

The study compared non-resistant (NR) and TMZ-resistant (R) U251-mKate human glioblastoma cells across four matched treatment states: control, simvastatin (ST), TMZ, and combined TMZ-ST. The resistant subline was the previously characterized U251 TMZ-resistant model used in our autophagy and lipid-metabolism studies [9–11]. The experimental design was intended to separate two related questions: the basal lipid state associated with acquired resistance and the extent to which that state is remodeled, or retained, during pharmacological perturbation.

U251-mKate cells were originally developed by Dr. Marcel Bally (Experimental Therapeutics, British Columbia Cancer Agency, Vancouver, BC, Canada). Cells were cultured in high-glucose Dulbecco’s Modified Eagle’s Medium (DMEM; 4 mg/mL glucose; Corning, 50-003-PB) supplemented with 10% fetal bovine serum (FBS; Gibco, 16000044) and 1% penicillin-streptomycin at 37 °C in a humidified atmosphere containing 5% CO2. Resistant cells were maintained in medium containing 250 μM TMZ to preserve the resistant phenotype; TMZ was removed 24 h before experimental treatment.

### 2.2 Simvastatin and temozolomide treatment

NR and R cells were seeded so that they reached approximately 30-40% and 50-60% confluence, respectively, at treatment initiation. Cells were pretreated with simvastatin (1 μM) for 4 h, followed by TMZ (100 μM) where indicated, and were maintained for 72 h. The four conditions within each resistance state were therefore control, ST, TMZ, and TMZ-ST. These conditions were selected to match the treatment framework used in our previous studies showing simvastatin-mediated potentiation of TMZ-induced death in non-resistant GB cells and failure of this sensitization in the resistant model [8,9,24].

### 2.3 Transmission electron microscopy

Cells were prepared for transmission electron microscopy (TEM) to evaluate intracellular membrane organization and autophagosome-like vesicular structures. Cells were fixed with Karnovsky’s fixative (Sigma-Aldrich, SBR00124) for 1 h at 4 °C. Cell pellets were suspended in 5% sucrose in 0.1 M Sorenson’s phosphate buffer and stored overnight at 4 °C, followed by postfixation with 1% osmium tetroxide in Sorenson’s buffer for 2 h. Samples were dehydrated through graded ethanol, embedded in Embed-812 resin (Sigma-Aldrich, 45347), sectioned (∼200 nm), mounted on copper grids, and contrasted with uranyl acetate and lead nitrate. Imaging was performed using a Philips CM10 transmission electron microscope, using the ultrastructural criteria applied in our published autophagy/mitophagy workflows [17].

TEM was interpreted as morphological evidence of vesicle accumulation and organelle architecture, not as a stand-alone measure of autophagic flux. Flux interpretation was anchored to the previously validated biochemical characterization of this resistant model [9–11].

### 2.4 Targeted LC-MS lipidomics

Lipids were extracted from NR and R cells from all four treatment conditions using a chloroform-methanol procedure (2:1, v/v). Cells were harvested, sonicated, and centrifuged at 1000 × g for 10 min. The supernatant was mixed with internal standards and centrifuged at 3500 rpm for 5 min to separate phases. The organic phase was collected, evaporated under nitrogen, reconstituted in water-saturated butanol followed by methanol containing 10 mM ammonium formate, and clarified at 10,000 × g for 10 min before injection.

Lipids were separated on a Zorbax C18 column. Mobile phases contained 10 mM ammonium formate, with a linear gradient from 0% to 100% phase B over 8 min followed by re-equilibration. Diacylglycerol and triacylglycerol subclasses were resolved under isocratic conditions at 100 μL/min. The column was maintained at 50 °C and 5 μL of sample was injected.

Detection was performed using an AB Sciex 4000 QTRAP triple-quadrupole/linear-ion-trap mass spectrometer in scheduled multiple-reaction-monitoring mode. The targeted panel contained 304 lipid species spanning 25 analytical lipid classes. With the exception of free fatty acids, analytes were detected in positive electrospray ionization mode. Lipids were identified using class-specific precursor ions or characteristic neutral-loss transitions and quantified relative to internal standards added before extraction. Species are reported using the total carbon number and degree of unsaturation of the acyl chains when positional isomers were not resolved.

### 2.5 Global and species-level lipidomic analysis

Global lipidomic structure was assessed using hierarchical clustering/heatmaps and principal component analysis (PCA) after log2 transforming and z-score normalization. Differential abundance between NR and R cells was examined separately within each matched treatment condition using volcano plots. Partial least-squares discriminant analysis (PLS-DA) was used as an exploratory feature-prioritization method; lipids with variable-importance-in-projection (VIP) scores ≥1.4 were retained as high-contribution features. Because the number of biological replicates was small relative to the number of lipid features, PLS-DA was not used to estimate predictive performance and its VIP results were interpreted together with the univariate and family-level analyses rather than as independent evidence of classification accuracy.

### 2.6 Structure-informed target prediction and pathway hypothesis generation

For discriminant lipids selected from the treatment-stratified analyses, canonical SMILES strings were retrieved from the Human Metabolome Database (HMDB). Where several structures were available, a canonical representation was used to standardize downstream queries. Putative human protein targets were inferred with SwissTargetPrediction [18]. The predicted proteins were assembled in STRING using a full network and a minimum interaction score of 0.700 (high confidence), followed by KEGG enrichment analysis.

This workflow was used for hypothesis generation (**Figure 1**). It does not measure pathway activation in the cells and does not establish that a complex lipid directly engages a predicted protein target. Accordingly, pathway results were interpreted only as structure-informed associations that could help prioritize membrane-trafficking, signaling, and metabolic mechanisms for future validation.

#### Analytical workflow

**Figure 1.**
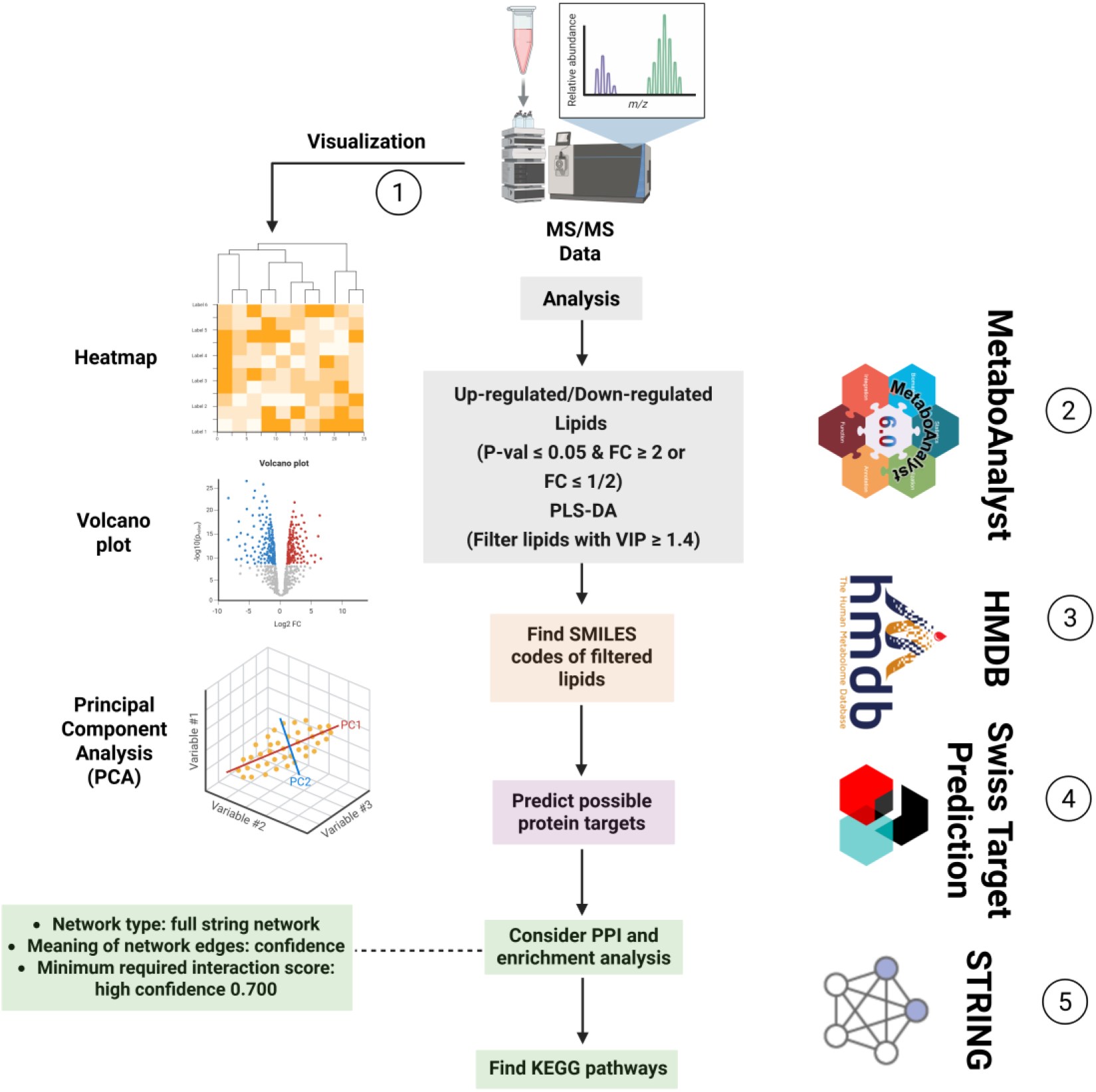
Analytical workflow used for visualization, discriminant-lipid prioritization, SMILES retrieval, structure-informed target prediction, STRING network assembly, and KEGG enrichment. The pathway-inference arm is hypothesis generating and is interpreted separately from experimentally measured lipid abundance.

### 2.7 Family-resolved lipidomic separation

To determine whether resistance was represented by coordinated remodeling of lipid families, the quantified lipidome was partitioned into 19 analysis families and each family was evaluated separately while retaining the fully crossed design of two resistance states across four treatments with three biological replicates per condition (24 samples). Intensities were log2 transformed and z-score normalized. Full-dimensional permutational multivariate analysis of variance (PERMANOVA) was used as the primary quantitative measure of NR/R separation for each family and reported as R². Centroid distance, within-group dispersion, dispersion ratio, and silhouette score were calculated as complementary descriptive measures.

Family-level PCA, uniform manifold approximation and projection (UMAP), isometric feature mapping (ISOMAP), and locally linear embedding (LLE) were used to visualize family-level structure. Because nonlinear embeddings can visually amplify local separation, these projections were treated as descriptive/robustness views; family ranking was based on full-dimensional PERMANOVA rather than on apparent separation in a two-dimensional plot.

### 2.8. Exploratory family-level linear SVM analysis

As a secondary proof-of-concept analysis, a linear support-vector machine (SVM) was fitted separately within each lipid family using log2-transformed family features; z-score standardization was fitted within each training fold and applied to held-out samples. Resistance status (NR versus R) was the binary outcome. Three internal validation schemes were evaluated: leave-one-out cross-validation (LOOCV), in which one sample was held out at a time; leave-one-condition-out (LOCO), in which one treatment-status condition (for example, R-TMZ) was withheld; and leave-one-treatment-out (LOTO), in which all NR and R samples belonging to one treatment (control, ST, TMZ, or TMZ-ST) were withheld together. LOTO was treated as the most stringent internal test because the held-out pharmacological context was absent from model fitting. Performance was summarized using accuracy, balanced accuracy, ROC-AUC, F1 score, Matthews correlation coefficient (MCC), support-vector counts, and SVM decision margins. MCC was used as the principal summary metric because it remains informative when classification outcomes are near the extremes.

For each lipid family, significance under LOTO was assessed by exhaustive enumeration of all C(8,4) = 70 assignments of four conditions to the resistant label, preserving replicate block structure. MCC p-values were adjusted across lipid families using the Benjamini-Hochberg false-discovery-rate procedure. Because the total dataset comprised only 24 samples and no independent external cohort was available, the SVM analysis was explicitly considered exploratory internal validation rather than evidence of a clinically generalizable classifier. The minimum attainable p-value under exhaustive enumeration is 0.014.

#### 2.8.1 Within-family feature stability and reduced-signature sensitivity analysis

Within-family coefficient stability and feature-retention analyses were examined to determine whether the observed family-level discrimination required the full measured lipid set. Lipids were ranked within families by per-fold coefficient mean, standard deviation and sign consistency, and retention curves were used to identify reduced subsets that preserved the LOTO performance of the corresponding full-family model. The resulting reduced signatures were re-evaluated by LOTO and permutation testing. These analyses were used only to assess internal redundancy and robustness; because feature selection and evaluation were performed within the same small dataset and no external cohort was available, reduced signatures were not interpreted as validated lipid biomarkers.

### 2.9 Exploratory quantitative TEM morphometry

The available annotated basal-condition TEM image set was also evaluated quantitatively as an exploratory morphometric analysis. Autophagosomes (AP) and autolysosomes (AL) were enumerated, and AP fraction was defined as AP/(AP + AL). Median individual AP and AL areas were summarized in square micrometers. Between-state differences were reported with Hedges’ g, nominal p-values, and false-discovery-rate-adjusted q values as provided in the morphometric analysis. Because multiple vesicular structures can be nested within the same micrograph or biological preparation, these measurements were treated as descriptive/exploratory and were not used as confirmatory biological-replicate inference.

### 2.10 Statistical analysis and reproducibility

All experimental conditions contained three independent biological replicates, defined as separately cultured and treated cell populations processed independently through lipid extraction and LC-MS analysis. Data are reported as mean ± SD where appropriate. Two-group comparisons used two-tailed Student’s t tests when applicable; multi-group analyses used one-way or two-way ANOVA followed by Tukey’s multiple-comparison test. A two-sided p-value ≤ 0.05 was considered significant. Lipid annotation and class assignment were supported by the LIPID MAPS framework. PERMANOVA effect sizes are reported as R²; embedding analyses were considered descriptive and were not substituted for full-dimensional inference. he analysis code supporting this study is available on GitHub at https://github.com/UIowaJinCho/lipidomics-tmz-tem.

## 3. Results

### 3.1 Resistance produces a treatment-resilient global lipidomic state

We first asked whether acquired TMZ resistance was associated with a basal lipidomic state that remained distinguishable during therapy. Hierarchical clustering of the most strongly changing species showed a clear resistance-associated pattern, with untreated R cells enriched in several LPC, sphingomyelin, and phosphatidylcholine species relative to NR controls. In contrast, the most conspicuous treatment-associated remodeling in NR cells occurred after combined TMZ-ST exposure, where multiple PI and PE species increased relative to the NR baseline (**Figure 1A**).

PCA supported a persistent difference between NR and R lipidomes across the complete dataset (**Figure 1B**). When each treatment was examined separately, NR and R samples remained separated under control, ST, TMZ, and TMZ-ST conditions (**Figure 1C-F**). The combination-treated NR group showed a pronounced displacement and broader spread in the reduced-dimensional representation, whereas the corresponding R samples remained positioned within the resistance-associated region. With three biological replicates per condition, these PCA patterns should be viewed as descriptive evidence of treatment-associated lipid redistribution rather than a direct kinetic measure of metabolic plasticity. Nevertheless, the persistence of NR/R separation under every matched treatment indicated that the resistant state was not erased by either TMZ, simvastatin, or their combination.

**Figure 1.**
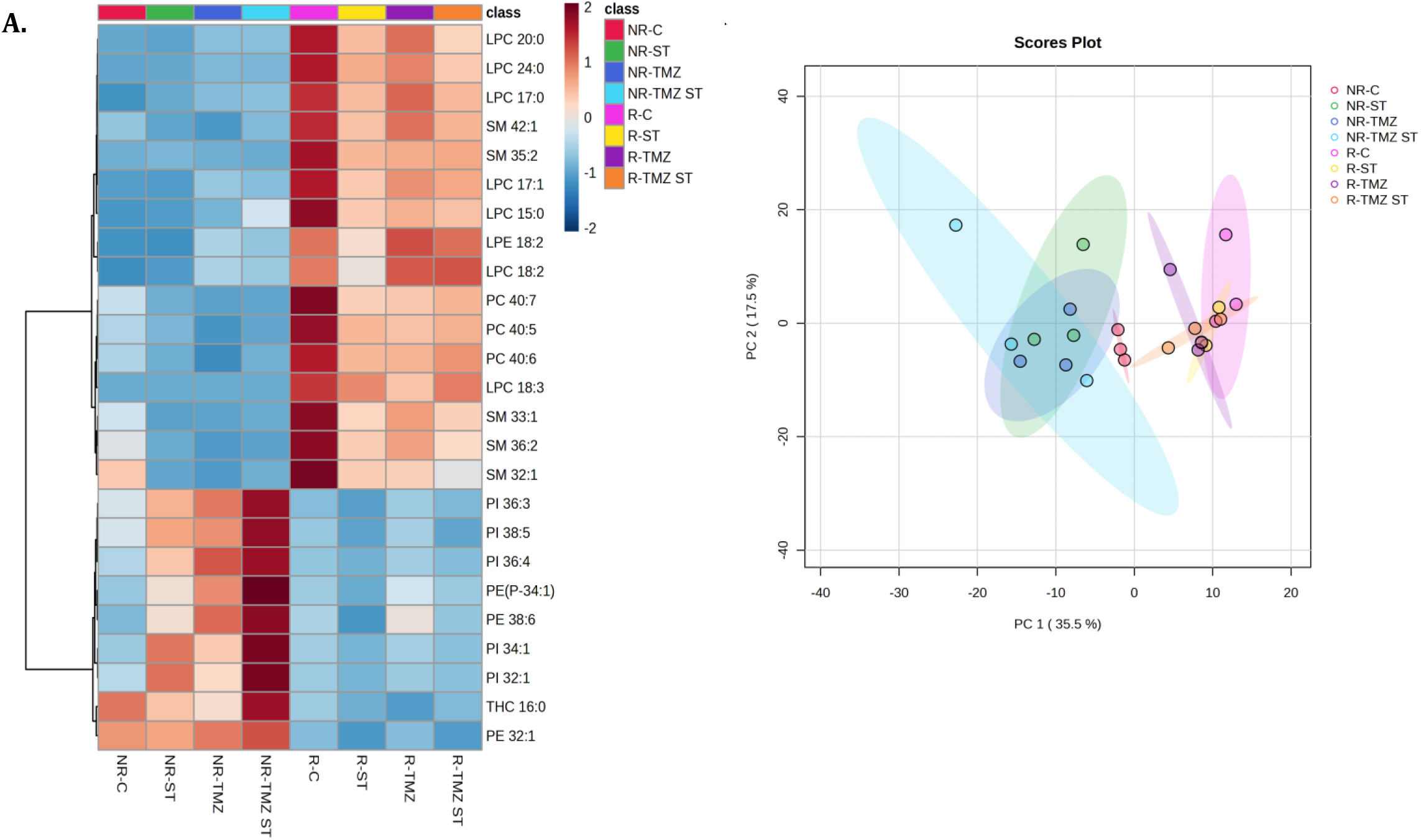
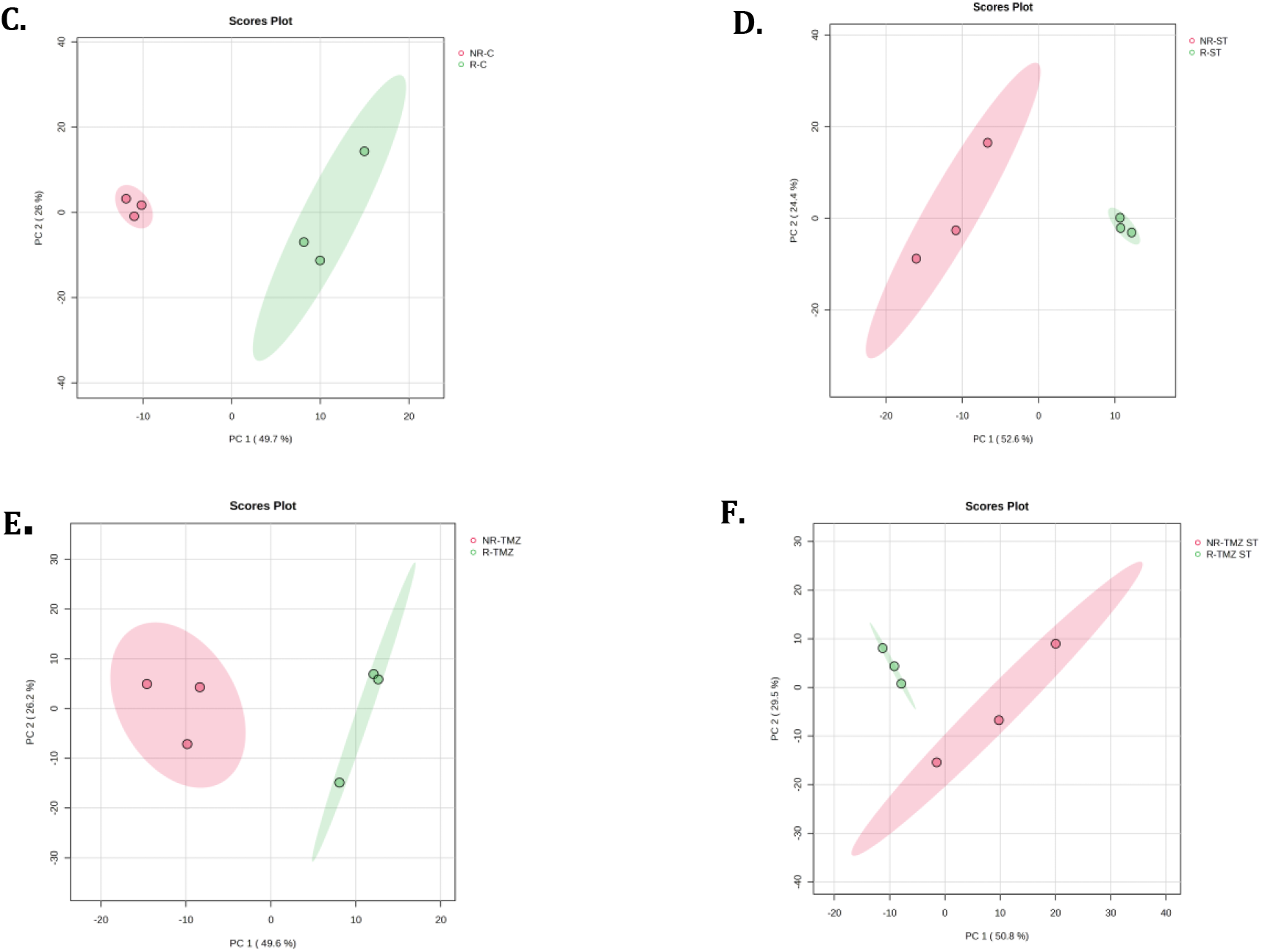
Global lipidomic remodeling separates non-resistant and TMZ-resistant glioblastoma cells across treatment conditions. (A) Hierarchical clustering heatmap of the top 25 changing lipid species across NR-control, NR-ST, NR-TMZ, NR-TMZ-ST and the corresponding resistant conditions. (B) PCA of all eight experimental groups. (C-F) Treatment-stratified PCA for control (C), ST (D), TMZ (E), and TMZ-ST (F), showing persistent NR/R separation within each matched condition. PCA is used as a descriptive representation of global lipid structure; group dispersion should not be interpreted as a direct kinetic measure of lipid flux.

### 3.2 Treatment-stratified differential analysis identifies recurrent lysophospholipid, sphingolipid, and cholesteryl-ester remodeling

Volcano-plot analysis resolved the species-level changes underlying the global separation. At baseline, R cells were enriched in representative lysophospholipid and sphingolipid species, including LPC 24:0 and SM 35:2, whereas several glycerolipid and phospholipid species, including DG 16:0-16:0 and PI 38:2, were lower than in NR cells (**Figure 2A**). This established that acquired resistance was accompanied by a pre-existing lipidomic shift before acute treatment.

**Figure 2.**
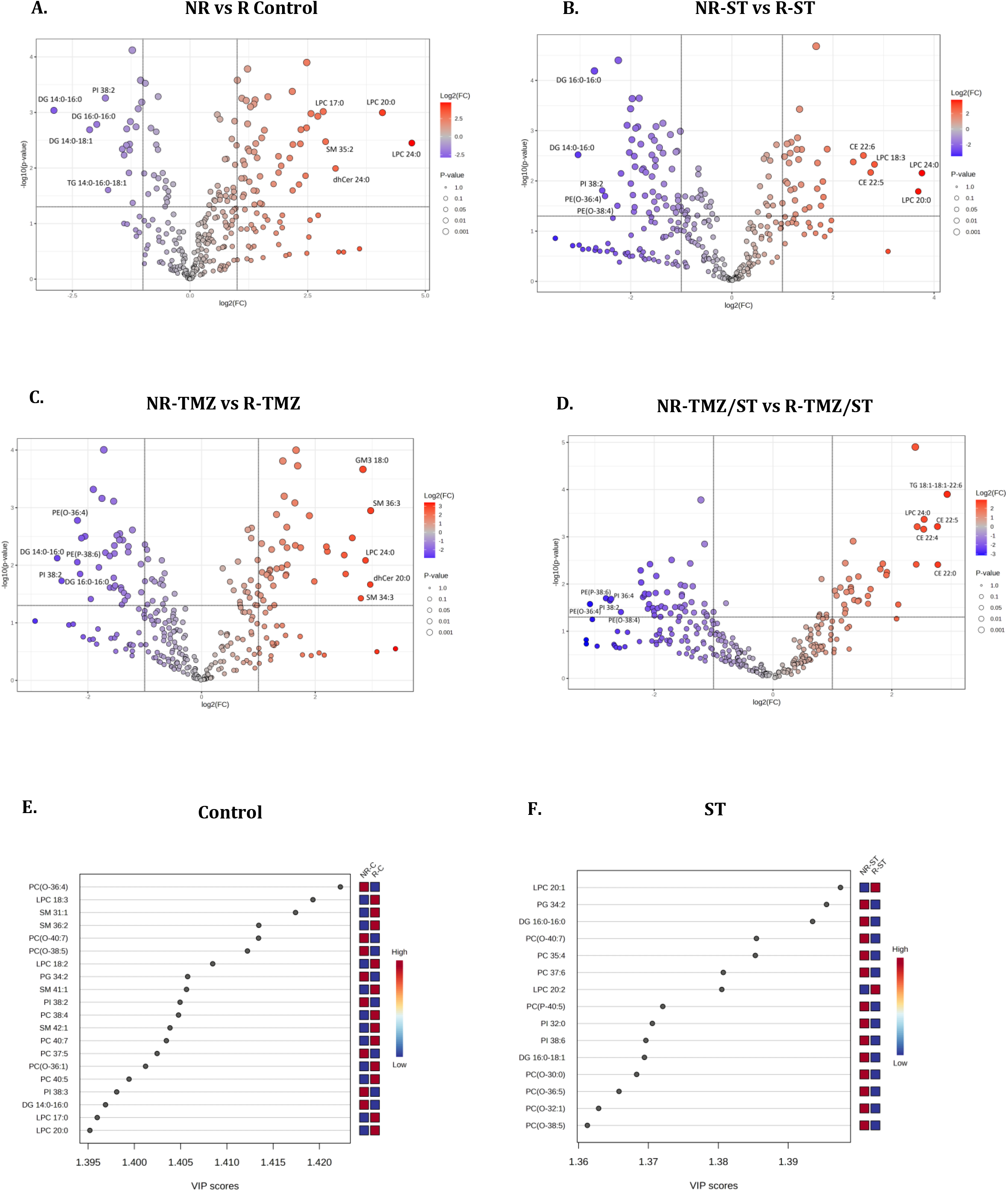
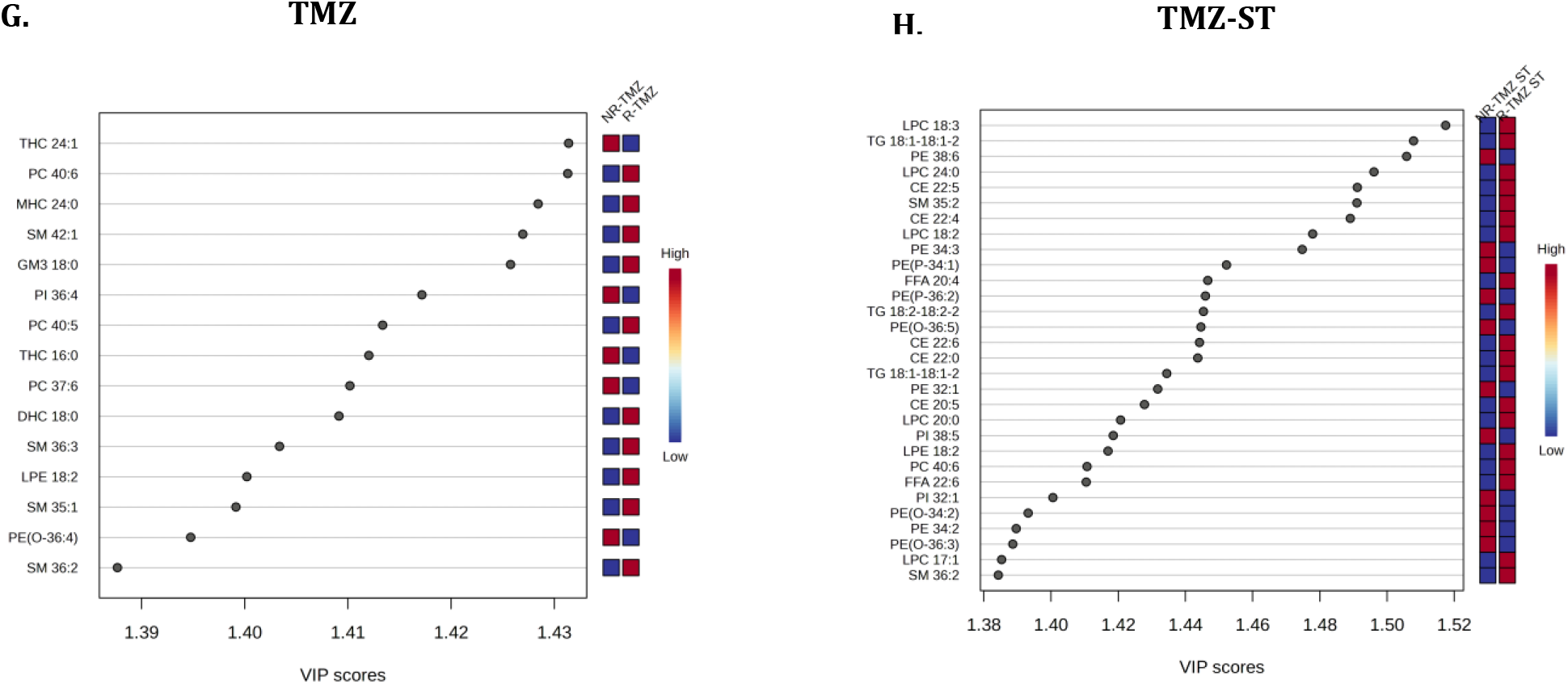
Treatment-stratified species-level lipid differences define recurrent features of the resistant state. Volcano plots compare NR and R cells under control (A), ST (B), TMZ (C), and TMZ-ST (D) conditions; positive log2 fold change denotes relative enrichment in resistant cells. Representative resistance-associated features include LPC and SM species at baseline, CE and LPC species after ST, GM3/SM/LPC species after TMZ, and CE/TG/LPC species after TMZ-ST, whereas selected DG, PI, and PE species are relatively higher in NR cells. (E-H) PLS-DA VIP rankings for the corresponding comparisons using a pre-specified VIP threshold of 1.4. PLS-DA is used for exploratory contributor ranking and not as an estimate of predictive accuracy.

After ST exposure, the resistant state remained associated with higher LPC species and with cholesteryl esters such as CE 22:6 and CE 22:5, while selected DG and PI species remained lower (**Figure 2B**). After TMZ treatment, R cells showed prominent enrichment of GM3 18:0, SM 36:3, LPC 24:0, and related sphingolipid/lysophospholipid species, whereas NR cells retained higher levels of several PE, PI, and DG species (**Figure 2C**). The largest species-level divergence was observed after combined TMZ-ST treatment, where R cells showed increased neutral-lipid storage species, including CE 22:5 and TG 18:1-18:1-22:6, together with LPC 24:0, whereas several PE and PI species were relatively reduced (**Figure 2D**). These findings are consistent with our previous observation that the same combination strongly increases cell death in non-resistant GB cells but does not overcome resistance in the R model [8,9].

Exploratory PLS-DA/VIP analysis provided a complementary ranking of lipids contributing to group separation (**Figure 2E-H**). Baseline contributors included ether/plasmalogen PC, LPC, and SM species; TMZ emphasized sphingolipid features including GM3 and SM; and the combination condition yielded a broader set of CE, triglyceride, lysophospholipid, and sphingomyelin contributors. No ST-only lipid met the pre-specified VIP ≥1.4 threshold despite substantial univariate differences, underscoring the distinction between species-level significance and multivariate contribution. Taken together, the volcano and VIP analyses identify a recurrent resistance-associated signature centered on lysophospholipid/sphingolipid remodeling and treatment-dependent cholesterol storage rather than a single invariant metabolite.

### 3.3 Lipid-family analysis shows that resistance is encoded by coordinated membrane remodeling

We next asked whether the resistance phenotype was better represented at the level of coordinated lipid families. Full-dimensional PERMANOVA across the 19 family-level datasets showed the strongest resistance-associated effects for PI (R²=0.52), PG (R²=0.49), LPC (R²=0.46), LPE (R²=0.45), ether/plasmalogen PC (R²=0.45), and PC (R²=0.41), followed by SM and ether/plasmalogen PE (both R²=0.39) and PE (R²=0.33) (**Figure 3A**; **Table 1**).

**Figure 3.**
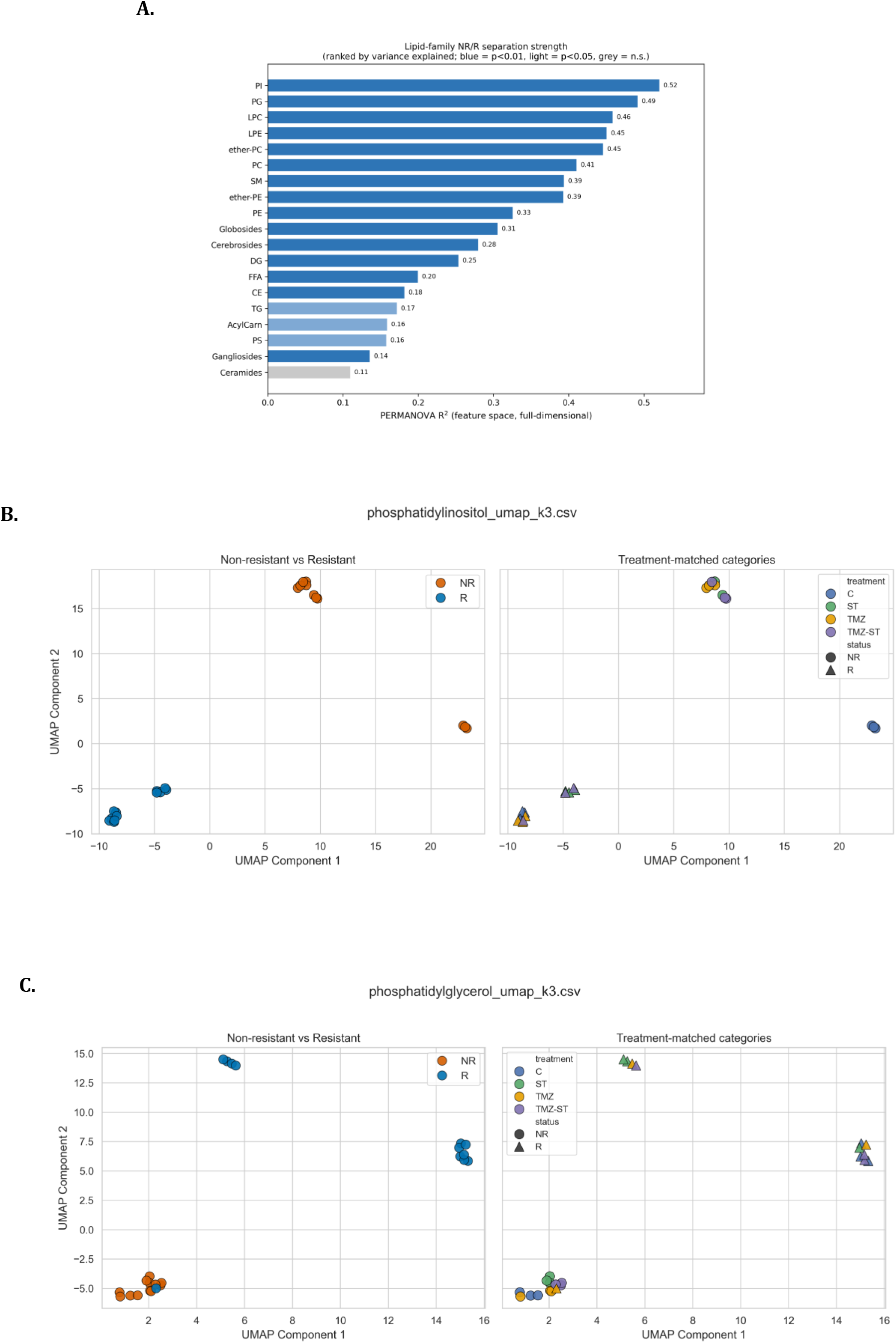
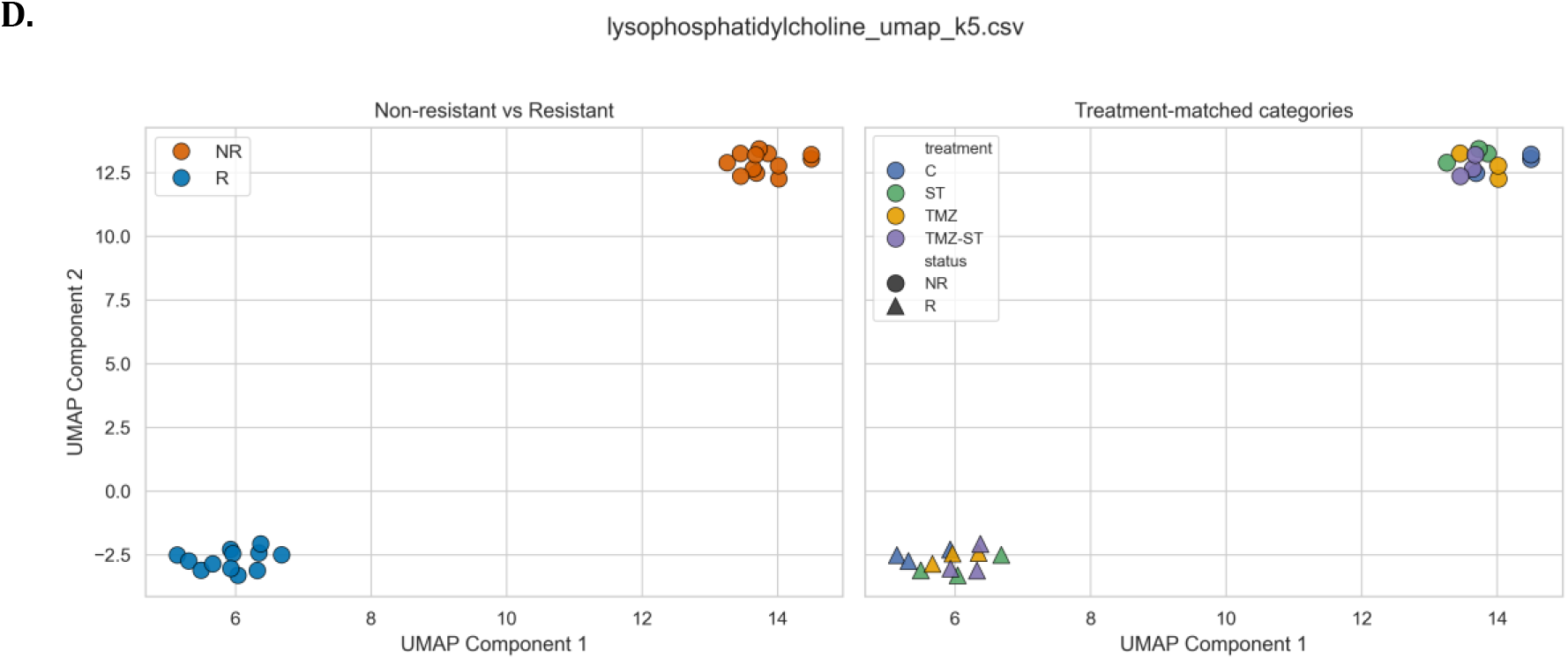
Family-resolved analysis identifies membrane-remodeling lipid families as the dominant features associated with TMZ resistance. (A) Nineteen lipid families ranked by full-dimensional PERMANOVA R² for NR versus R status across control, ST, TMZ, and TMZ-ST conditions. (B-D) Representative UMAP projections for PI (B; R²=0.52), PG (C; R²=0.49), and LPC (D; R²=0.46), shown by resistance status and treatment × status. Full-dimensional PERMANOVA is the primary quantitative output; UMAP and the additional PCA, ISOMAP, and LLE embeddings in Supplementary Figures S1-S4 are visualization/robustness tools.

**Table 1.** Lipid families most strongly associated with TMZ resistance, ranked by full-dimensional PERMANOVA effect size.

| Lipid family | PERMANOVA R<sup>2</sup> | Interpretation |
| --- | --- | --- |
| PI | 0.52 | Strongest resistance-associated family; membrane identity and phosphoinositide/autophagy signaling context |
| PG | 0.49 | Strong mitochondrial/membrane-associated separation |
| LPC | 0.46 | Strong lysophospholipid/membrane-remodeling signature |
| LPE | 0.45 | Parallel lysophospholipid remodeling |
| Ether/plasmalogen PC | 0.45 | Membrane remodeling with potential oxidative-stress relevance |
| PC | 0.41 | Bulk membrane phospholipid remodeling |
| SM | 0.39 | Moderate-to-strong sphingomyelin contribution |
| Ether/plasmalogen PE | 0.39 | Moderate plasmalogen/oxidative-stress-associated contribution |
| PE | 0.33 | Moderate autophagy-membrane-associated separation |

The full-dimensional statistics further showed that the leading families were separated by more than visual embedding alone. For PI, PG, LPC, LPE, ether/plasmalogen PC, and PC, normalized centroid distances ranged from 1.82 to 2.49 and silhouette values from 0.36 to 0.41, while PERMANOVA effect sizes ranged from R² = 0.411 to 0.521 (F = 15.37-23.91; permutation P = 0.0001 for all six families). These full-dimensional statistics are derived from a common distance structure and are reported together as a consistent description of family-level separation.

Representative UMAP projections for PI, PG, and LPC showed clear low-dimensional separation of NR and R samples while retaining treatment labels (**Figure 3B-D**). The remaining UMAP, ISOMAP, LLE, and PCA projections are provided in Supplementary **Figures S1-S4**. Importantly, several families that appeared visually separated in a nonlinear embedding produced more modest full-dimensional effect sizes, reinforcing the decision to use PERMANOVA rather than an embedding as the primary family-ranking criterion.

The univariate class-resolved analysis provided an orthogonal view of the same lipid architecture (**Supplementary Figure S5**). Across treatments, R cells recurrently showed enrichment of LPC/LPE, selected sphingolipid and glycosphingolipid subclasses, and multiple cholesteryl esters, together with depletion or redistribution of PC, PE, PI, PS, PG, and DG pools. The magnitude and direction of individual species were treatment dependent, but the family-level and species-level analyses converged on a broader membrane-remodeling phenotype. Notably, ceramides and gangliosides were not among the strongest global family discriminators despite treatment-specific species changes. This observation is compatible with our previous finding that BCL2L13 modifies ceramide metabolism and autophagy without determining TMZ resistance by itself [11].

#### 3.3.1 Exploratory family-level SVM supports treatment-held-out resistance discrimination

We next asked whether the family-level lipid states that separated NR from R cells could retain discrimination when the pharmacological context itself was withheld. Nine families, phosphatidylcholine, globosides, ether/plasmalogen PE, ether/plasmalogen PC, triglycerides, LPE, PE, LPC, and cerebrosides, achieved MCC = 1.00 under LOOCV, LOCO, and LOTO. In the LOTO analysis, these families also achieved balanced accuracy and ROC-AUC of 1.00, with positive worst-case margin gaps and FDR-adjusted MCC q = 0.0353. Several additional families, including sphingomyelins, ceramides, phosphatidylserine, free fatty acids, cholesteryl esters, and phosphatidylglycerol, retained high but imperfect LOTO performance (MCC approximately 0.92).

Importantly, supervised discrimination did not simply reproduce the PERMANOVA ranking. PI was the strongest family by full-dimensional effect size (R² = 0.521) but showed lower treatment-held-out SVM performance (LOTO MCC = 0.775; balanced accuracy = 0.875), whereas PC, ether/plasmalogen PC/PE, LPC/LPE, and PE combined strong PERMANOVA separation with complete treatment-held-out classification. This difference indicates that effect-size ranking and cross-treatment classification stability capture complementary properties of the resistant lipidome rather than a single interchangeable metric. The analysis therefore supports a distributed family-level resistance signature but, given the small dataset, does not establish a predictive biomarker model.

**Table 2.**
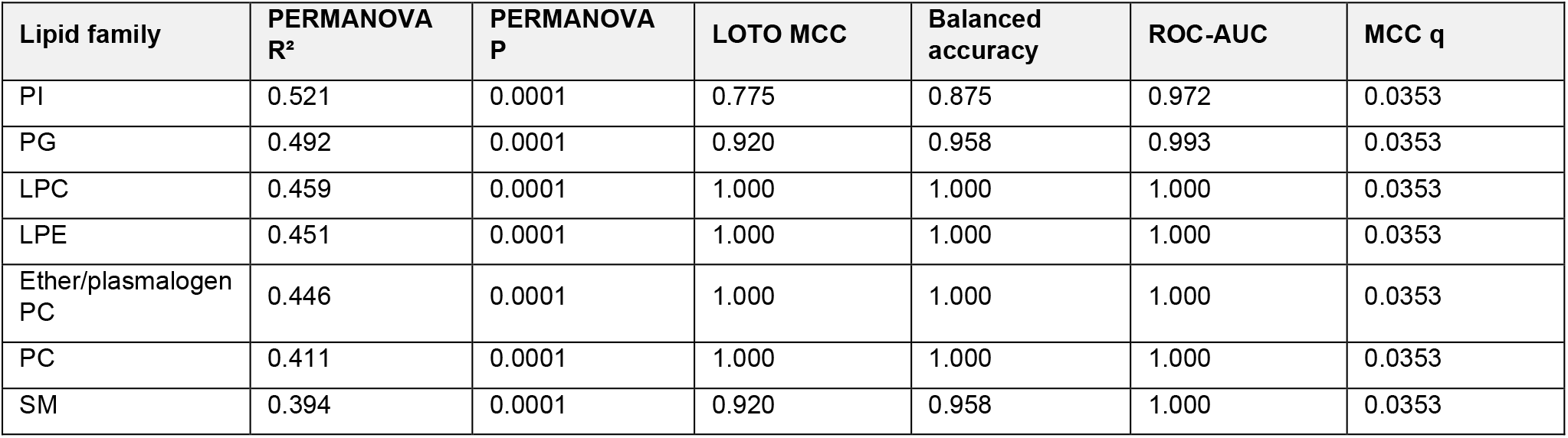
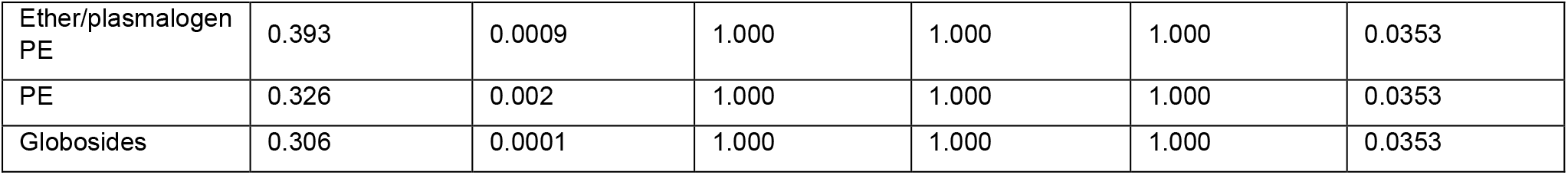
Concordance between full-dimensional family separation and exploratory leave-one-treatment-out SVM discrimination.

Full family-level SVM results across LOOCV, LOCO, and LOTO are provided in Supplementary **Table S2**. Reduced-signature retention analyses are provided in Supplementary **Table S3**.

### 3.4 Structure-informed pathway inference prioritizes membrane-trafficking and survival networks

To generate mechanistic hypotheses from the discriminant lipid structures, we mapped canonical SMILES representations to predicted human protein targets and then performed STRING/KEGG enrichment. In untreated R versus NR cells, predicted target sets were enriched for pathways including Rap1, Ras, PI3K-Akt, phospholipase D, focal adhesion, MAPK-related signaling, and central carbon metabolism (**Figure 4A**). Under TMZ, the predicted associations additionally emphasized neuroactive ligand-receptor interaction, calcium signaling, and lipid-metabolic pathways including arachidonic-, linoleic-, alpha-linolenic-, and ether-lipid metabolism (**Figure 4B**). Under TMZ-ST, PPAR signaling, PI3K-Akt, insulin-resistance-related pathways, arachidonic acid metabolism, and inflammatory mediator regulation of TRP channels were among the prominent enriched terms (**Figure 4C**).

**Figure 4.**
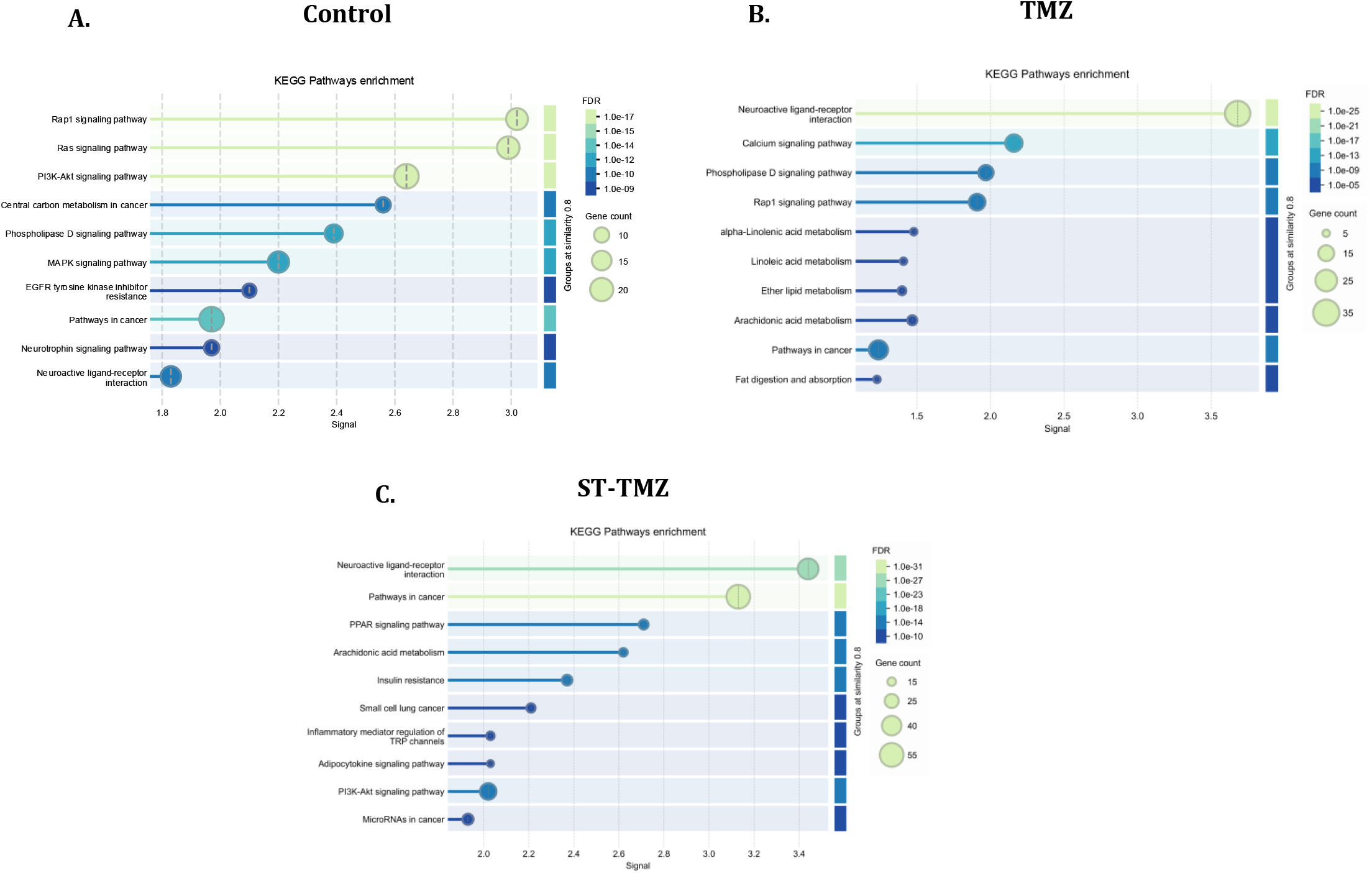
Structure-informed pathway inference from discriminant lipids prioritizes membrane-trafficking, survival, and lipid-metabolic pathways. Canonical SMILES strings for high-contribution lipids were mapped to predicted human protein targets and subjected to STRING/KEGG enrichment. Panels show untreated NR versus R (A), TMZ-treated NR versus R (B), and TMZ-ST-treated NR versus R (C) comparisons. Recurrent enriched terms include Rap1, Ras, PI3K-Akt, phospholipase D, calcium, PPAR, and lipid-metabolic pathways. These results are hypothesis generating and do not establish direct lipid-protein binding or pathway activation.

These enrichments were not interpreted as direct measurements of pathway activation. Instead, they identify candidate signaling contexts through which the observed lipid architecture could intersect with vesicle trafficking, membrane homeostasis, survival signaling, and stress adaptation. Their biological relevance was therefore considered together with the lipid-family results and the ultrastructural phenotype rather than used as stand-alone mechanistic proof.

### 3.5 Resistant cells maintain a vesicle-rich ultrastructure across treatment states

TEM was used to determine whether the resistant lipid state was accompanied by persistent changes in intracellular membrane architecture. Untreated R cells contained more abundant membrane-bound and double-membrane autophagosome-like structures than NR controls, together with enlarged vacuolar compartments and irregular mitochondrial morphology (**Figure 5A**). These features are consistent with the previously established accumulation of autophagic structures in the TMZ-resistant model [9–11].

**Figure 5.**
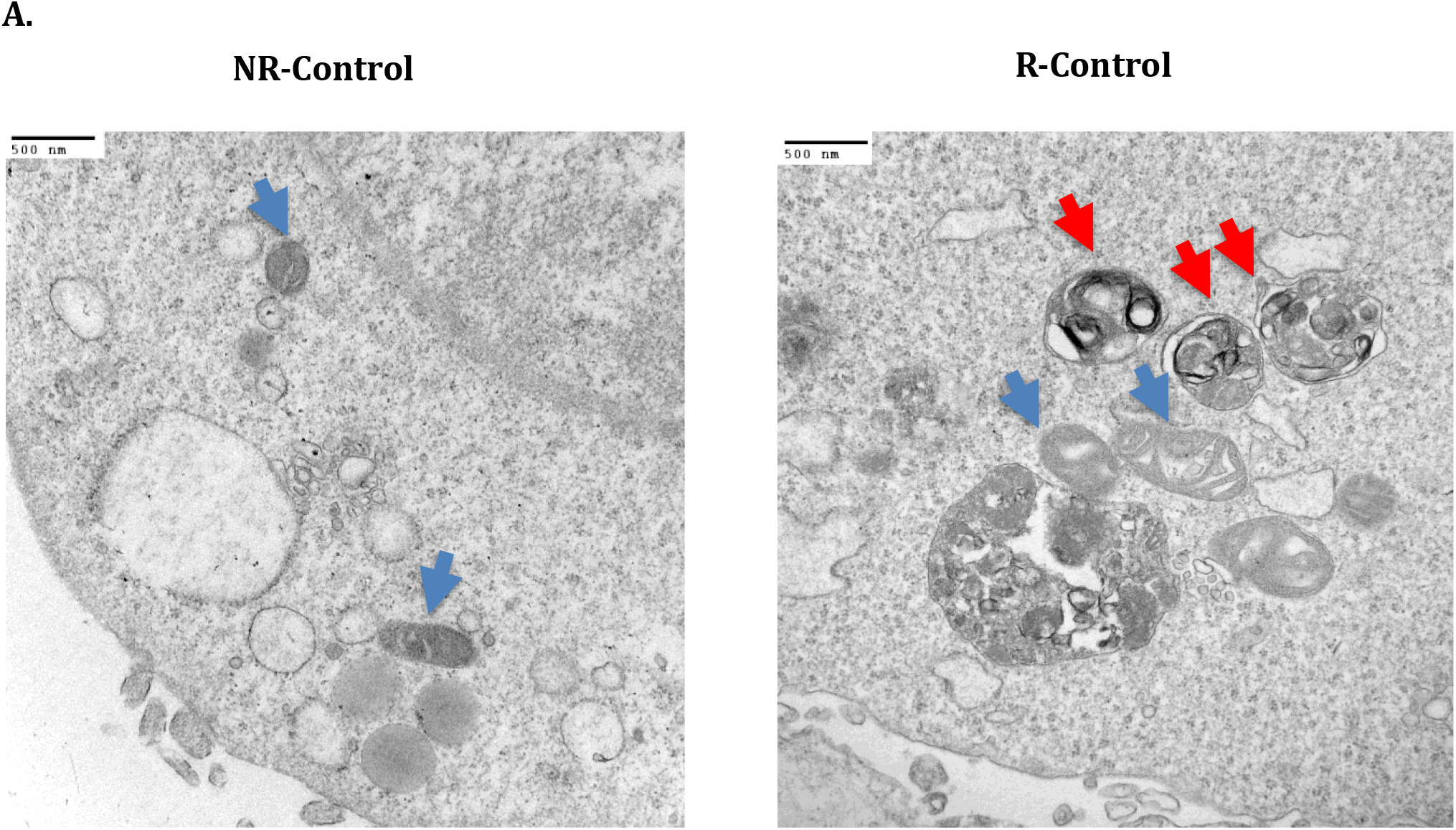
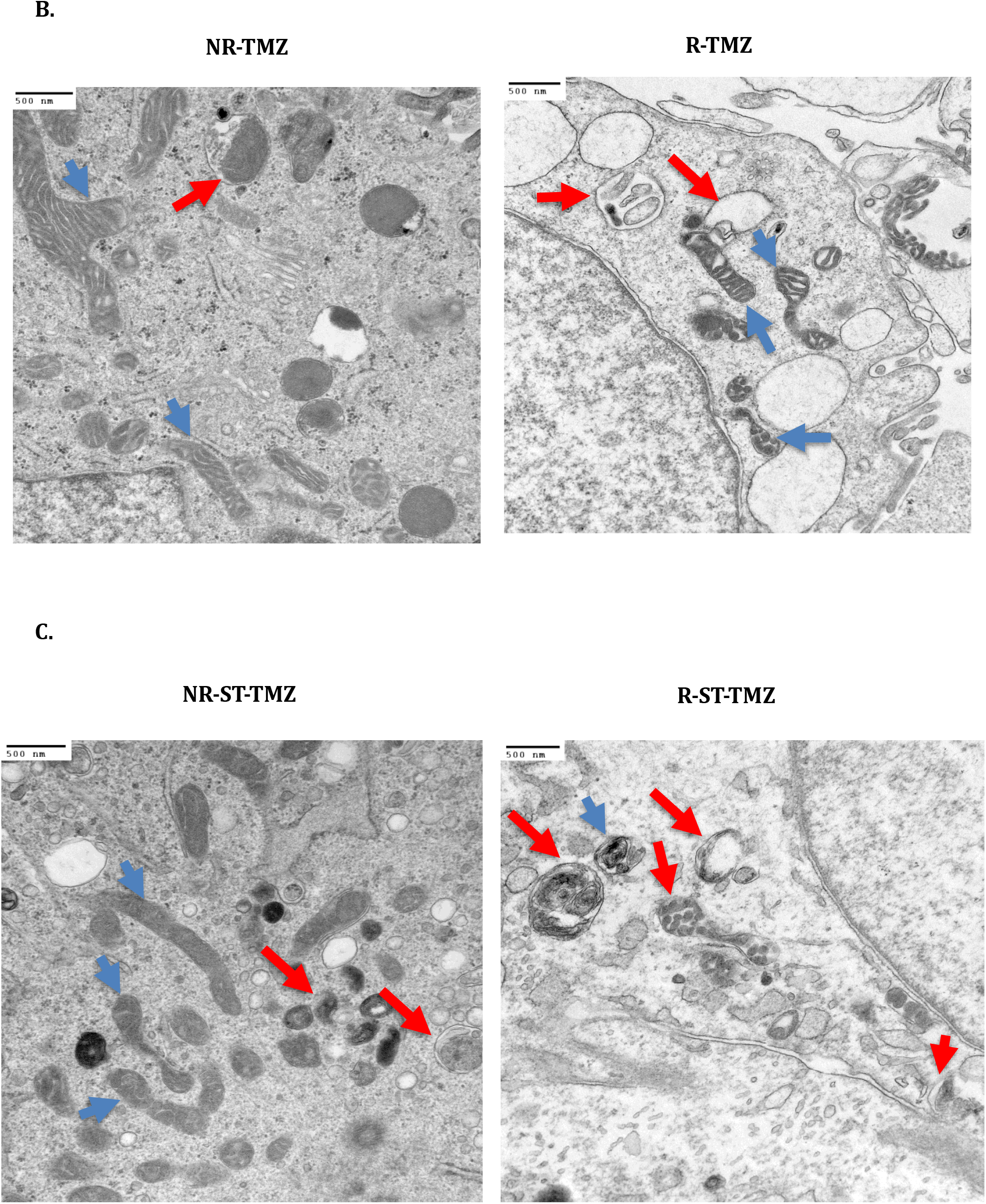
Ultrastructural membrane remodeling in non-resistant and TMZ-resistant glioblastoma cells. TEM was used to compare NR and R cells under control (A), TMZ (B), and TMZ-ST (C) conditions. Untreated R cells show abundant membrane-bound/autophagosome-like vesicles relative to NR controls. TMZ increases vesicular structures in NR cells, whereas R cells retain a dense vesicle-rich architecture. TMZ-ST produces pronounced vesicle accumulation in NR cells, consistent with the previously established late-stage autophagy inhibition induced by this combination, while R cells retain a vesicle-rich phenotype resembling their baseline state. Red arrows indicate vesicular/autophagosome-like structures; blue arrows indicate mitochondria. TEM provides morphological context and is not, by itself, a measure of autophagic flux. Exploratory basal-condition AP/AL morphometry is summarized in **Table 3**.

Exploratory morphometry of the untreated TEM dataset provided a directional quantitative complement to the qualitative ultrastructure. The annotated NR images contained 31 autophagosomes and 12 autolysosomes (43 structures total), whereas R images contained 52 autophagosomes and 8 autolysosomes (60 structures total). The AP fraction was therefore higher in R than NR cells (0.882 versus 0.676; difference +0.206; Hedges’ g = 1.36; P = 0.10; q = 0.20). Exact permutation over all 20 replicate-label assignments has a minimum stainable two-sided P of 0.10, which this comparison achieved. Median AP area was modestly lower in R cells (0.444 versus 0.543 μm²; -18%; g = -0.44; q = 0.60), whereas median AL area was directionally larger (0.852 versus 0.516 μm²; +65%; g = 1.17; q = 0.267). These results support a directional shift toward autophagosome predominance and altered vesicle size distribution, but they are not confirmatory because the structure counts are nested observations and the statistical comparisons are underpowered.

**Table 3.** Exploratory quantitative TEM morphometry in untreated NR and R cells.

| Measure | NR | R | Difference | Hedges' g | P | q |
| --- | --- | --- | --- | --- | --- | --- |
| AP fraction | 0.676 | 0.882 | +0.206 | 1.36 | 0.10 | 0.20 |
| Median AP area ($\mu\text{m}^2$) | 0.543 | 0.444 | -18% | -0.44 | 0.60 | 0.60 |
| Median AL area ($\mu\text{m}^2$) | 0.516 | 0.852 | +65% | 1.17 | 0.20 | 0.267 |

AP, autophagosome; AL, autolysosome. AP fraction = $\text{AP}/(\text{AP} + \text{AL})$ . Values are exploratory morphometric summaries from the available annotated basal TEM dataset and should not be interpreted as biological-replicate-level confirmatory statistics.

AP, autophagosome; AL, autolysosome. AP fraction = AP/(AP + AL). Values are exploratory morphometric summaries from the available annotated basal TEM dataset and should not be interpreted as biological-replicate-level confirmatory statistics.

After TMZ treatment, NR cells developed additional vesicular/vacuolar structures consistent with an acute stress response, whereas R cells retained a dense population of autophagosome-like and multilamellar compartments (**Figure 5B**). Combined TMZ-ST treatment produced marked vesicle accumulation in NR cells, compatible with the late-stage autophagy inhibition previously demonstrated for this drug combination [8,24]. In contrast, R cells again retained a vesicle-rich architecture closely resembling their baseline state (**Figure 5C**).

TEM alone cannot measure flux; however, when interpreted in the context of our prior biochemical flux studies, the persistent vesicular phenotype provides ultrastructural support for a model in which resistant cells enter treatment with a pre-existing late-stage autophagy defect and a remodeled membrane system. The relative stability of this architecture across treatments parallels the persistence of the resistance-associated lipidome.

## 4. Discussion

The central finding of this study is that acquired TMZ resistance in U251 GB cells is associated with a broad, treatment-resilient reorganization of the lipidome. This state is evident before acute treatment and remains distinguishable during TMZ, simvastatin, and TMZ-ST exposure. The most informative signal is not a single lipid species. Rather, resistance is represented by coordinated changes across membrane-associated families, particularly PI, PG, LPC, LPE, ether/plasmalogen PC, and PC together with treatment-dependent accumulation of sphingolipids and cholesteryl esters. This distinction matters because it changes the biological interpretation from a search for one resistance metabolite to a model of membrane-system adaptation.

The study was motivated directly by a sequence of observations from our group. We previously showed that simvastatin sensitizes non-resistant GB cells to TMZ by blocking autophagosome-lysosome fusion and increasing cell death [8]. We subsequently demonstrated that the combination engages the unfolded protein response while autophagy flux is inhibited [24,25]. In the resistant U251 model, however, autophagy flux is already impaired, and the cells are refractory to both TMZ and statin-mediated sensitization [9–11]. The autophagy-cholesterol study further showed that resistant cells can accumulate selected cholesteryl esters despite suppression of de novo cholesterol synthesis and that simvastatin changes the cholesterol ester profile without restoring TMZ sensitivity [9]. The current data place those cholesterol findings within a larger lipidomic framework: the resistant phenotype includes cholesterol storage, but the strongest family-level separation is carried by PI, PG, lysophospholipid, and phosphatidylcholine-related families rather than by cholesteryl esters alone.

This broader view also helps reconcile our BCL2L13/ceramide observations. BCL2L13 altered autophagy-associated processing and ceramide composition in resistant and non-resistant GB cells, yet BCL2L13 depletion did not restore TMZ sensitivity [11]. In the present analysis, ceramide and ganglioside families show treatment-specific changes but do not dominate the full-dimensional NR/R separation. Together, the two studies argue that sphingolipid remodeling is part of the resistant state but is unlikely to be sufficient as a single causal axis. The resistance phenotype instead appears to emerge from coordinated remodeling across several membrane and storage-lipid systems.

The exploratory SVM analysis addresses the same contrast through a supervised rather than distance-based criterion. Several families that were prominent by PERMANOVA, particularly PC, ether/plasmalogen PC, LPC, LPE, ether/plasmalogen PE, and PE also retained complete separation when an entire treatment was withheld. At the same time, the ranking was not identical: PI produced the largest PERMANOVA effect, but its classification degraded when an entire treatment was withheld ((LOOCV MCC = 1.00; LOTO MCC = 0.775), while triglycerides and glycosphingolipid families showed stronger treatment-held-out classification than their PERMANOVA rank alone would predict. This is biologically and analytically informative because PERMANOVA quantifies overall multivariate displacement, whereas SVM performance asks whether a decision boundary learned in other treatment contexts remains useful in a withheld treatment. Agreement between the two criterion indicates that the family-level separation is not specific to a distance-based formulation; discordance cautions against reducing the phenotype to a single ranked list.

Feature-retention analyses further suggested that some family-level signals are internally redundant: the full LOTO performance could be retained with small subsets in several families, including PC, LPC, ether/plasmalogen PC/PE, LPE, PE, and triglycerides. However, these subsets were selected and tested within the same 24-sample dataset and therefore may be optimistic. We accordingly treat them as evidence that the family signal is not necessarily dependent on every measured species, not as a validated reduced biomarker panel.

The strong PI-family effect is particularly relevant to autophagy and vesicle regulation. Phosphatidylinositol-derived lipids organize membrane identity and recruit autophagy machinery, while phosphatidylethanolamine is required for LC3/ATG8 lipidation and autophagosomal membrane association [12–16,23]. In NR cells, combination treatment was accompanied by conspicuous PI/PE remodeling and extensive vesicle accumulation, consistent with an acute membrane response to the autophagosome-lysosome fusion defect induced by simvastatin-TMZ [8,24]. Resistant cells, by contrast, entered treatment with a distinct lipid composition and vesicle-rich architecture and did not converge toward the NR treatment state. We therefore interpret the reduced treatment-associated displacement of the R lipidome as evidence of a pre-established adaptive state, not as proof that individual lipid pools are kinetically static. Direct flux measurements with stable-isotope tracers will be required to distinguish synthesis, uptake, remodeling, and degradation.

PG, LPC, LPE, PC, and ether/plasmalogen lipids add complementary biological dimensions. PG is closely related to mitochondrial membrane metabolism and therefore may reflect the mitochondrial remodeling previously observed in therapy-tolerant GB cells [9]. Recurrent LPC/LPE enrichment is consistent with altered phospholipid turnover and membrane remodeling, although the present data do not establish whether this arises through phospholipase activity, reacylation pathways, uptake, or impaired degradation. Ether/plasmalogen lipids are particularly relevant to oxidative and lipid-peroxidation biology; their remodeling could influence susceptibility to ferroptotic stress, a pathway implicated in TMZ response and resistance [40–42]. These interpretations are mechanistic hypotheses generated from abundance patterns and should be tested through enzyme-level perturbation and isotope-resolved lipid flux.

The cholesteryl-ester phenotype remains an important component of the resistant state. GB cells can rely on cholesterol uptake, esterification, storage, and lipophagy to maintain cholesterol homeostasis, and SOAT1/ACAT1-dependent esterification has been linked to glioma growth [29–33]. Our prior work showed the apparently paradoxical combination of reduced cholesterol biosynthetic signaling with accumulation of selected long-chain cholesteryl esters in resistant cells [9]. The present treatment-stratified data extend that finding by showing that CE enrichment is especially evident under ST-containing conditions and remains prominent during TMZ-ST exposure. Because static lipid abundance cannot distinguish altered synthesis from uptake, esterification, hydrolysis, or export, the most defensible conclusion is that resistant cells preserve a cholesterol-storage phenotype despite mevalonate-pathway inhibition. This may help explain why simply reducing endogenous cholesterol synthesis with simvastatin does not recreate the sensitizing effect observed in NR cells.

The structure-informed pathway analysis provides a complementary, but explicitly exploratory, layer. Predicted target sets linked discriminant lipids to Rap1, PI3K-Akt, phospholipase D, calcium, PPAR, and lipid-metabolic pathways. Rap1 can coordinate lysosomal organization with nutrient signaling, and phospholipase D participates in autophagy regulation and membrane trafficking [34,35]. Lysosomal calcium-calcineurin-TFEB signaling provides another plausible connection between vesicle abundance, lysosome adaptation, and autophagy control [36]. Arachidonic acid and lysophosphatidic-acid signaling are also relevant to GB survival and motility [37–39]. However, SwissTargetPrediction was developed for small-molecule target inference, and predictions from complex endogenous lipids should not be equated with direct binding or pathway activation. These results therefore serve as a prioritization map for experiments rather than mechanistic validation.

The TEM observations reinforce the membrane-centered interpretation while also defining its limits. Resistant cells contained persistent autophagosome-like and vacuolar structures at baseline and after treatment. In NR cells, TMZ and particularly TMZ-ST produced acute vesicular remodeling. These observations are consistent with the biochemical flux phenotypes established in our prior studies [8–11,24], but ultrastructure alone cannot distinguish increased autophagosome formation from impaired clearance. Cholesterol accumulation itself can impair lysosomal fusion and SNARE function in other contexts [26,27], and autophagy participates directly in lipid mobilization through lipophagy [28]. This provides a plausible framework in which altered lipid storage and impaired lysosomal turnover reinforce one another, but the direction of causality remains to be established in GB.

The exploratory TEM morphometry adds a quantitative but deliberately limited layer to this interpretation. R cells showed a higher AP fraction than NR cells (0.882 versus 0.676; Hedges’ g = 1.36), and a higher AP density (3.854 versus 2.251 per 100 μm²; g = 1.75), and a directionally larger median AL area (+65%; g = 1.17), whereas median AP area was slightly smaller. No comparison reached q < 0.05, although the minimum attainable two-sided P under exact permutation at n = 3 per state is 0.10, which the AP fraction and AP density comparisons achieved. Inference was performed at the biological-replicate level (n = 3 per state), which limits power. Thus, the morphometry should not be used as stand-alone evidence of flux blockade. Its value is as directional support for the previously established biochemical phenotype: resistant cells contain proportionally more autophagosome-like structures and an altered vesicle-size landscape at baseline.

Several limitations are important. First, the study uses one established GB cell model and three biological replicates per condition; the high-dimensional analyses therefore require cautious interpretation and independent replication. Second, the lipidomics platform is targeted and reports abundance rather than metabolic flux, so pathway directionality cannot be inferred from concentration alone. Third, PLS-DA was used only for exploratory feature prioritization and, with the present sample size, should not be interpreted as a validated classifier. Fourth, the structure-to-target/KEGG workflow is hypothesis generating and requires biochemical validation. Fifth, TEM is qualitative with respect to flux and does not quantify autophagosome-lysosome fusion. Finally, we did not functionally perturb the lipid families or enzymes identified by PERMANOVA. Clinical attempts to inhibit autophagy alongside TMZ have demonstrated feasibility but also the difficulty of achieving selective, tolerable target engagement [43]. These limitations define the next experimental steps: isotope tracing, direct measurements of lipid uptake and esterification, targeted perturbation of SOAT1/ACAT1 and phospholipid-remodeling enzymes, quantitative lysosomal-fusion assays, and validation in patient-derived GB models [30,33,44,45]. The new SVM and reduced-signature analyses remain internal to the same 24-sample experiment; the high number of families with near-perfect classification, together with extensive enumeration of the 70 possible condition-level label assignments, which sets a minimum attainable P of 0.014, requires external validation before any predictive interpretation. Likewise, the TEM morphometric statistics are exploratory because inference rests on three biological replicates per state, for which the minimum attainable two-sided P under exact permutation is 0.10.

Despite these limitations, the study provides a coherent extension of our prior work. The sensitizing effect of simvastatin-TMZ in non-resistant GB cells is associated with acute autophagy disruption and substantial lipid remodeling, whereas acquired TMZ resistance is accompanied by a pre-existing autophagy defect and a lipidomic state that remains distinct across the same perturbations. The strongest statistical signal lies in coordinated membrane-lipid families, while cholesteryl esters provide a prominent storage phenotype and sphingolipids contribute treatment-specific features. This treatment-resilient lipid architecture offers a mechanistic explanation for why a statin strategy that is effective in a sensitive state can fail after resistance is established and identifies lipid buffering, membrane remodeling, and vesicle homeostasis as testable vulnerabilities.

## 5. Conclusions

Acquired TMZ resistance in U251 glioblastoma cells is associated with a stable, treatment-resilient lipidomic architecture that persists during TMZ, simvastatin, and combined TMZ-ST exposure. The resistant state is defined most strongly at the family level by coordinated remodeling of PI, PG, LPC, LPE, ether/plasmalogen PC, and PC, with recurrent treatment-dependent contributions from sphingolipids and cholesteryl esters. In the context of our previous demonstration of late-stage autophagy blockade and failure of statin-mediated resensitization, the present findings support a model in which chemoresistance is accompanied by broad membrane and lipid-storage adaptation rather than dependence on a single lipid species. Future work should test whether disrupting cholesterol esterification, lysophospholipid turnover, phosphoinositide remodeling, or vesicle-trafficking pathways can dismantle this resistant state and restore therapeutic vulnerability. Family-level SVM analysis supports this conclusion by showing that several membrane and storage-lipid families retain NR/R discrimination even when an entire treatment is withheld, while quantitative TEM morphometry provides directional support for a baseline shift toward autophagosome predominance. Both analyses remain proof-of-concept and require independent biological validation.

## Acknowledgment

The authors acknowledge the use of ChatGPT (OpenAI) under direct author supervision for English-language editing, manuscript organization, and refinement of scientific presentation. The tool was not used to generate experimental data or to independently determine scientific conclusions. All analyses, interpretations, references, figures, and final wording were reviewed and approved by the authors, who take full responsibility for the manuscript.

## Author Contributions

Kianoosh Naghibzadeh and Rui Vitorino performed lipidomics analyses and contributed to data interpretation and preparation of lipidomics-related results. Hyunjin Cho performed machine learning. Amir Barzegar Behrooz, Marco Cordani, and Carla Vitorino contributed glioblastoma-related expertise and participated in data interpretation and manuscript writing. Marek J. Los provided expertise in cell death mechanisms and contributed to interpretation of results. Mahboubeh Kavoosi performed univariate statistical analyses and prepared associated graphical representations. Stevan Pecic contributed expertise in lipid biology and data interpretation. Amir Ravandi contributed to lipidomics analysis, including access to and support from the lipidomics core facility, and assisted in data interpretation. Shahla Shojaei performed all experimental work. Saeid Ghavami conceived and designed the study, supervised the project, and led data interpretation and conceptualization of the results, as well as manuscript preparation.

