## Supplemental Table Figures for "Treatment-Resilient Lipid Remodeling Defines Temozolomide Resistance and Failure of Simvastatin Sensitization in Glioblastoma"

#### Supplementary Figure S1

**A.**

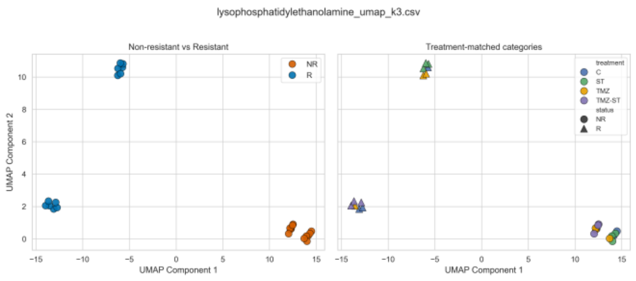


**B.**

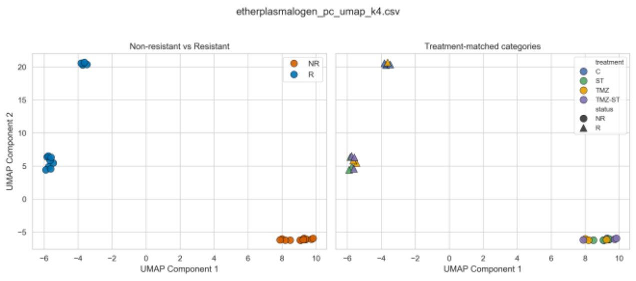


**C.**

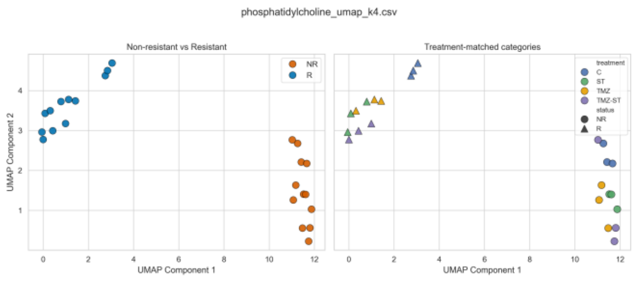


**D.**

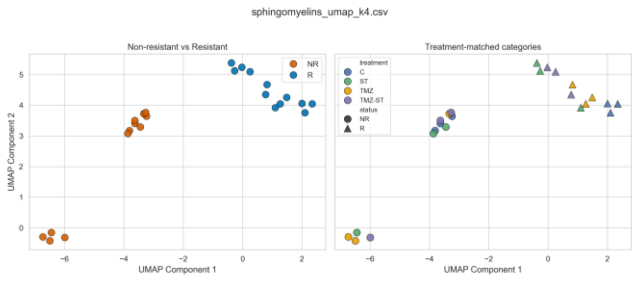


**E.**

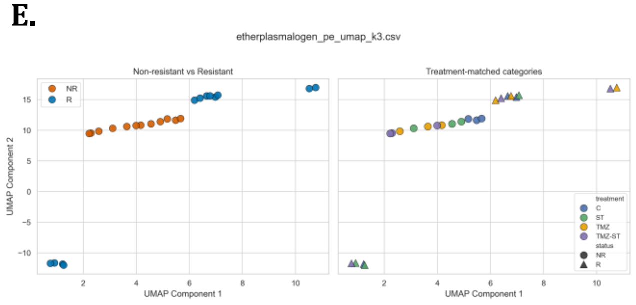


**F.**

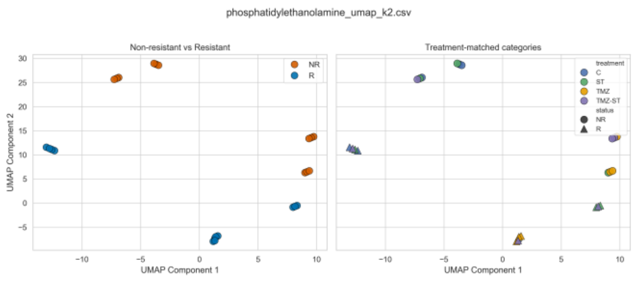


**G.**

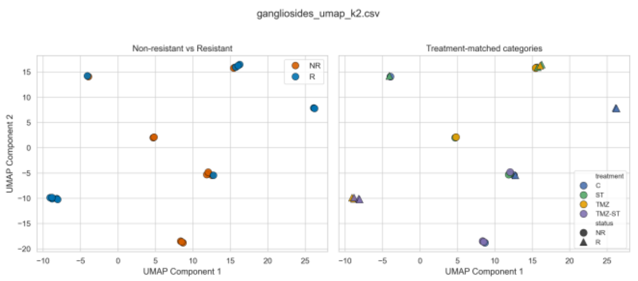


**H.**

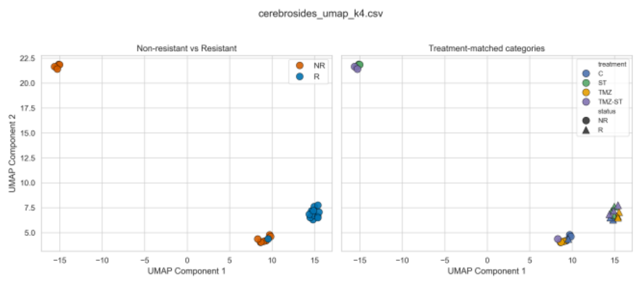

**I.**

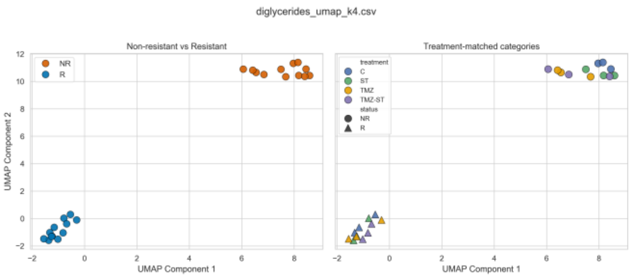


**J.**

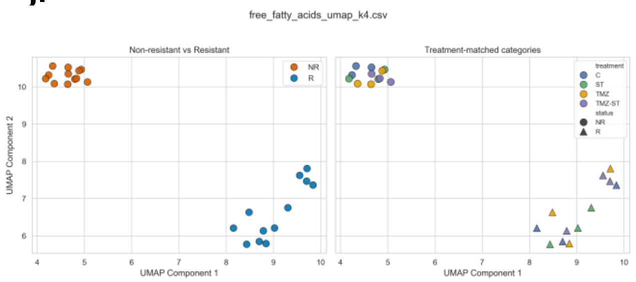


**K.**

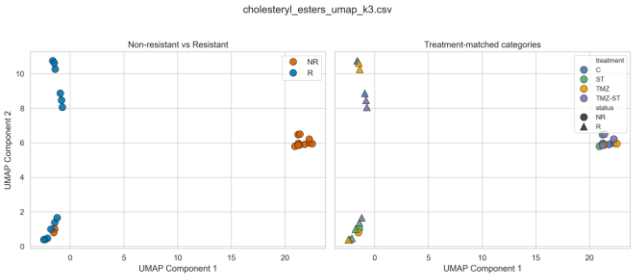


**L.**

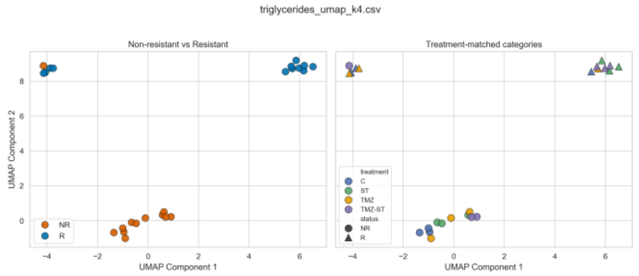


**M.**

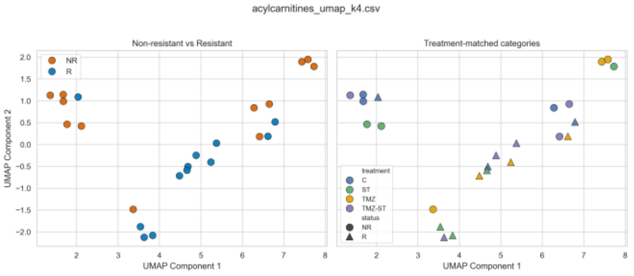


**N.**

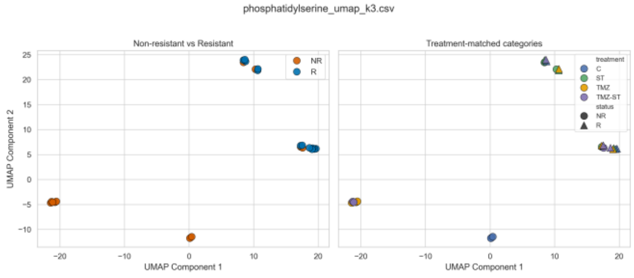


**O.**

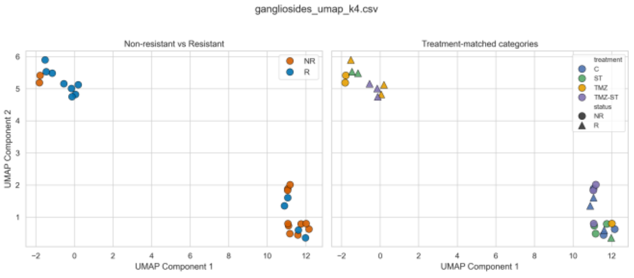


**P.**

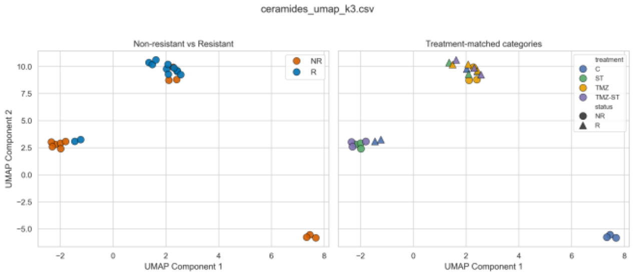


**Supplementary Figure S1**. UMAP embeddings of lipid families not shown in main Figure 3. Two-dimensional UMAP projections of log2-transformed, z-score-normalized lipid intensities compare non-resistant (NR) and resistant (R) U251 glioblastoma cells across control, ST, TMZ, and TMZ-ST conditions. For each family, the left plot is colored by resistance status and the right plot by treatment × status. UMAP is provided for visualization/robustness assessment; full-dimensional PERMANOVA is the primary family-level statistic.

### Supplementary Figure S2

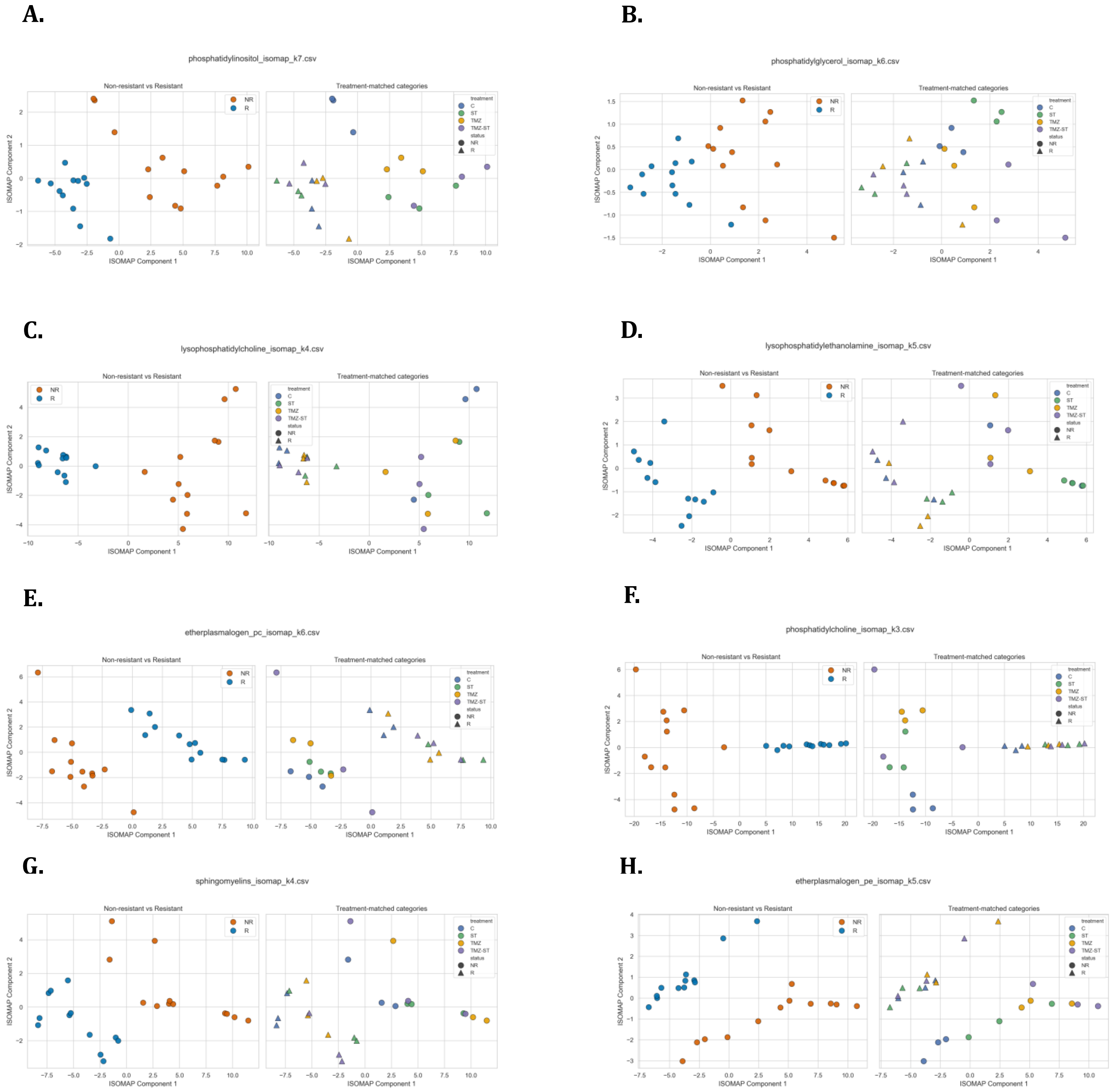

**I.**

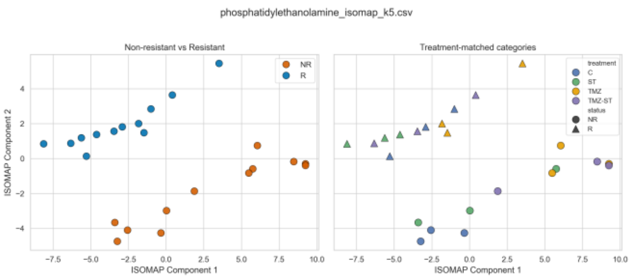
phosphatidylethanolamine_isomap_k5.csv
Non-resistant vs Resistant
Treatment-matched categories
ISOMAP Component 1
ISOMAP Component 2
NR
R
treatment
C
ST
TMZ
TMZ-ST
status
NR
R

**J.**

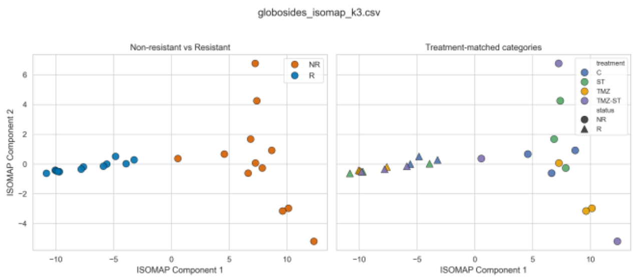
globosides_isomap_k3.csv
Non-resistant vs Resistant
Treatment-matched categories
ISOMAP Component 1
ISOMAP Component 2
NR
R
treatment
C
ST
TMZ
TMZ-ST
status
NR
R

**K.**

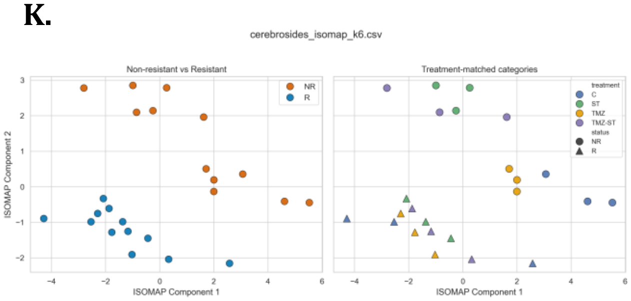
cerebrosides_isomap_k6.csv
Non-resistant vs Resistant
Treatment-matched categories
ISOMAP Component 1
ISOMAP Component 2
NR
R
treatment
C
ST
TMZ
TMZ-ST
status
NR
R

**L.**

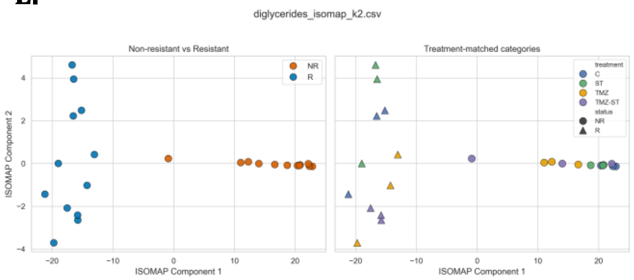
diglycerides_isomap_k2.csv
Non-resistant vs Resistant
Treatment-matched categories
ISOMAP Component 1
ISOMAP Component 2
NR
R
treatment
C
ST
TMZ
TMZ-ST
status
NR
R

**M.**

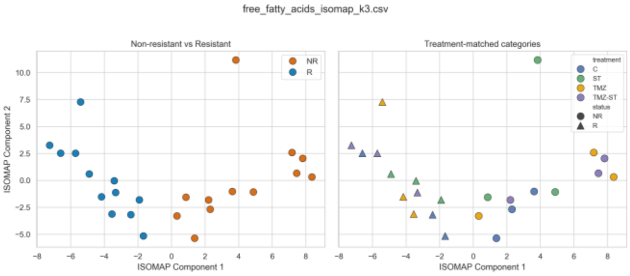
free_fatty_acids_isomap_k3.csv
Non-resistant vs Resistant
Treatment-matched categories
ISOMAP Component 1
ISOMAP Component 2
NR
R
treatment
C
ST
TMZ
TMZ-ST
status
NR
R

**N.**

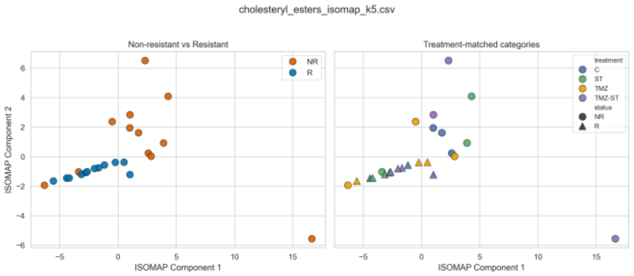
cholesteryl_esters_isomap_k5.csv
Non-resistant vs Resistant
Treatment-matched categories
ISOMAP Component 1
ISOMAP Component 2
NR
R
treatment
C
ST
TMZ
TMZ-ST
status
NR
R

**O.**

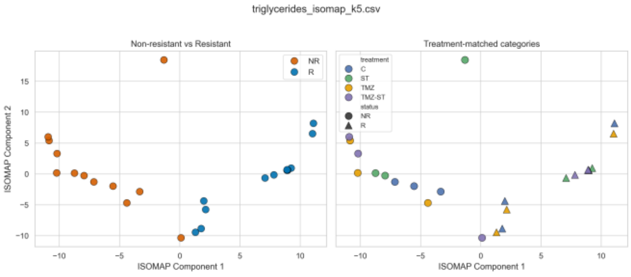
triglycerides_isomap_k5.csv
Non-resistant vs Resistant
Treatment-matched categories
ISOMAP Component 1
ISOMAP Component 2
NR
R
treatment
C
ST
TMZ
TMZ-ST
status
NR
R

**P.**

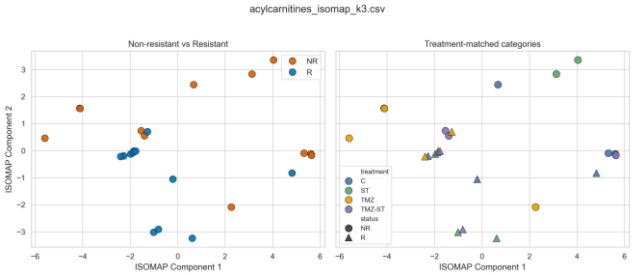
acylcarnitines_isomap_k3.csv
Non-resistant vs Resistant
Treatment-matched categories
ISOMAP Component 1
ISOMAP Component 2
NR
R
treatment
C
ST
TMZ
TMZ-ST
status
NR
R

**Q.**

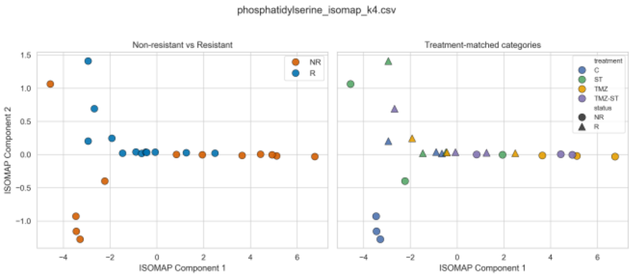


**R.**

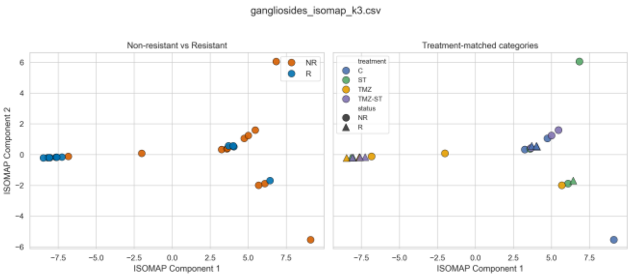


**S.**

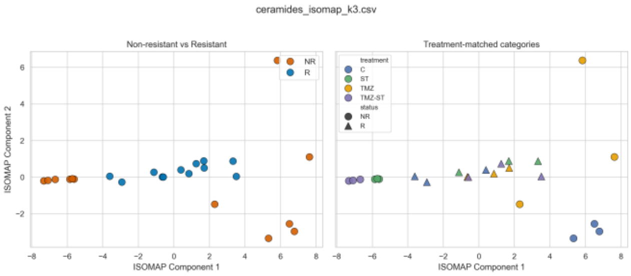


**Supplementary Figure S2**. ISOMAP embeddings of the lipid families. Isometric feature mapping projections of log2-transformed, z-score-normalized lipid intensities compare NR and R U251 glioblastoma cells across control, ST, TMZ, and TMZ-ST conditions. For each family, samples are displayed by resistance status and by treatment × status. These embeddings provide a nonlinear visualization complementary to the full-dimensional PERMANOVA analysis.

#### Supplementary Figure S3

**A.**

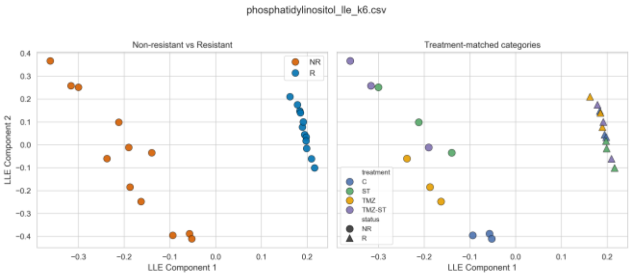


**B.**

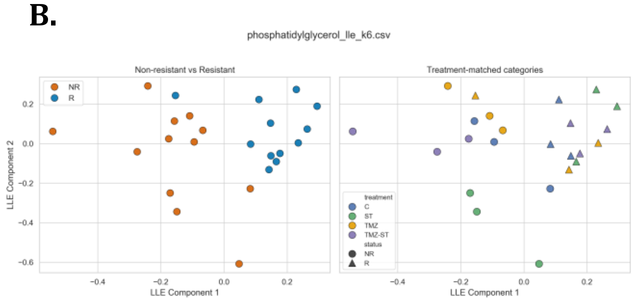


**C.**

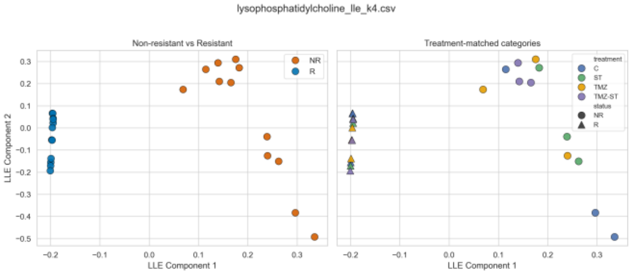


**D.**

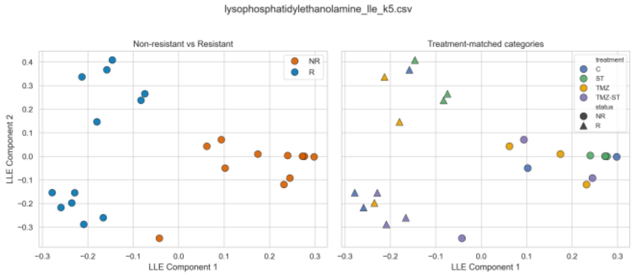


**E.**

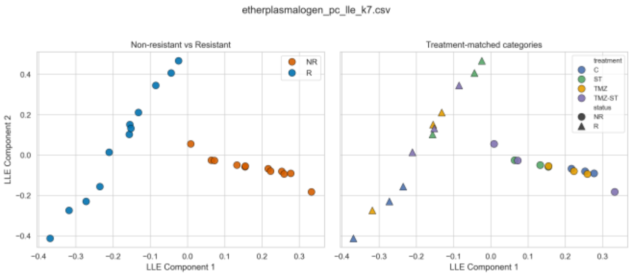


**F.**

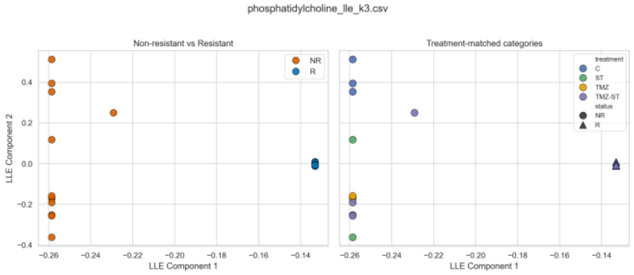


**G.**

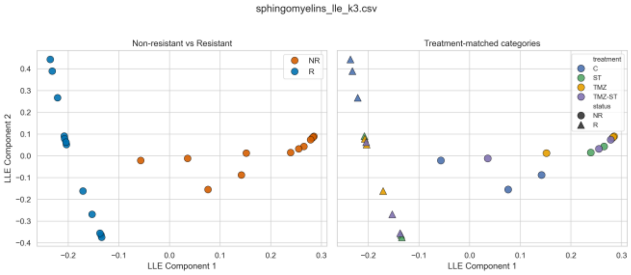


**H.**

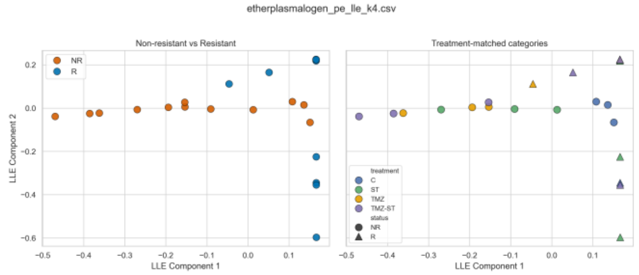

**I.**

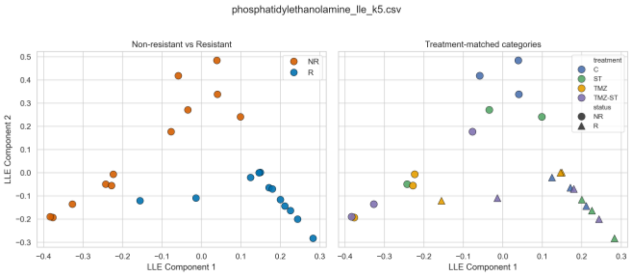
phosphatidylethanolamine_lle_k5.csv
Non-resistant vs Resistant
Treatment-matched categories
NR
R
treatment
C
ST
TMZ
TMZ-ST
status
NR
R
LLE Component 1
LLE Component 2

**J.**

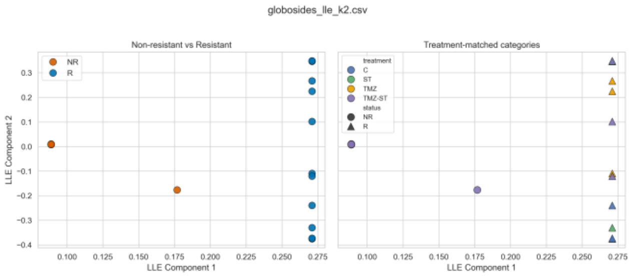
globosides_lle_k2.csv
Non-resistant vs Resistant
Treatment-matched categories
NR
R
treatment
C
ST
TMZ
TMZ-ST
status
NR
R
LLE Component 1
LLE Component 2

**K.**

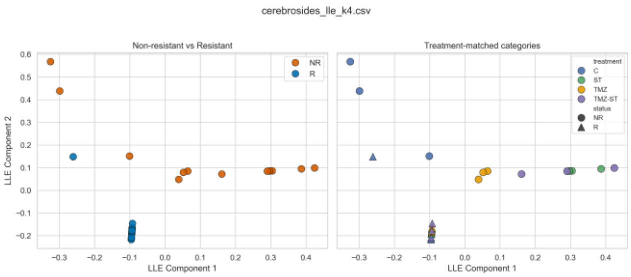
cerebrosides_lle_k4.csv
Non-resistant vs Resistant
Treatment-matched categories
NR
R
treatment
C
ST
TMZ
TMZ-ST
status
NR
R
LLE Component 1
LLE Component 2

**L.**

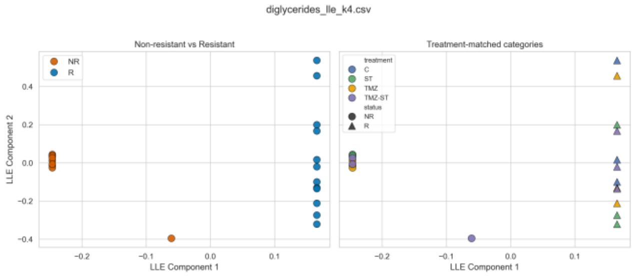
diglycerides_lle_k4.csv
Non-resistant vs Resistant
Treatment-matched categories
NR
R
treatment
C
ST
TMZ
TMZ-ST
status
NR
R
LLE Component 1
LLE Component 2

**M.**

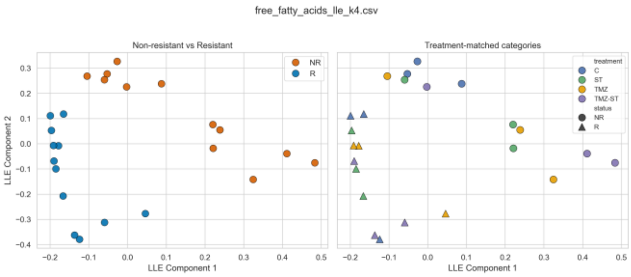
free_fatty_acids_lle_k4.csv
Non-resistant vs Resistant
Treatment-matched categories
NR
R
treatment
C
ST
TMZ
TMZ-ST
status
NR
R
LLE Component 1
LLE Component 2

**N.**

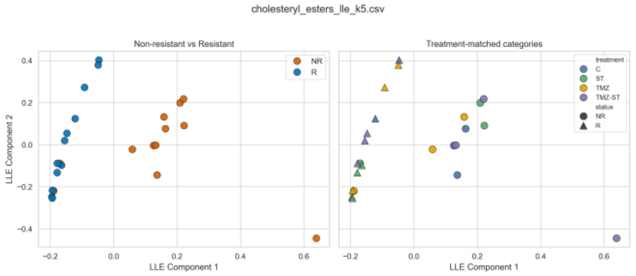
cholesteryl_esters_lle_k5.csv
Non-resistant vs Resistant
Treatment-matched categories
NR
R
treatment
C
ST
TMZ
TMZ-ST
status
NR
R
LLE Component 1
LLE Component 2

**O.**

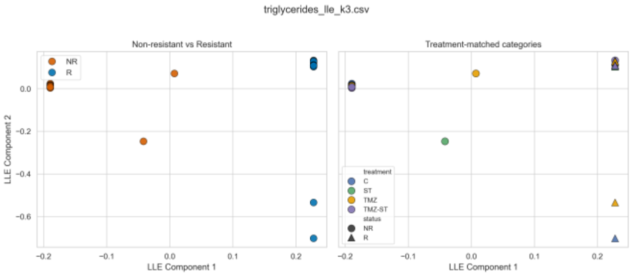
triglycerides_lle_k3.csv
Non-resistant vs Resistant
Treatment-matched categories
NR
R
treatment
C
ST
TMZ
TMZ-ST
status
NR
R
LLE Component 1
LLE Component 2

**P.**

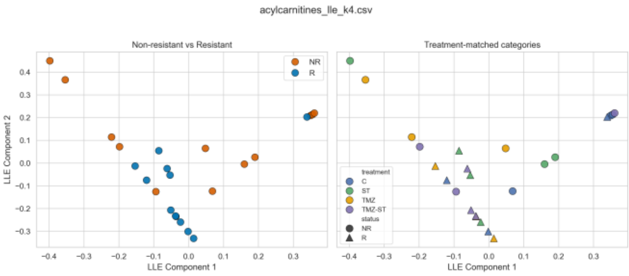
acylcarnitines_lle_k4.csv
Non-resistant vs Resistant
Treatment-matched categories
NR
R
treatment
C
ST
TMZ
TMZ-ST
status
NR
R
LLE Component 1
LLE Component 2

**Q.**

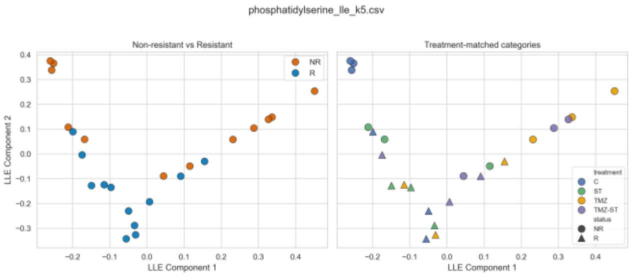


**R.**

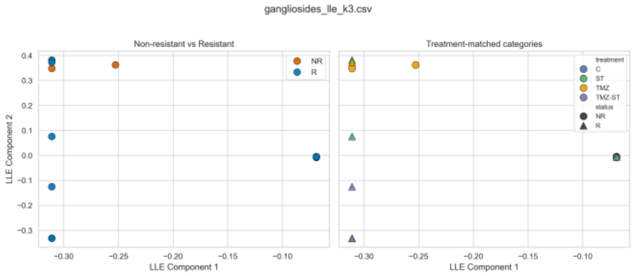


**S.**

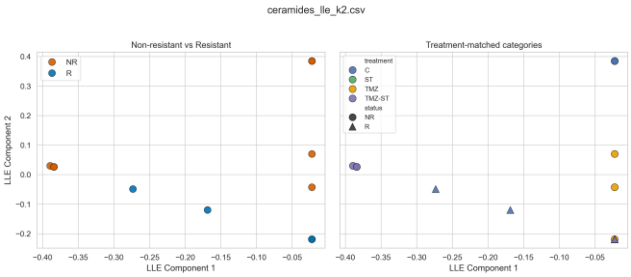


**Supplementary Figure S3.** LLE embeddings of the lipid families. Locally linear embedding projections of log2-transformed, z-score-normalized lipid intensities compare NR and R U251 glioblastoma cells across control, ST, TMZ, and TMZ-ST conditions. For each family, samples are displayed by resistance status and by treatment × status. LLE is used as a visualization/robustness tool rather than as the primary inferential statistic.

#### Supplementary Figure S4

**A.**

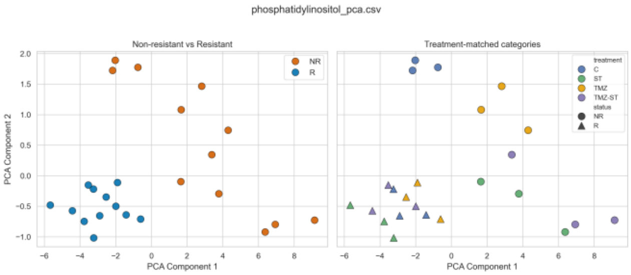


**B.**

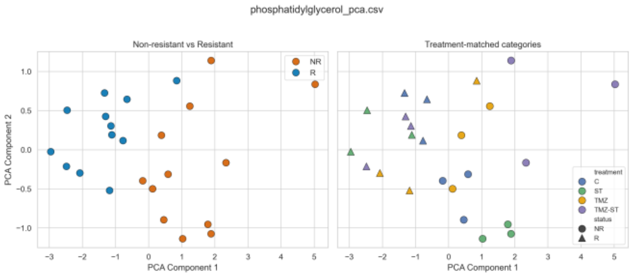


**C.**

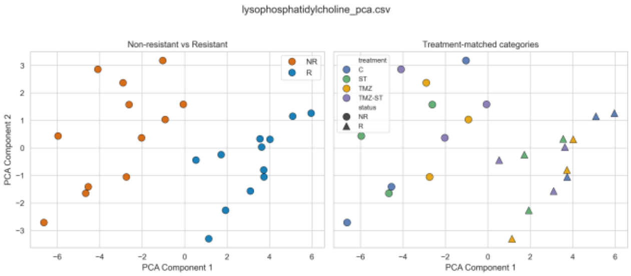


**D.**

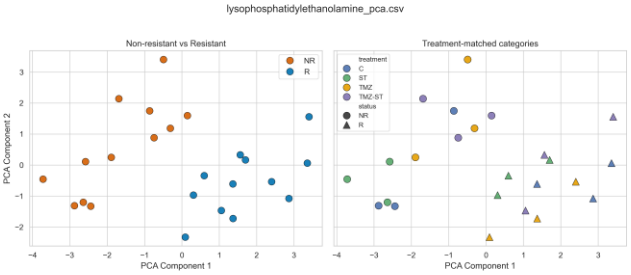


**E.**

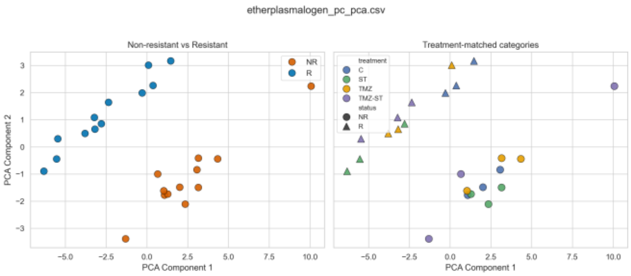


**F.**

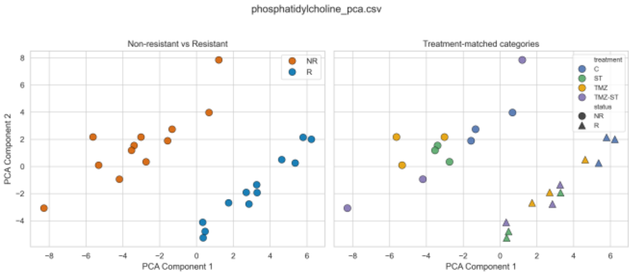


**G.**

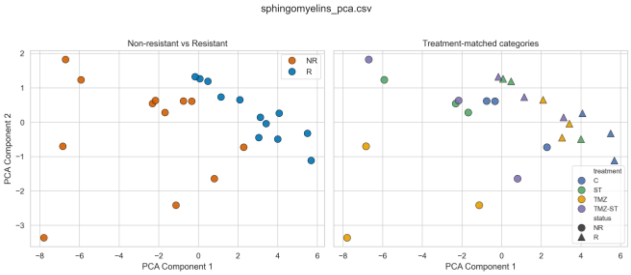


**H.**

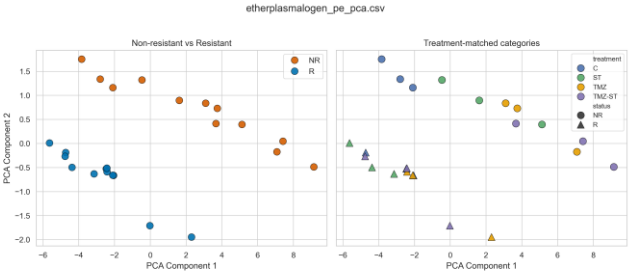

**I.**

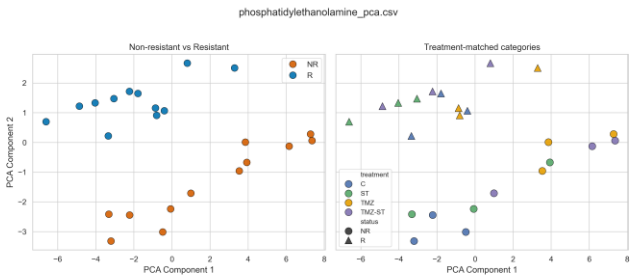
phosphatidylethanolamine_pca.csv
Non-resistant vs Resistant
Treatment-matched categories
NR
R
treatment
C
ST
TMZ
TMZ-ST
status
NR
R
PCA Component 1
PCA Component 2

**J.**

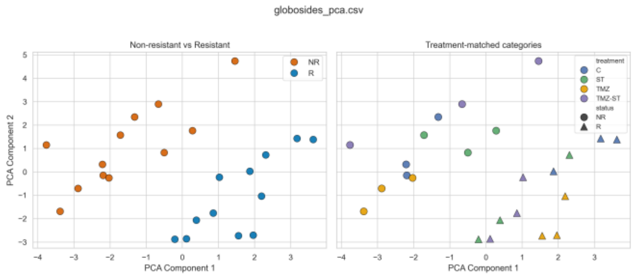
globosides_pca.csv
Non-resistant vs Resistant
Treatment-matched categories
NR
R
treatment
C
ST
TMZ
TMZ-ST
status
NR
R
PCA Component 1
PCA Component 2

**K.**

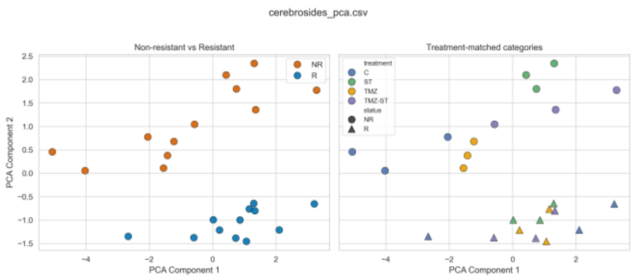
cerebrosides_pca.csv
Non-resistant vs Resistant
Treatment-matched categories
NR
R
treatment
C
ST
TMZ
TMZ-ST
status
NR
R
PCA Component 1
PCA Component 2

**L.**

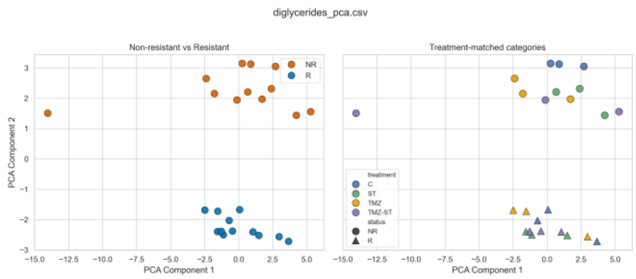
diglycerides_pca.csv
Non-resistant vs Resistant
Treatment-matched categories
NR
R
treatment
C
ST
TMZ
TMZ-ST
status
NR
R
PCA Component 1
PCA Component 2

**M.**

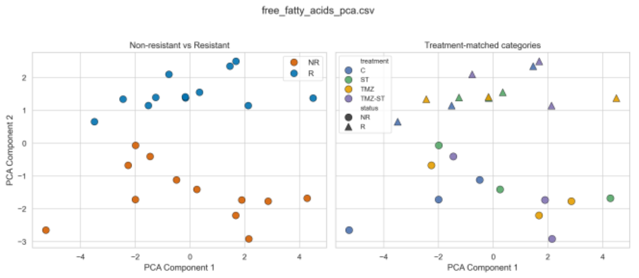
free_fatty_acids_pca.csv
Non-resistant vs Resistant
Treatment-matched categories
NR
R
treatment
C
ST
TMZ
TMZ-ST
status
NR
R
PCA Component 1
PCA Component 2

**N.**

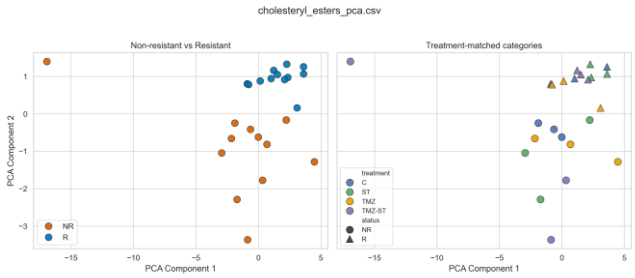
cholesteryl_esters_pca.csv
Non-resistant vs Resistant
Treatment-matched categories
NR
R
treatment
C
ST
TMZ
TMZ-ST
status
NR
R
PCA Component 1
PCA Component 2

**O.**

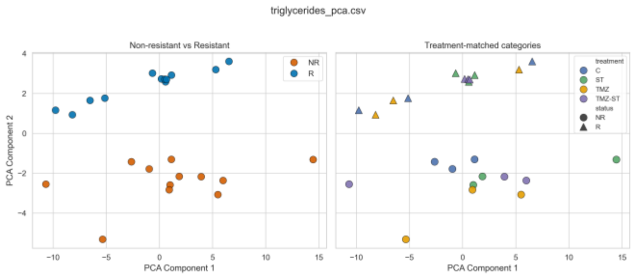
triglycerides_pca.csv
Non-resistant vs Resistant
Treatment-matched categories
NR
R
treatment
C
ST
TMZ
TMZ-ST
status
NR
R
PCA Component 1
PCA Component 2

**P.**

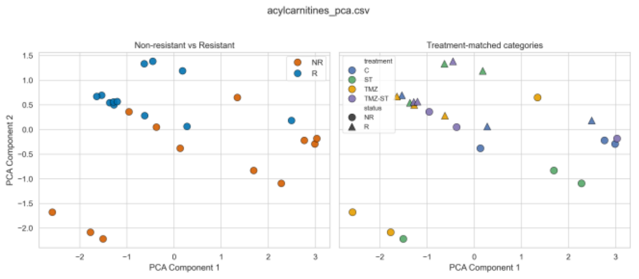
acylcarnitines_pca.csv
Non-resistant vs Resistant
Treatment-matched categories
NR
R
treatment
C
ST
TMZ
TMZ-ST
status
NR
R
PCA Component 1
PCA Component 2

**Q.**

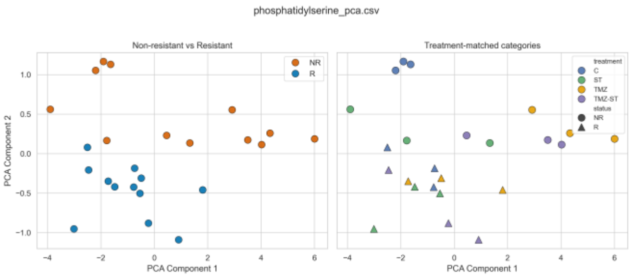


**R.**

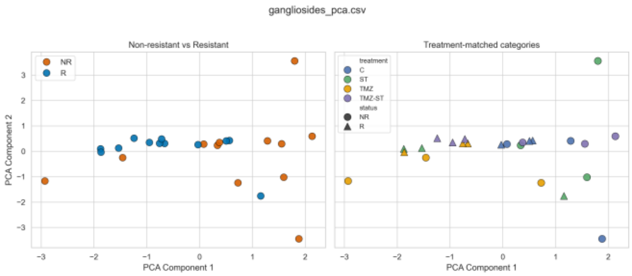


**S.**

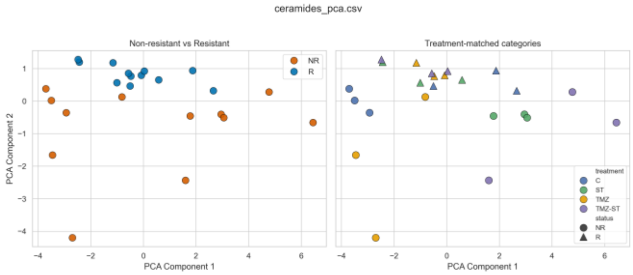


**Supplementary Figure S4**. PCA embeddings of the lipid families. PCA projections of log2-transformed, z-score-normalized lipid intensities compare NR and R U251 glioblastoma cells across control, ST, TMZ, and TMZ-ST conditions. For each family, samples are displayed by resistance status and by treatment × status. PCA provides the linear-projection complement to UMAP, ISOMAP, and LLE.

#### Supplementary Figure S5

**A.**

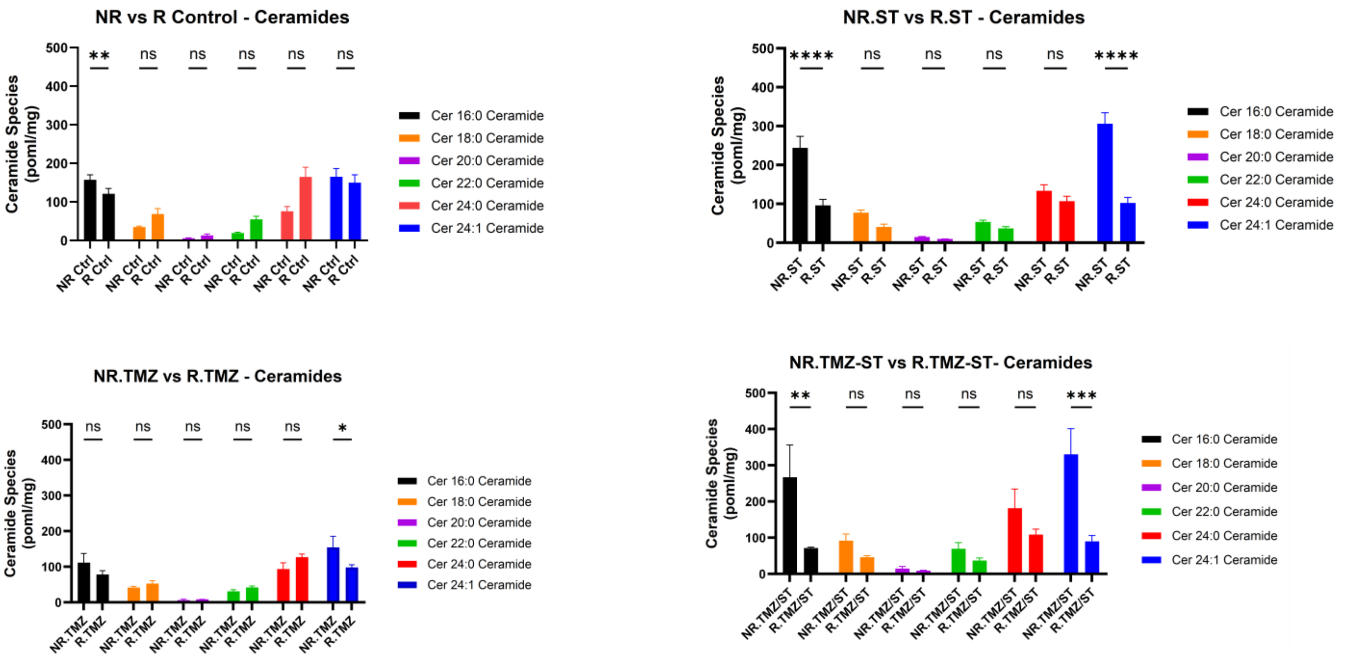


**B.**

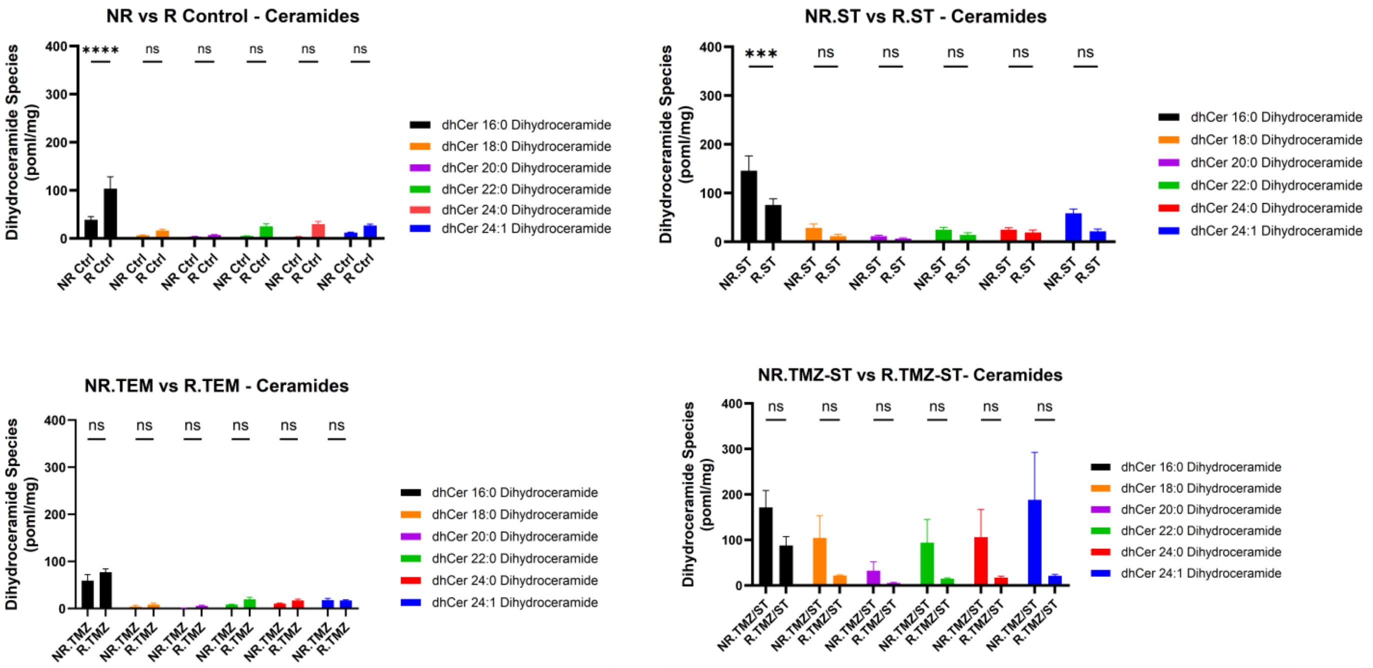

**C.**

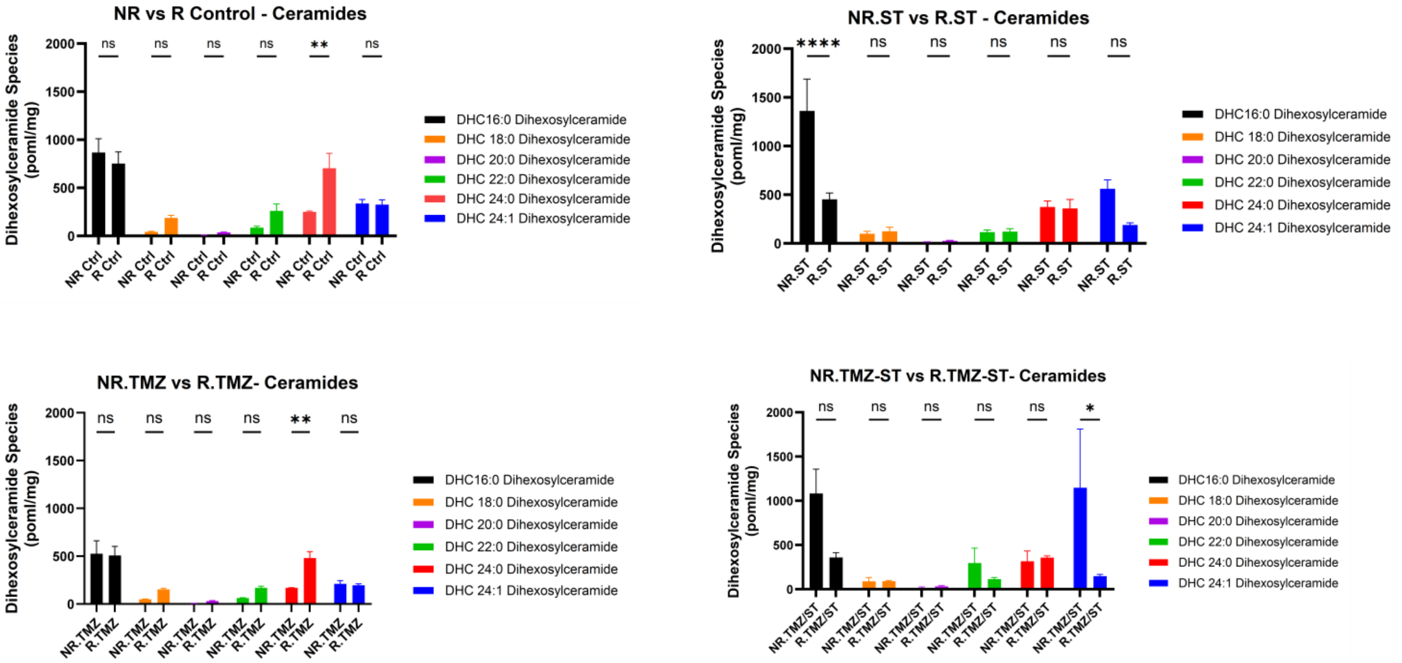
NR vs R Control - Ceramides
Dihexosylceramide Species (poml/mg)
ns ns ns ns ** ns
NR Ctrl R Ctrl
DHC16:0 Dihexosylceramide
DHC 18:0 Dihexosylceramide
DHC 20:0 Dihexosylceramide
DHC 22:0 Dihexosylceramide
DHC 24:0 Dihexosylceramide
DHC 24:1 Dihexosylceramide
NR.ST vs R.ST - Ceramides
Dihexosylceramide Species (poml/mg)
**** ns ns ns ns ns
NR.ST R.ST
DHC16:0 Dihexosylceramide
DHC 18:0 Dihexosylceramide
DHC 20:0 Dihexosylceramide
DHC 22:0 Dihexosylceramide
DHC 24:0 Dihexosylceramide
DHC 24:1 Dihexosylceramide
NR.TMZ vs R.TMZ- Ceramides
Dihexosylceramide Species (poml/mg)
ns ns ns ns ** ns
NR.TMZ R.TMZ
DHC16:0 Dihexosylceramide
DHC 18:0 Dihexosylceramide
DHC 20:0 Dihexosylceramide
DHC 22:0 Dihexosylceramide
DHC 24:0 Dihexosylceramide
DHC 24:1 Dihexosylceramide
NR.TMZ-ST vs R.TMZ-ST- Ceramides
Dihexosylceramide Species (poml/mg)
ns ns ns ns ns *
NR.TMZ/ST R.TMZ/ST
DHC16:0 Dihexosylceramide
DHC 18:0 Dihexosylceramide
DHC 20:0 Dihexosylceramide
DHC 22:0 Dihexosylceramide
DHC 24:0 Dihexosylceramide
DHC 24:1 Dihexosylceramide

**D.**

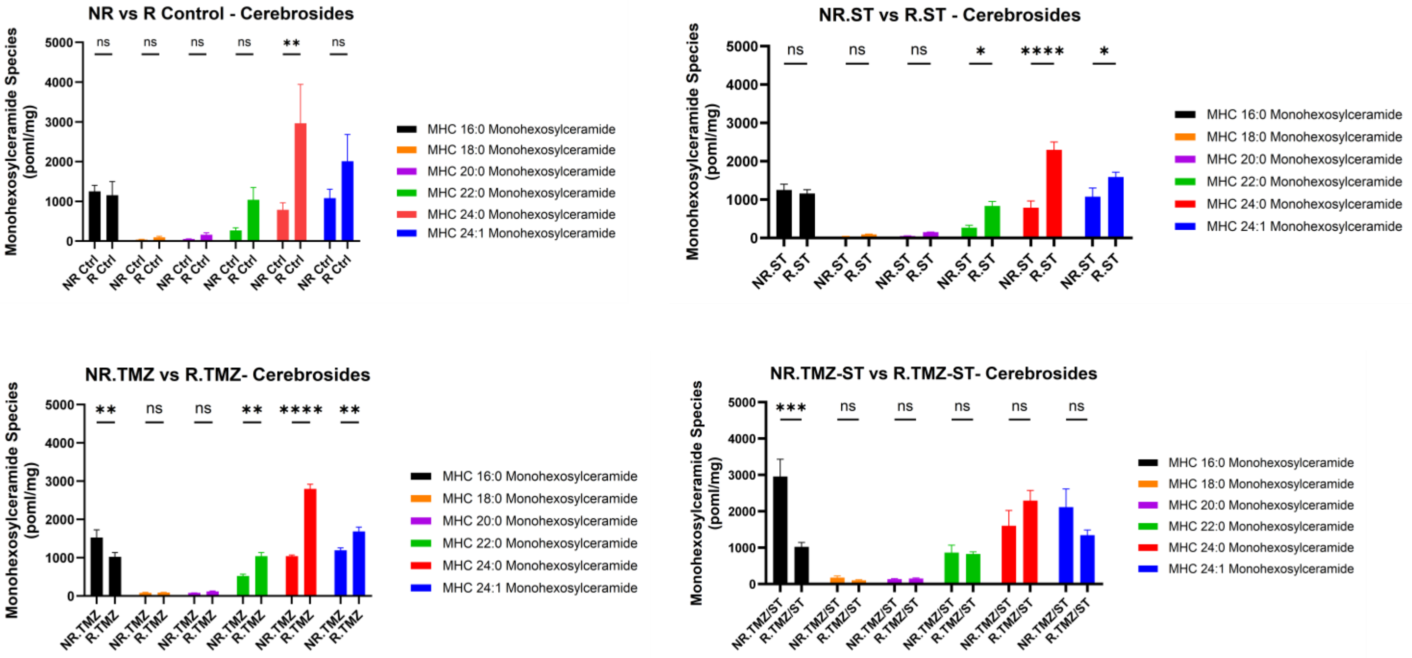
NR vs R Control - Cerebrosides
Monohexosylceramide Species (poml/mg)
ns ns ns ns ** ns
NR Ctrl R Ctrl
MHC 16:0 Monohexosylceramide
MHC 18:0 Monohexosylceramide
MHC 20:0 Monohexosylceramide
MHC 22:0 Monohexosylceramide
MHC 24:0 Monohexosylceramide
MHC 24:1 Monohexosylceramide
NR.ST vs R.ST - Cerebrosides
Monohexosylceramide Species (poml/mg)
ns ns ns * **** *
NR.ST R.ST
MHC 16:0 Monohexosylceramide
MHC 18:0 Monohexosylceramide
MHC 20:0 Monohexosylceramide
MHC 22:0 Monohexosylceramide
MHC 24:0 Monohexosylceramide
MHC 24:1 Monohexosylceramide
NR.TMZ vs R.TMZ- Cerebrosides
Monohexosylceramide Species (poml/mg)
** ns ns ** **** **
NR.TMZ R.TMZ
MHC 16:0 Monohexosylceramide
MHC 18:0 Monohexosylceramide
MHC 20:0 Monohexosylceramide
MHC 22:0 Monohexosylceramide
MHC 24:0 Monohexosylceramide
MHC 24:1 Monohexosylceramide
NR.TMZ-ST vs R.TMZ-ST- Cerebrosides
Monohexosylceramide Species (poml/mg)
*** ns ns ns ns ns
NR.TMZ/ST R.TMZ/ST
MHC 16:0 Monohexosylceramide
MHC 18:0 Monohexosylceramide
MHC 20:0 Monohexosylceramide
MHC 22:0 Monohexosylceramide
MHC 24:0 Monohexosylceramide
MHC 24:1 Monohexosylceramide

**E.**

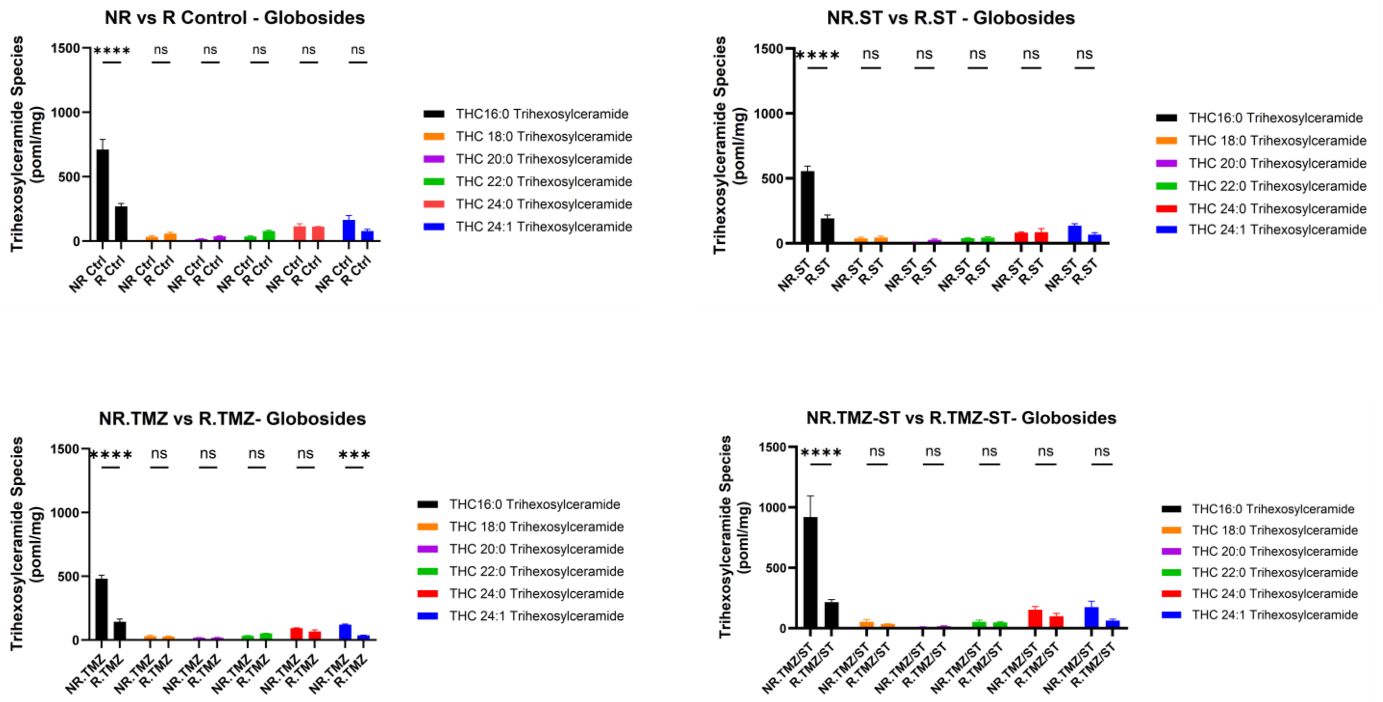
NR vs R Control - Globosides
Trihexosylceramide Species (poml/mg)
NR.ST vs R.ST - Globosides
NR.TMZ vs R.TMZ- Globosides
NR.TMZ-ST vs R.TMZ-ST- Globosides
THC16:0 Trihexosylceramide
THC 18:0 Trihexosylceramide
THC 20:0 Trihexosylceramide
THC 22:0 Trihexosylceramide
THC 24:0 Trihexosylceramide
THC 24:1 Trihexosylceramide
NR Ctrl
R Ctrl
NR.ST
R.ST
NR.TMZ
R.TMZ
NR.TMZ/ST
R.TMZ/ST
****
***
ns

**F.**

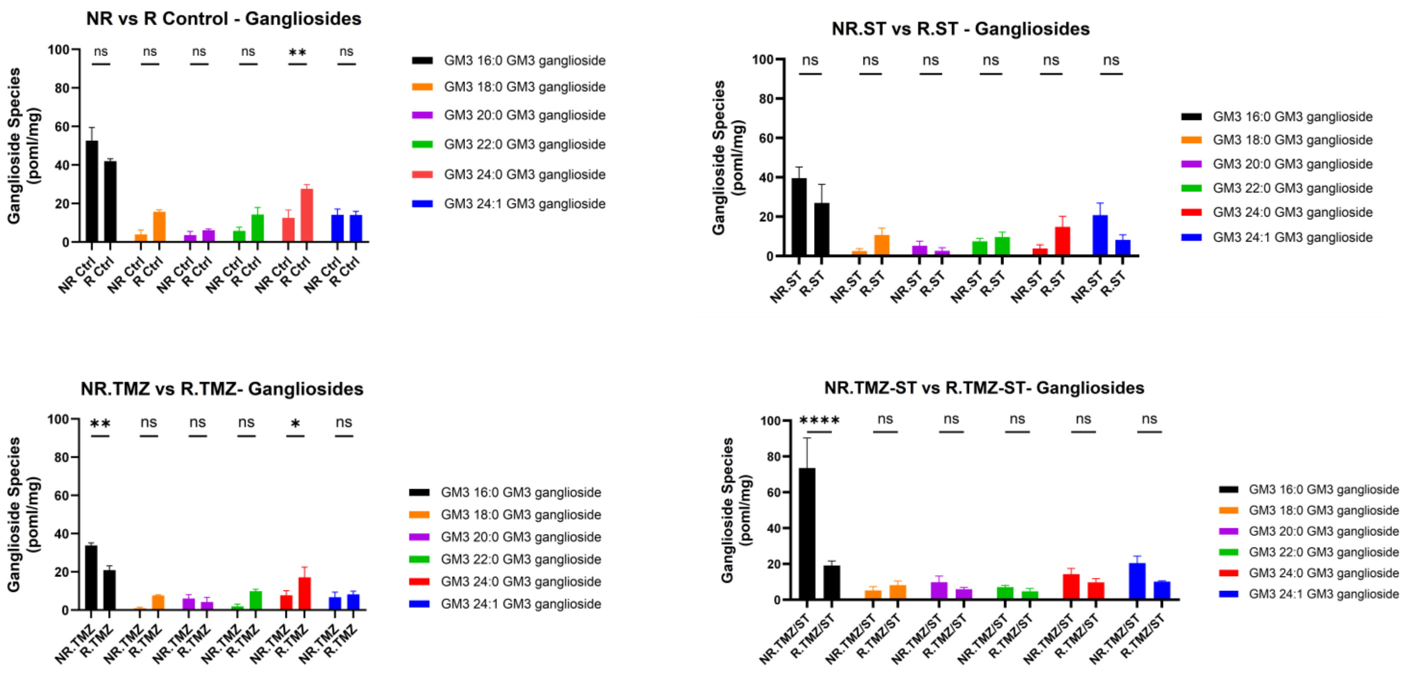
NR vs R Control - Gangliosides
Ganglioside Species (poml/mg)
NR.ST vs R.ST - Gangliosides
NR.TMZ vs R.TMZ- Gangliosides
NR.TMZ-ST vs R.TMZ-ST- Gangliosides
GM3 16:0 GM3 ganglioside
GM3 18:0 GM3 ganglioside
GM3 20:0 GM3 ganglioside
GM3 22:0 GM3 ganglioside
GM3 24:0 GM3 ganglioside
GM3 24:1 GM3 ganglioside
NR Ctrl
R Ctrl
NR.ST
R.ST
NR.TMZ
R.TMZ
NR.TMZ/ST
R.TMZ/ST
****
**
*
ns

**G.**

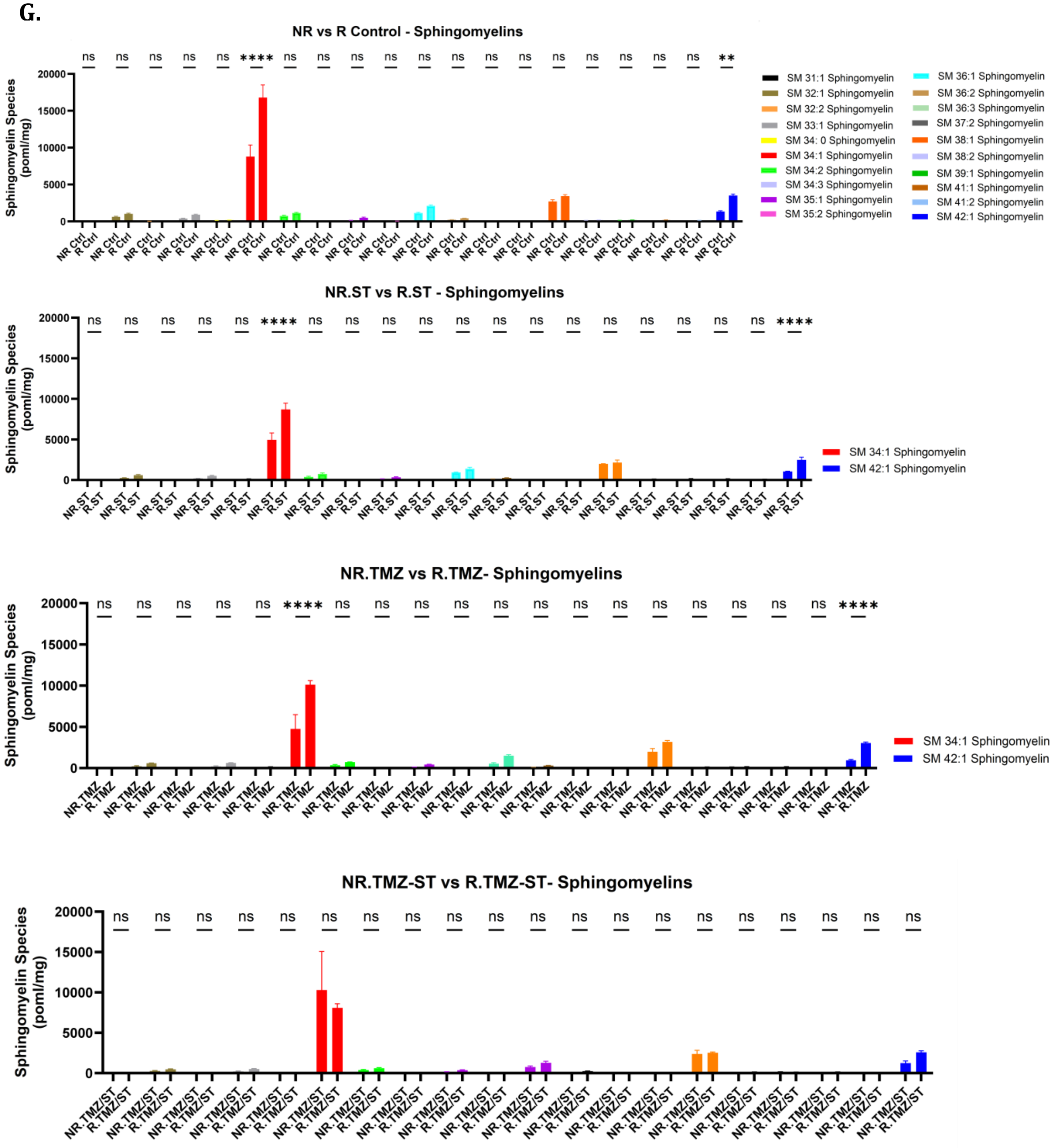
NR vs R Control - Sphingomyelins
Sphingomyelin Species (poml/mg)
ns ns ns ns ns **** ns ns ns ns ns ns ns ns ns ns ns ns ns **
NR Ctrl
R Ctrl
SM 31:1 Sphingomyelin
SM 32:1 Sphingomyelin
SM 32:2 Sphingomyelin
SM 33:1 Sphingomyelin
SM 34: 0 Sphingomyelin
SM 34:1 Sphingomyelin
SM 34:2 Sphingomyelin
SM 34:3 Sphingomyelin
SM 35:1 Sphingomyelin
SM 35:2 Sphingomyelin
SM 36:1 Sphingomyelin
SM 36:2 Sphingomyelin
SM 36:3 Sphingomyelin
SM 37:2 Sphingomyelin
SM 38:1 Sphingomyelin
SM 38:2 Sphingomyelin
SM 39:1 Sphingomyelin
SM 41:1 Sphingomyelin
SM 41:2 Sphingomyelin
SM 42:1 Sphingomyelin
NR.ST vs R.ST - Sphingomyelins
Sphingomyelin Species (poml/mg)
ns ns ns ns ns **** ns ns ns ns ns ns ns ns ns ns ns ns ns ****
NR.ST
R.ST
SM 34:1 Sphingomyelin
SM 42:1 Sphingomyelin
NR.TMZ vs R.TMZ- Sphingomyelins
Sphingomyelin Species (poml/mg)
ns ns ns ns ns **** ns ns ns ns ns ns ns ns ns ns ns ns ns ****
NR.TMZ
R.TMZ
SM 34:1 Sphingomyelin
SM 42:1 Sphingomyelin
NR.TMZ-ST vs R.TMZ-ST- Sphingomyelins
Sphingomyelin Species (poml/mg)
ns ns ns ns ns ns ns ns ns ns ns ns ns ns ns ns ns ns ns ns
NR.TMZ/ST
R.TMZ/ST
0
5000
10000
15000
20000

**H.**

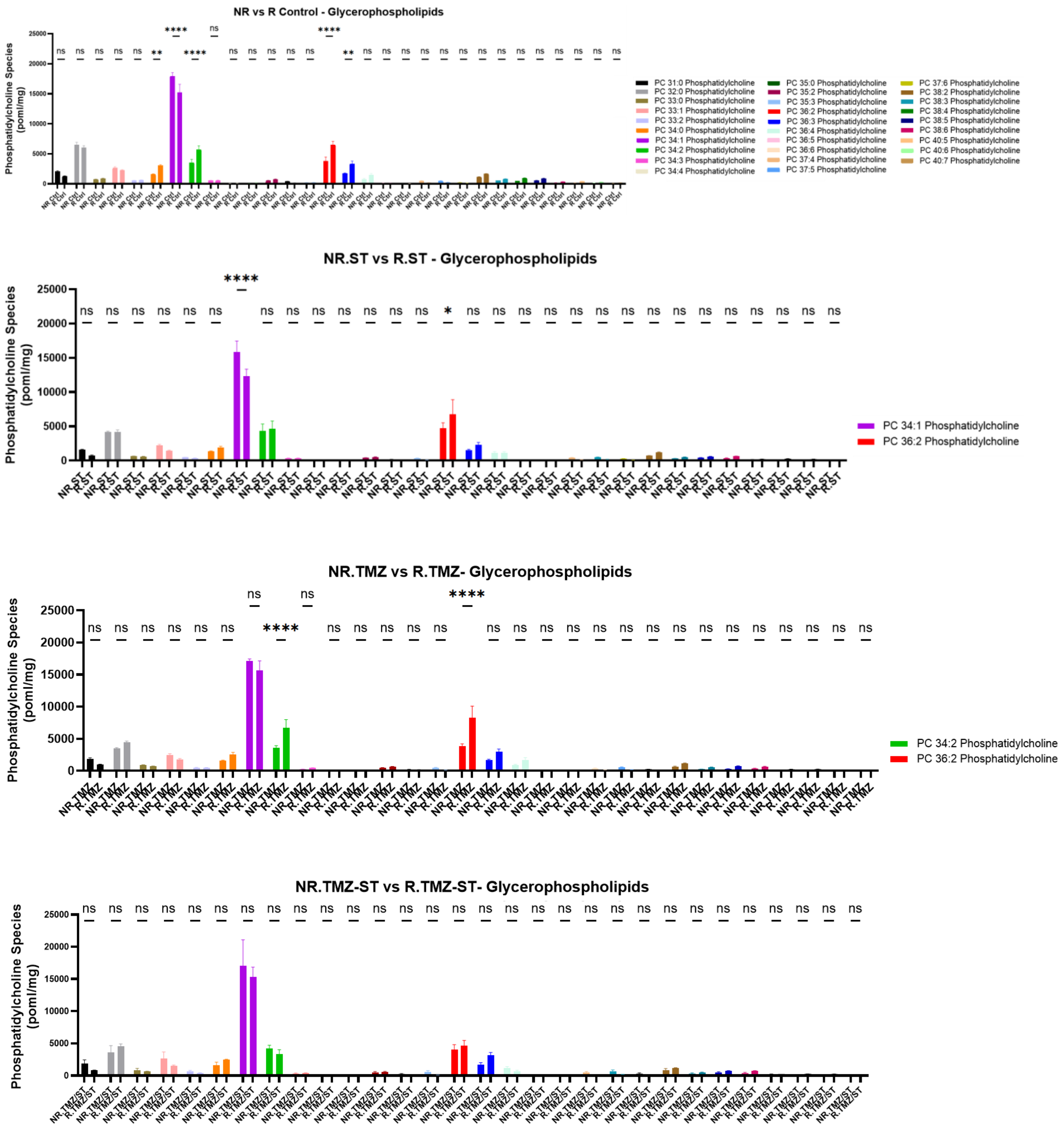

NR vs R Control - Glycerophospholipids
Phosphatidylcholine Species (poml/mg)
PC 31:0 Phosphatidylcholine
PC 32:0 Phosphatidylcholine
PC 33:0 Phosphatidylcholine
PC 33:1 Phosphatidylcholine
PC 33:2 Phosphatidylcholine
PC 34:0 Phosphatidylcholine
PC 34:1 Phosphatidylcholine
PC 34:2 Phosphatidylcholine
PC 34:3 Phosphatidylcholine
PC 34:4 Phosphatidylcholine
PC 35:0 Phosphatidylcholine
PC 35:2 Phosphatidylcholine
PC 35:3 Phosphatidylcholine
PC 36:2 Phosphatidylcholine
PC 36:3 Phosphatidylcholine
PC 36:4 Phosphatidylcholine
PC 36:5 Phosphatidylcholine
PC 36:6 Phosphatidylcholine
PC 37:4 Phosphatidylcholine
PC 37:5 Phosphatidylcholine
PC 37:6 Phosphatidylcholine
PC 38:2 Phosphatidylcholine
PC 38:3 Phosphatidylcholine
PC 38:4 Phosphatidylcholine
PC 38:5 Phosphatidylcholine
PC 38:6 Phosphatidylcholine
PC 40:5 Phosphatidylcholine
PC 40:6 Phosphatidylcholine
PC 40:7 Phosphatidylcholine
NR.ST vs R.ST - Glycerophospholipids
Phosphatidylcholine Species (poml/mg)
PC 34:1 Phosphatidylcholine
PC 36:2 Phosphatidylcholine
NR.TMZ vs R.TMZ- Glycerophospholipids
Phosphatidylcholine Species (poml/mg)
PC 34:2 Phosphatidylcholine
PC 36:2 Phosphatidylcholine
NR.TMZ-ST vs R.TMZ-ST- Glycerophospholipids
Phosphatidylcholine Species (poml/mg)

**I.**

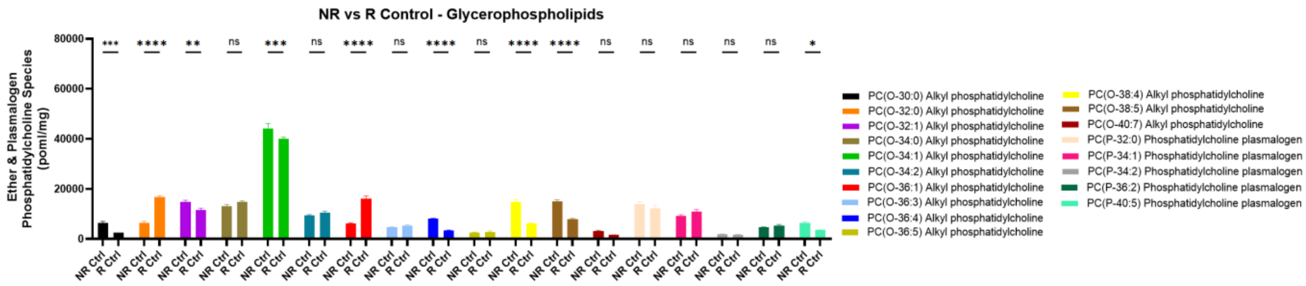
NR vs R Control - Glycerophospholipids
Ether & Plasmalogen Phosphatidylcholine Species (poml/mg)
PC(O-30:0) Alkyl phosphatidylcholine
PC(O-32:0) Alkyl phosphatidylcholine
PC(O-32:1) Alkyl phosphatidylcholine
PC(O-34:0) Alkyl phosphatidylcholine
PC(O-34:1) Alkyl phosphatidylcholine
PC(O-34:2) Alkyl phosphatidylcholine
PC(O-36:1) Alkyl phosphatidylcholine
PC(O-36:3) Alkyl phosphatidylcholine
PC(O-36:4) Alkyl phosphatidylcholine
PC(O-36:5) Alkyl phosphatidylcholine
PC(O-38:4) Alkyl phosphatidylcholine
PC(O-38:5) Alkyl phosphatidylcholine
PC(O-40:7) Alkyl phosphatidylcholine
PC(P-32:0) Phosphatidylcholine plasmalogen
PC(P-34:1) Phosphatidylcholine plasmalogen
PC(P-34:2) Phosphatidylcholine plasmalogen
PC(P-36:2) Phosphatidylcholine plasmalogen
PC(P-40:5) Phosphatidylcholine plasmalogen

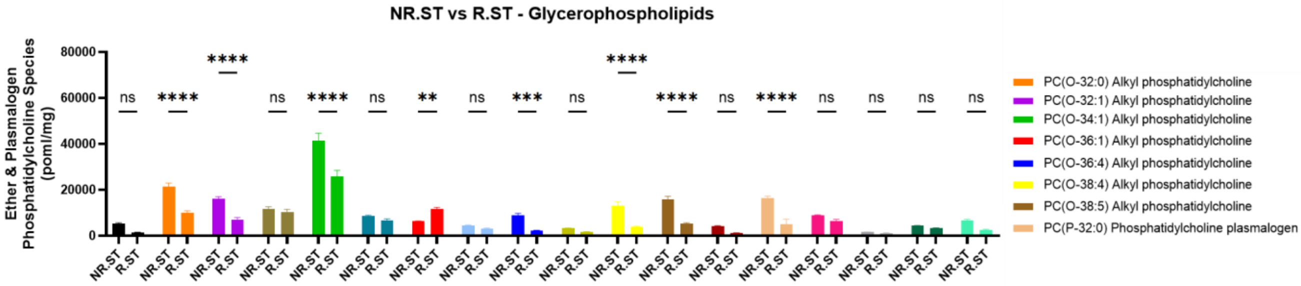
NR.ST vs R.ST - Glycerophospholipids
Ether & Plasmalogen Phosphatidylcholine Species (poml/mg)
PC(O-32:0) Alkyl phosphatidylcholine
PC(O-32:1) Alkyl phosphatidylcholine
PC(O-34:1) Alkyl phosphatidylcholine
PC(O-36:1) Alkyl phosphatidylcholine
PC(O-36:4) Alkyl phosphatidylcholine
PC(O-38:4) Alkyl phosphatidylcholine
PC(O-38:5) Alkyl phosphatidylcholine
PC(P-32:0) Phosphatidylcholine plasmalogen

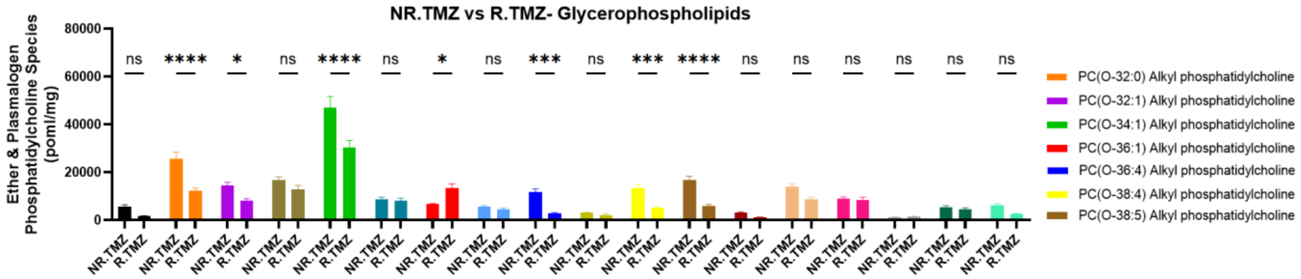
NR.TMZ vs R.TMZ- Glycerophospholipids
Ether & Plasmalogen Phosphatidylcholine Species (poml/mg)
PC(O-32:0) Alkyl phosphatidylcholine
PC(O-32:1) Alkyl phosphatidylcholine
PC(O-34:1) Alkyl phosphatidylcholine
PC(O-36:1) Alkyl phosphatidylcholine
PC(O-36:4) Alkyl phosphatidylcholine
PC(O-38:4) Alkyl phosphatidylcholine
PC(O-38:5) Alkyl phosphatidylcholine

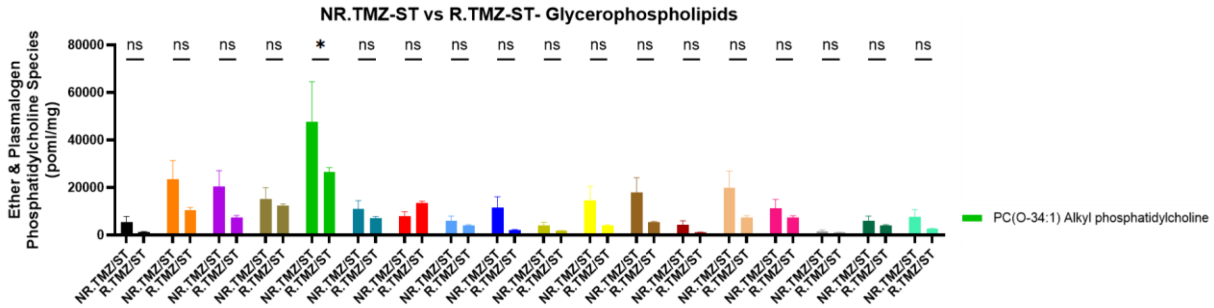
NR.TMZ-ST vs R.TMZ-ST- Glycerophospholipids
Ether & Plasmalogen Phosphatidylcholine Species (poml/mg)
PC(O-34:1) Alkyl phosphatidylcholine

**J.**

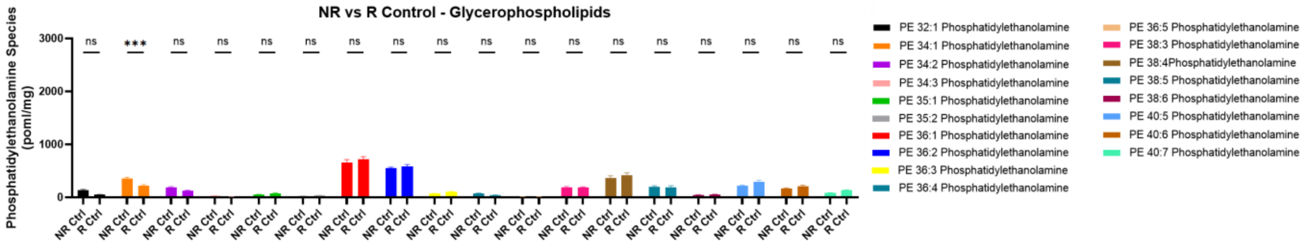
NR vs R Control - Glycerophospholipids
Phosphatidylethanolamine Species (poml/mg)
PE 32:1 Phosphatidylethanolamine
PE 34:1 Phosphatidylethanolamine
PE 34:2 Phosphatidylethanolamine
PE 34:3 Phosphatidylethanolamine
PE 35:1 Phosphatidylethanolamine
PE 35:2 Phosphatidylethanolamine
PE 36:1 Phosphatidylethanolamine
PE 36:2 Phosphatidylethanolamine
PE 36:3 Phosphatidylethanolamine
PE 36:4 Phosphatidylethanolamine
PE 36:5 Phosphatidylethanolamine
PE 38:3 Phosphatidylethanolamine
PE 38:4Phosphatidylethanolamine
PE 38:5 Phosphatidylethanolamine
PE 38:6 Phosphatidylethanolamine
PE 40:5 Phosphatidylethanolamine
PE 40:6 Phosphatidylethanolamine
PE 40:7 Phosphatidylethanolamine

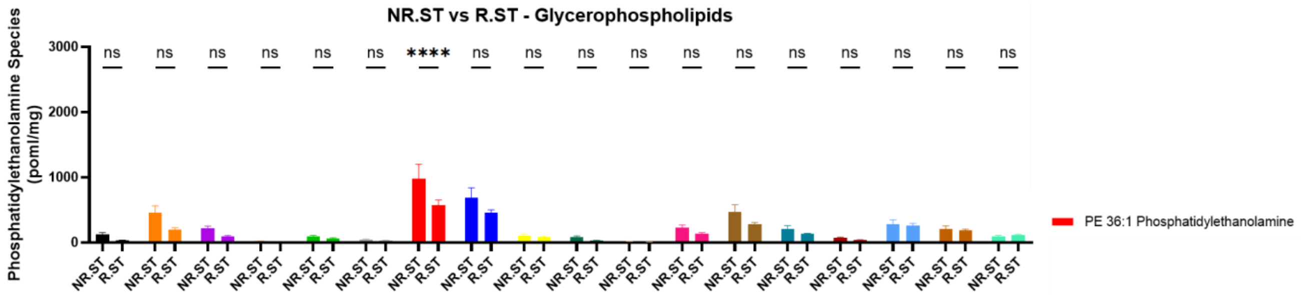
NR.ST vs R.ST - Glycerophospholipids
Phosphatidylethanolamine Species (poml/mg)
PE 36:1 Phosphatidylethanolamine

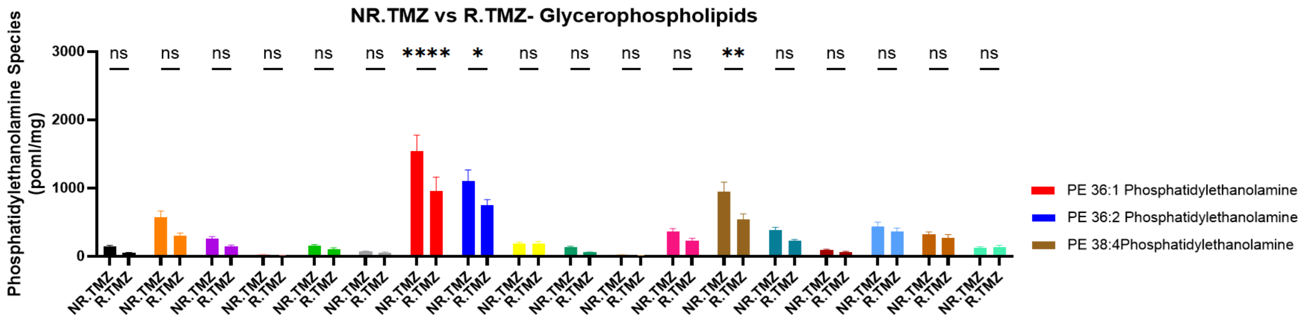
NR.TMZ vs R.TMZ- Glycerophospholipids
Phosphatidylethanolamine Species (poml/mg)
PE 36:1 Phosphatidylethanolamine
PE 36:2 Phosphatidylethanolamine
PE 38:4Phosphatidylethanolamine

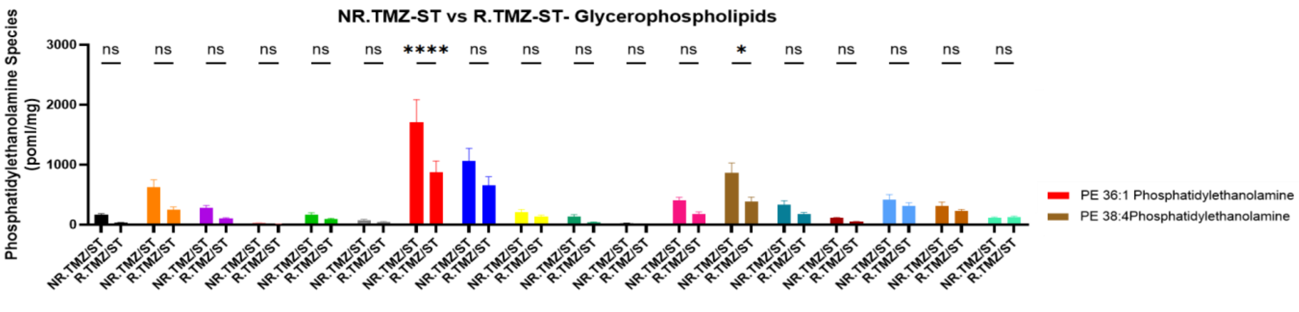
NR.TMZ-ST vs R.TMZ-ST- Glycerophospholipids
Phosphatidylethanolamine Species (poml/mg)
PE 36:1 Phosphatidylethanolamine
PE 38:4Phosphatidylethanolamine

**K.**

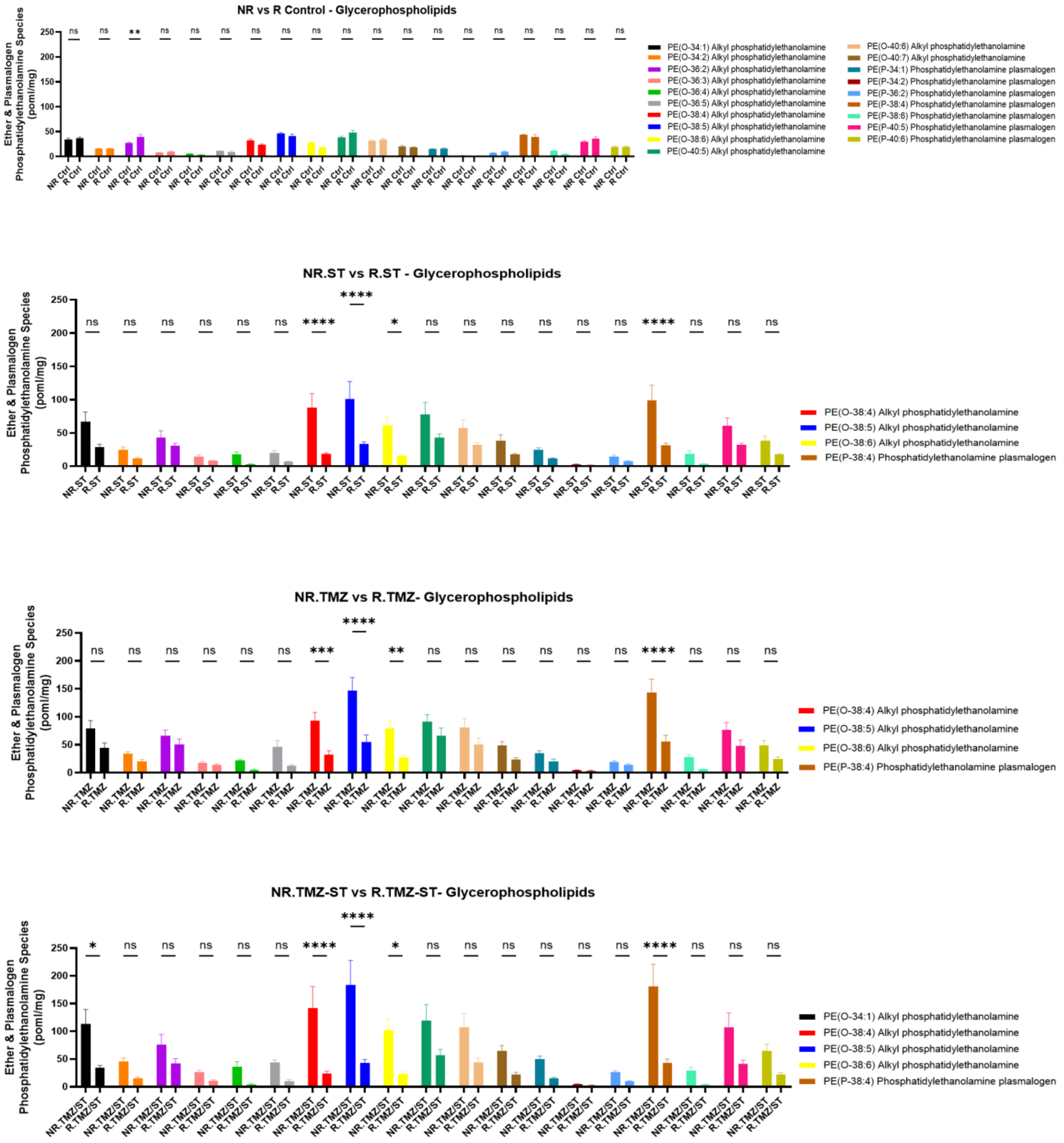
NR vs R Control - Glycerophospholipids
Ether & Plasmalogen Phosphatidylethanolamine Species (poml/mg)
PE(O-34:1) Alkyl phosphatidylethanolamine
PE(O-34:2) Alkyl phosphatidylethanolamine
PE(O-36:2) Alkyl phosphatidylethanolamine
PE(O-36:3) Alkyl phosphatidylethanolamine
PE(O-36:4) Alkyl phosphatidylethanolamine
PE(O-36:5) Alkyl phosphatidylethanolamine
PE(O-38:4) Alkyl phosphatidylethanolamine
PE(O-38:5) Alkyl phosphatidylethanolamine
PE(O-38:6) Alkyl phosphatidylethanolamine
PE(O-40:5) Alkyl phosphatidylethanolamine
PE(O-40:6) Alkyl phosphatidylethanolamine
PE(O-40:7) Alkyl phosphatidylethanolamine
PE(P-34:1) Phosphatidylethanolamine plasmalogen
PE(P-34:2) Phosphatidylethanolamine plasmalogen
PE(P-36:2) Phosphatidylethanolamine plasmalogen
PE(P-38:4) Phosphatidylethanolamine plasmalogen
PE(P-38:6) Phosphatidylethanolamine plasmalogen
PE(P-40:5) Phosphatidylethanolamine plasmalogen
PE(P-40:6) Phosphatidylethanolamine plasmalogen
NR.ST vs R.ST - Glycerophospholipids
PE(O-38:4) Alkyl phosphatidylethanolamine
PE(O-38:5) Alkyl phosphatidylethanolamine
PE(O-38:6) Alkyl phosphatidylethanolamine
PE(P-38:4) Phosphatidylethanolamine plasmalogen
NR.TMZ vs R.TMZ- Glycerophospholipids
PE(O-38:4) Alkyl phosphatidylethanolamine
PE(O-38:5) Alkyl phosphatidylethanolamine
PE(O-38:6) Alkyl phosphatidylethanolamine
PE(P-38:4) Phosphatidylethanolamine plasmalogen
NR.TMZ-ST vs R.TMZ-ST- Glycerophospholipids
PE(O-34:1) Alkyl phosphatidylethanolamine
PE(O-38:4) Alkyl phosphatidylethanolamine
PE(O-38:5) Alkyl phosphatidylethanolamine
PE(O-38:6) Alkyl phosphatidylethanolamine
PE(P-38:4) Phosphatidylethanolamine plasmalogen

**L.**

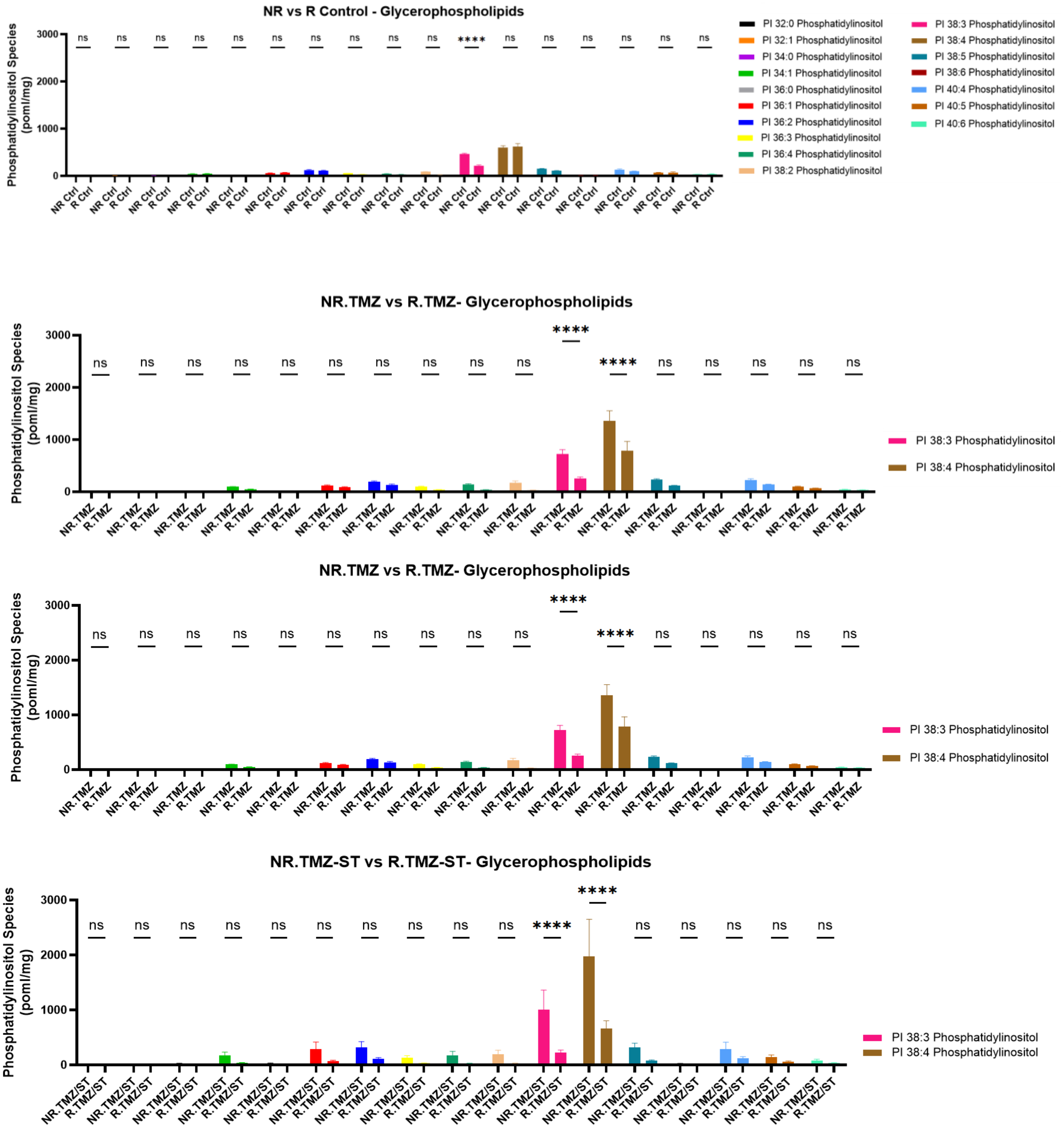
NR vs R Control - Glycerophospholipids
Phosphatidylinositol Species (poml/mg)
3000
2000
1000
0
NR Ctrl
R Ctrl
****
ns
PI 32:0 Phosphatidylinositol
PI 32:1 Phosphatidylinositol
PI 34:0 Phosphatidylinositol
PI 34:1 Phosphatidylinositol
PI 36:0 Phosphatidylinositol
PI 36:1 Phosphatidylinositol
PI 36:2 Phosphatidylinositol
PI 36:3 Phosphatidylinositol
PI 36:4 Phosphatidylinositol
PI 38:2 Phosphatidylinositol
PI 38:3 Phosphatidylinositol
PI 38:4 Phosphatidylinositol
PI 38:5 Phosphatidylinositol
PI 38:6 Phosphatidylinositol
PI 40:4 Phosphatidylinositol
PI 40:5 Phosphatidylinositol
PI 40:6 Phosphatidylinositol
NR.TMZ vs R.TMZ- Glycerophospholipids
Phosphatidylinositol Species (poml/mg)
NR.TMZ
R.TMZ
PI 38:3 Phosphatidylinositol
PI 38:4 Phosphatidylinositol
NR.TMZ vs R.TMZ- Glycerophospholipids
Phosphatidylinositol Species (poml/mg)
NR.TMZ
R.TMZ
PI 38:3 Phosphatidylinositol
PI 38:4 Phosphatidylinositol
NR.TMZ-ST vs R.TMZ-ST- Glycerophospholipids
Phosphatidylinositol Species (poml/mg)
NR.TMZ/ST
R.TMZ/ST
PI 38:3 Phosphatidylinositol
PI 38:4 Phosphatidylinositol

**M.**

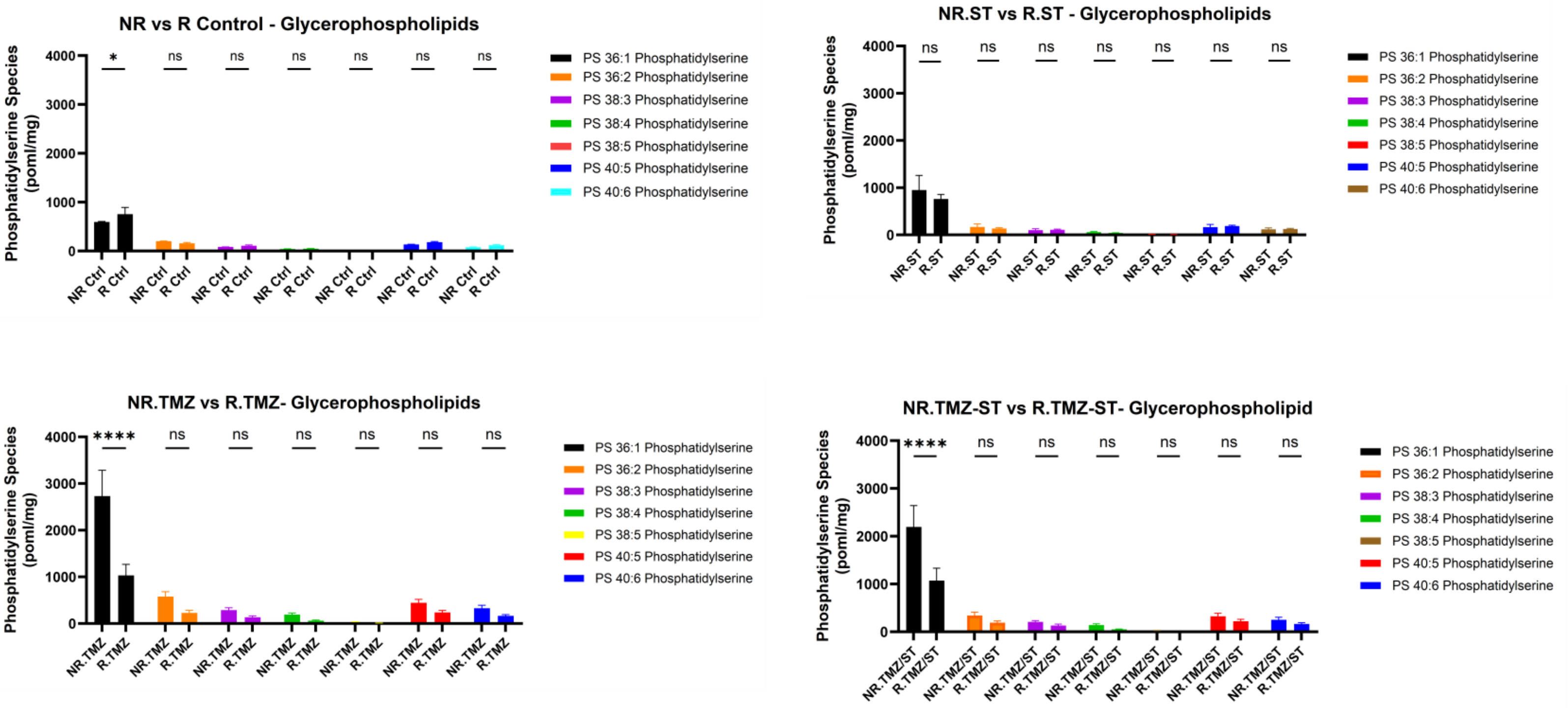
NR vs R Control - Glycerophospholipids
NR.ST vs R.ST - Glycerophospholipids
NR.TMZ vs R.TMZ- Glycerophospholipids
NR.TMZ-ST vs R.TMZ-ST- Glycerophospholipid
Phosphatidylserine Species (poml/mg)
PS 36:1 Phosphatidylserine
PS 36:2 Phosphatidylserine
PS 38:3 Phosphatidylserine
PS 38:4 Phosphatidylserine
PS 38:5 Phosphatidylserine
PS 40:5 Phosphatidylserine
PS 40:6 Phosphatidylserine
NR Ctrl
R Ctrl
NR.ST
R.ST
NR.TMZ
R.TMZ
NR.TMZ/ST
R.TMZ/ST
ns
*
****

**N.**

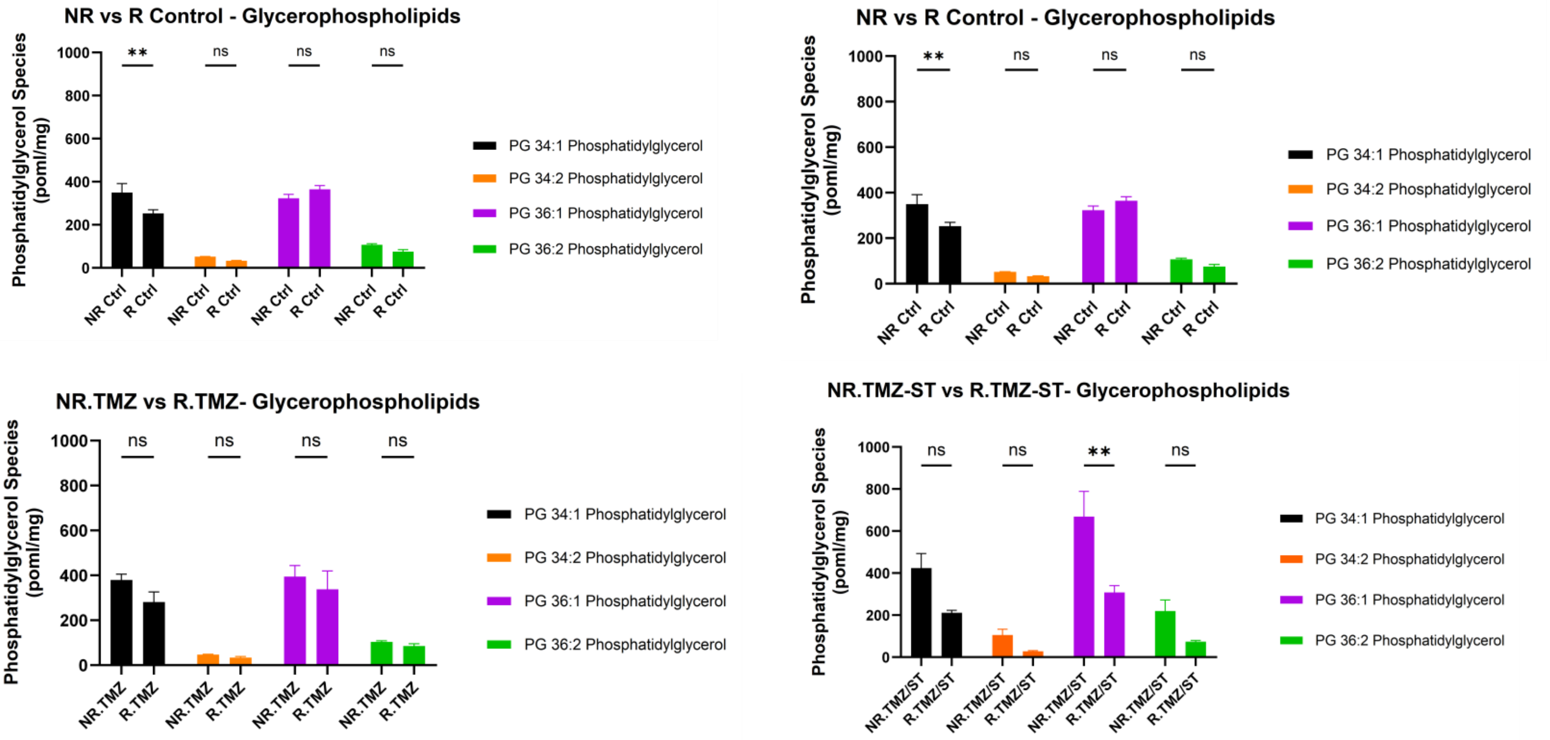
NR vs R Control - Glycerophospholipids
NR vs R Control - Glycerophospholipids
NR.TMZ vs R.TMZ- Glycerophospholipids
NR.TMZ-ST vs R.TMZ-ST- Glycerophospholipids
Phosphatidylglycerol Species (poml/mg)
PG 34:1 Phosphatidylglycerol
PG 34:2 Phosphatidylglycerol
PG 36:1 Phosphatidylglycerol
PG 36:2 Phosphatidylglycerol
NR Ctrl
R Ctrl
NR.TMZ
R.TMZ
NR.TMZ/ST
R.TMZ/ST
ns
**

O.

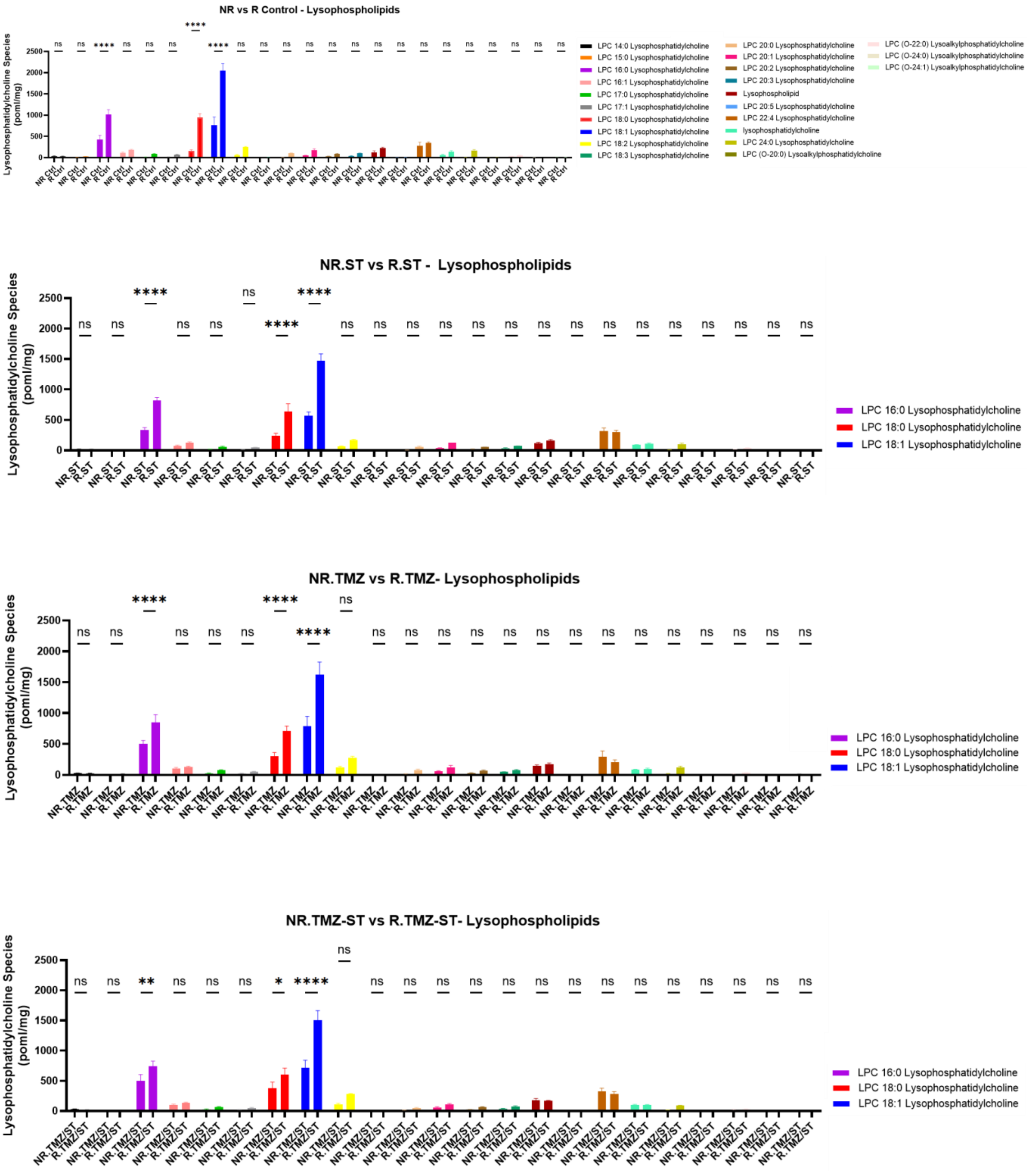
NR vs R Control - Lysophospholipids
Lysophosphatidylcholine Species (poml/mg)
LPC 14:0 Lysophosphatidylcholine
LPC 15:0 Lysophosphatidylcholine
LPC 16:0 Lysophosphatidylcholine
LPC 16:1 Lysophosphatidylcholine
LPC 17:0 Lysophosphatidylcholine
LPC 17:1 Lysophosphatidylcholine
LPC 18:0 Lysophosphatidylcholine
LPC 18:1 Lysophosphatidylcholine
LPC 18:2 Lysophosphatidylcholine
LPC 18:3 Lysophosphatidylcholine
LPC 20:0 Lysophosphatidylcholine
LPC 20:1 Lysophosphatidylcholine
LPC 20:2 Lysophosphatidylcholine
LPC 20:3 Lysophosphatidylcholine
Lysophospholipid
LPC 20:5 Lysophosphatidylcholine
LPC 22:4 Lysophosphatidylcholine
lysophosphatidylcholine
LPC 24:0 Lysophosphatidylcholine
LPC (O-20:0) Lysoalkylphosphatidylcholine
LPC (O-22:0) Lysoalkylphosphatidylcholine
LPC (O-24:0) Lysoalkylphosphatidylcholine
LPC (O-24:1) Lysoalkylphosphatidylcholine
NR.ST vs R.ST - Lysophospholipids
Lysophosphatidylcholine Species (poml/mg)
LPC 16:0 Lysophosphatidylcholine
LPC 18:0 Lysophosphatidylcholine
LPC 18:1 Lysophosphatidylcholine
NR.TMZ vs R.TMZ- Lysophospholipids
Lysophosphatidylcholine Species (poml/mg)
LPC 16:0 Lysophosphatidylcholine
LPC 18:0 Lysophosphatidylcholine
LPC 18:1 Lysophosphatidylcholine
NR.TMZ-ST vs R.TMZ-ST- Lysophospholipids
Lysophosphatidylcholine Species (poml/mg)
LPC 16:0 Lysophosphatidylcholine
LPC 18:0 Lysophosphatidylcholine
LPC 18:1 Lysophosphatidylcholine

**P.**

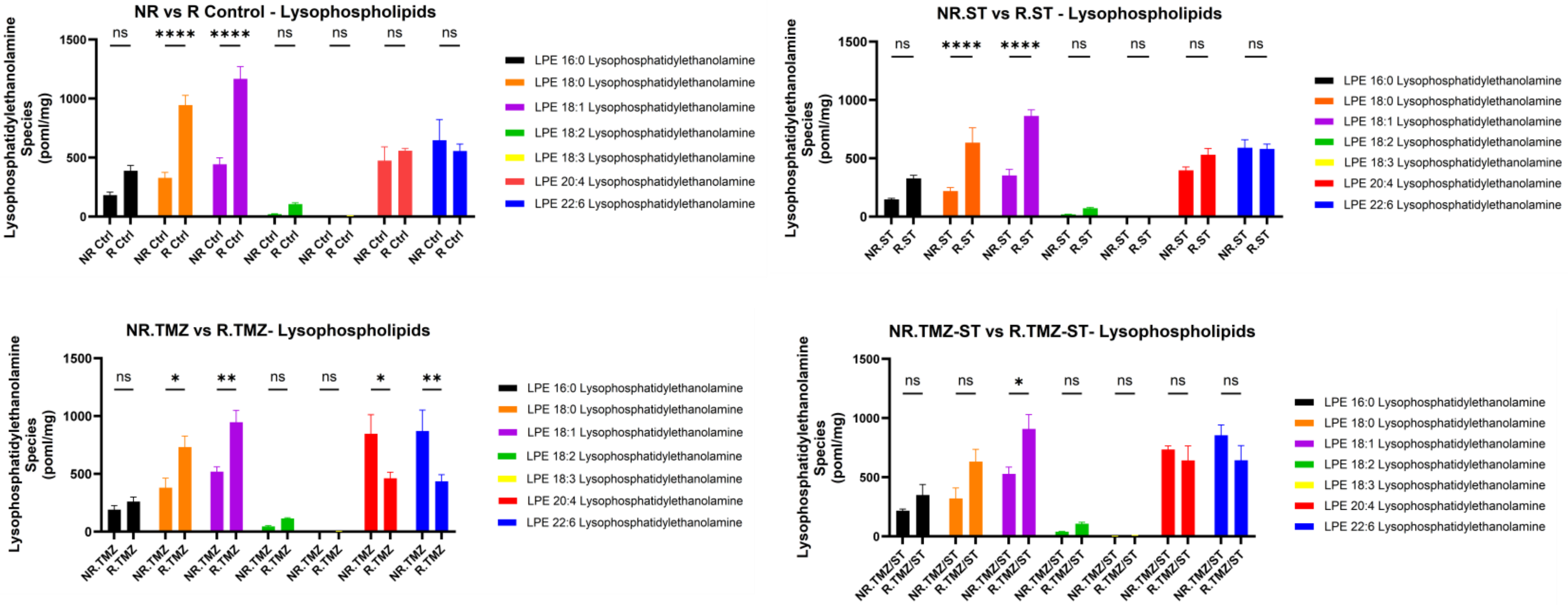
NR vs R Control - Lysophospholipids
NR.ST vs R.ST - Lysophospholipids
NR.TMZ vs R.TMZ- Lysophospholipids
NR.TMZ-ST vs R.TMZ-ST- Lysophospholipids
Lysophosphatidylethanolamine Species (poml/mg)
ns **** **** ns ns ns ns
ns **** **** ns ns ns ns
ns * ** ns ns * **
ns ns * ns ns ns ns
LPE 16:0 Lysophosphatidylethanolamine
LPE 18:0 Lysophosphatidylethanolamine
LPE 18:1 Lysophosphatidylethanolamine
LPE 18:2 Lysophosphatidylethanolamine
LPE 18:3 Lysophosphatidylethanolamine
LPE 20:4 Lysophosphatidylethanolamine
LPE 22:6 Lysophosphatidylethanolamine

**Q.**

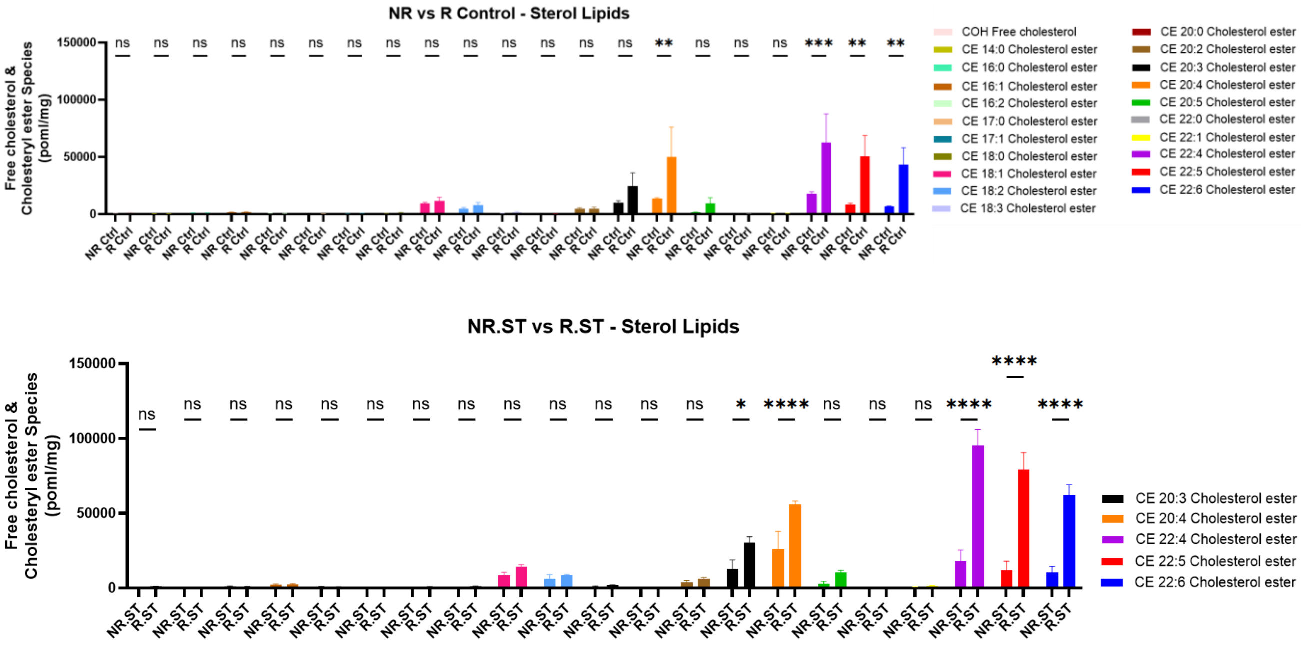
NR vs R Control - Sterol Lipids
NR.ST vs R.ST - Sterol Lipids
Free cholesterol & Cholesteryl ester Species (poml/mg)
ns ns ns ns ns ns ns ns ns ns ns ns ns ns ** ns ns ns *** ** **
ns ns ns ns ns ns ns ns ns ns ns ns ns * **** ns ns ns **** **** ****
COH Free cholesterol
CE 14:0 Cholesterol ester
CE 16:0 Cholesterol ester
CE 16:1 Cholesterol ester
CE 16:2 Cholesterol ester
CE 17:0 Cholesterol ester
CE 17:1 Cholesterol ester
CE 18:0 Cholesterol ester
CE 18:1 Cholesterol ester
CE 18:2 Cholesterol ester
CE 18:3 Cholesterol ester
CE 20:0 Cholesterol ester
CE 20:2 Cholesterol ester
CE 20:3 Cholesterol ester
CE 20:4 Cholesterol ester
CE 20:5 Cholesterol ester
CE 22:0 Cholesterol ester
CE 22:1 Cholesterol ester
CE 22:4 Cholesterol ester
CE 22:5 Cholesterol ester
CE 22:6 Cholesterol ester

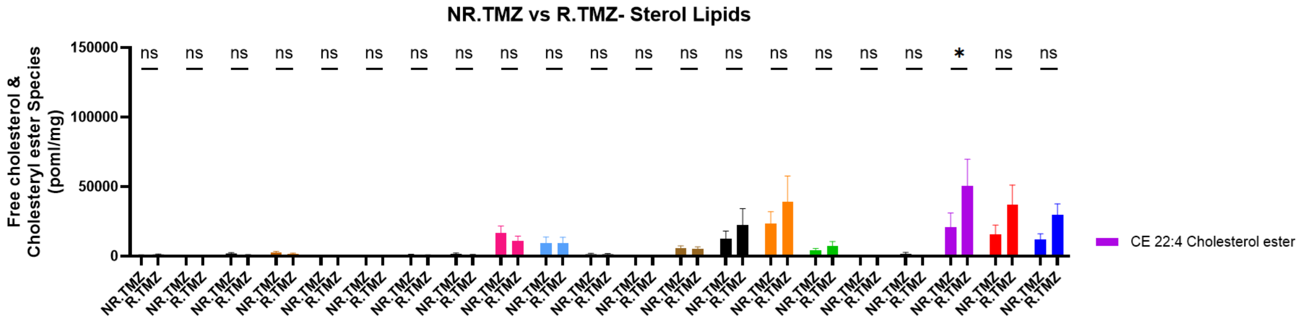
NR.TMZ vs R.TMZ- Sterol Lipids
Free cholesterol & Cholesteryl ester Species (poml/mg)
CE 22:4 Cholesterol ester

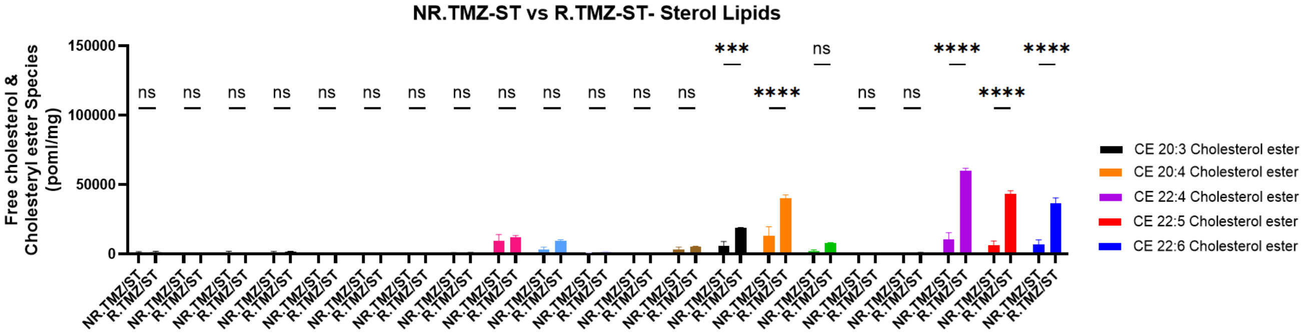
NR.TMZ-ST vs R.TMZ-ST- Sterol Lipids
Free cholesterol & Cholesteryl ester Species (poml/mg)
CE 20:3 Cholesterol ester
CE 20:4 Cholesterol ester
CE 22:4 Cholesterol ester
CE 22:5 Cholesterol ester
CE 22:6 Cholesterol ester

**R.**

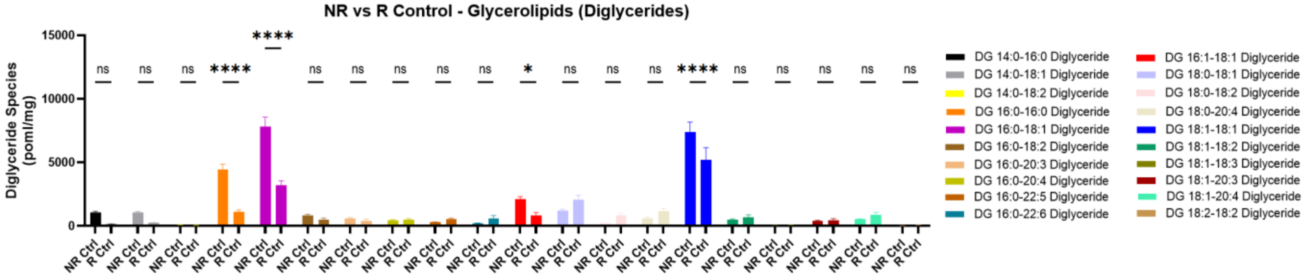
NR vs R Control - Glycerolipids (Diglycerides)
Diglyceride Species (poml/mg)
DG 14:0-16:0 Diglyceride
DG 14:0-18:1 Diglyceride
DG 14:0-18:2 Diglyceride
DG 16:0-16:0 Diglyceride
DG 16:0-18:1 Diglyceride
DG 16:0-18:2 Diglyceride
DG 16:0-20:3 Diglyceride
DG 16:0-20:4 Diglyceride
DG 16:0-22:5 Diglyceride
DG 16:0-22:6 Diglyceride
DG 16:1-18:1 Diglyceride
DG 18:0-18:1 Diglyceride
DG 18:0-18:2 Diglyceride
DG 18:0-20:4 Diglyceride
DG 18:1-18:1 Diglyceride
DG 18:1-18:2 Diglyceride
DG 18:1-18:3 Diglyceride
DG 18:1-20:3 Diglyceride
DG 18:1-20:4 Diglyceride
DG 18:2-18:2 Diglyceride

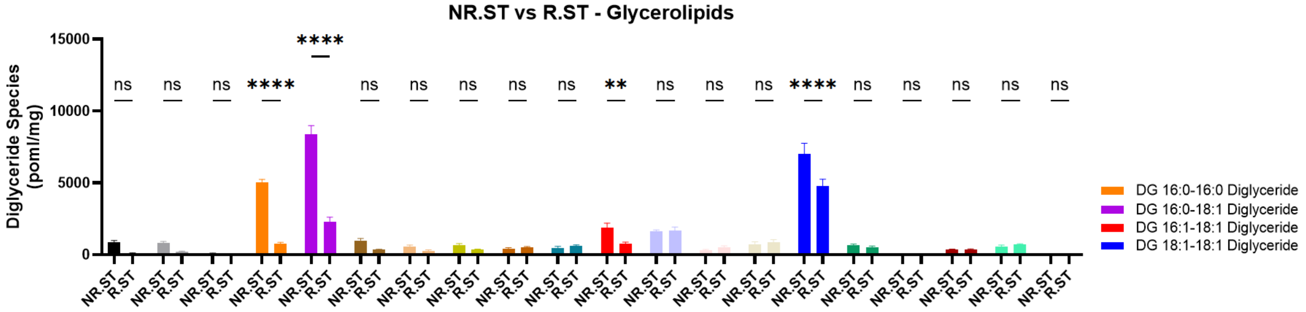
NR.ST vs R.ST - Glycerolipids
Diglyceride Species (poml/mg)
DG 16:0-16:0 Diglyceride
DG 16:0-18:1 Diglyceride
DG 16:1-18:1 Diglyceride
DG 18:1-18:1 Diglyceride

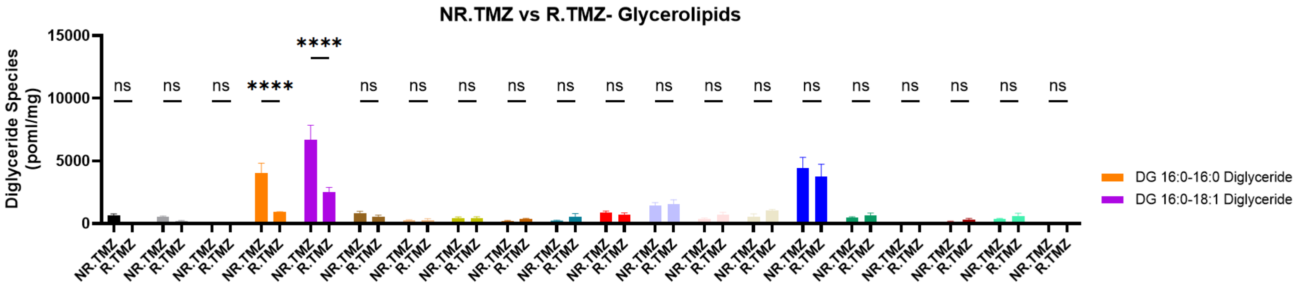
NR.TMZ vs R.TMZ- Glycerolipids
Diglyceride Species (poml/mg)
15000
10000
5000
0
ns
****
DG 16:0-16:0 Diglyceride
DG 16:0-18:1 Diglyceride
NR.TMZ
R.TMZ

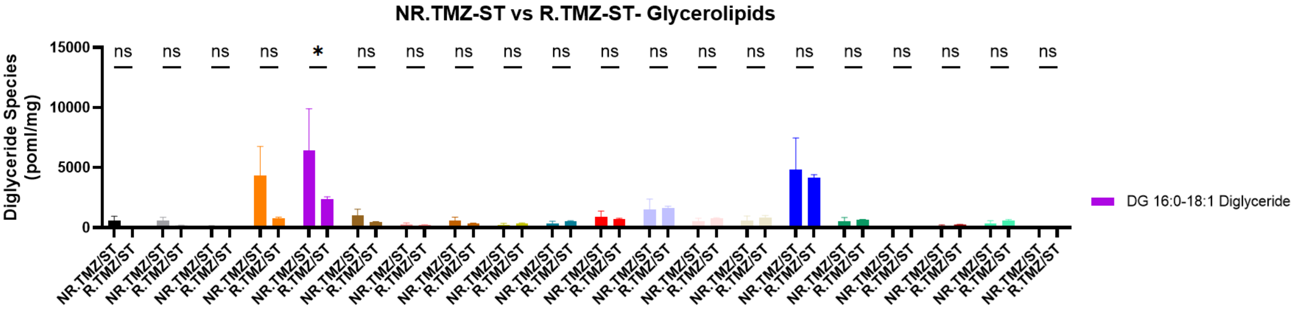
NR.TMZ-ST vs R.TMZ-ST- Glycerolipids
Diglyceride Species (poml/mg)
15000
10000
5000
0
ns
*
DG 16:0-18:1 Diglyceride
NR.TMZ/ST
R.TMZ/ST
NR.TMZ
R.TMZ

**S.**

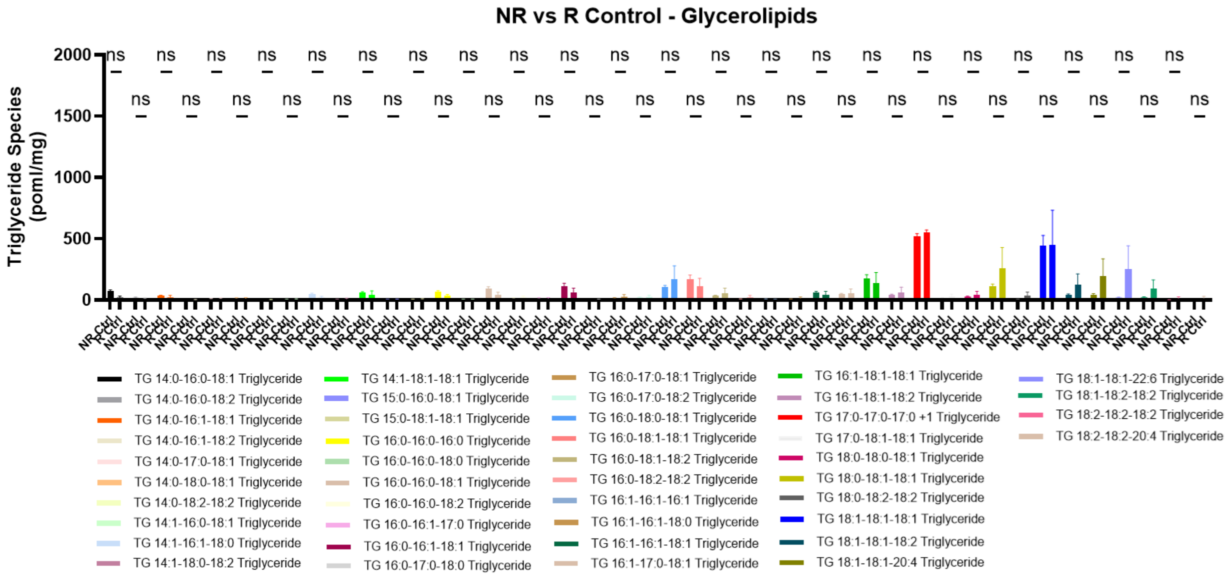
NR vs R Control - Glycerolipids
Triglyceride Species (poml/mg)
2000
1500
1000
500
0
ns
NR.Ctrl
R.Ctrl
TG 14:0-16:0-18:1 Triglyceride
TG 14:0-16:0-18:2 Triglyceride
TG 14:0-16:1-18:1 Triglyceride
TG 14:0-16:1-18:2 Triglyceride
TG 14:0-17:0-18:1 Triglyceride
TG 14:0-18:0-18:1 Triglyceride
TG 14:0-18:2-18:2 Triglyceride
TG 14:1-16:0-18:1 Triglyceride
TG 14:1-16:1-18:0 Triglyceride
TG 14:1-18:0-18:2 Triglyceride
TG 14:1-18:1-18:1 Triglyceride
TG 15:0-16:0-18:1 Triglyceride
TG 15:0-18:1-18:1 Triglyceride
TG 16:0-16:0-16:0 Triglyceride
TG 16:0-16:0-18:0 Triglyceride
TG 16:0-16:0-18:1 Triglyceride
TG 16:0-16:0-18:2 Triglyceride
TG 16:0-16:1-17:0 Triglyceride
TG 16:0-16:1-18:1 Triglyceride
TG 16:0-17:0-18:0 Triglyceride
TG 16:0-17:0-18:1 Triglyceride
TG 16:0-17:0-18:2 Triglyceride
TG 16:0-18:0-18:1 Triglyceride
TG 16:0-18:1-18:1 Triglyceride
TG 16:0-18:1-18:2 Triglyceride
TG 16:0-18:2-18:2 Triglyceride
TG 16:1-16:1-16:1 Triglyceride
TG 16:1-16:1-18:0 Triglyceride
TG 16:1-16:1-18:1 Triglyceride
TG 16:1-17:0-18:1 Triglyceride
TG 16:1-18:1-18:1 Triglyceride
TG 16:1-18:1-18:2 Triglyceride
TG 17:0-17:0-17:0 +1 Triglyceride
TG 17:0-18:1-18:1 Triglyceride
TG 18:0-18:0-18:1 Triglyceride
TG 18:0-18:1-18:1 Triglyceride
TG 18:0-18:2-18:2 Triglyceride
TG 18:1-18:1-18:1 Triglyceride
TG 18:1-18:1-18:2 Triglyceride
TG 18:1-18:1-20:4 Triglyceride
TG 18:1-18:1-22:6 Triglyceride
TG 18:1-18:2-18:2 Triglyceride
TG 18:2-18:2-18:2 Triglyceride
TG 18:2-18:2-20:4 Triglyceride

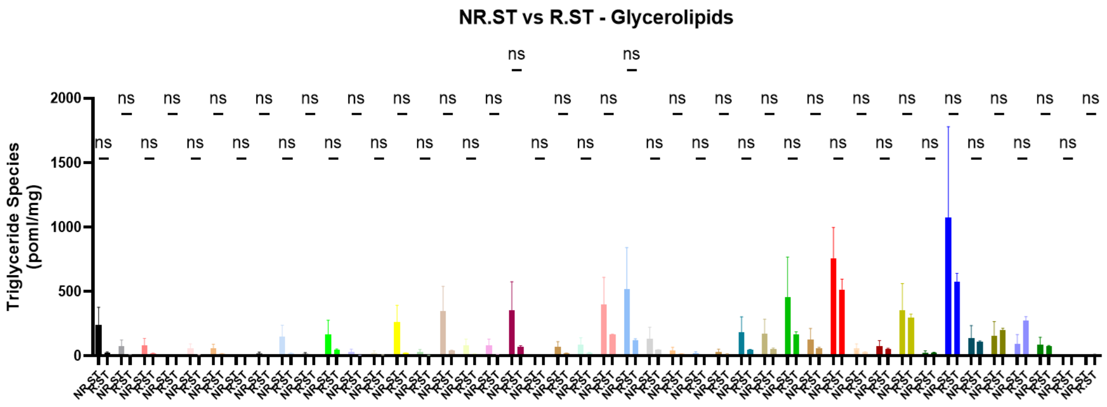
NR.ST vs R.ST - Glycerolipids
Triglyceride Species (poml/mg)
2000
1500
1000
500
0

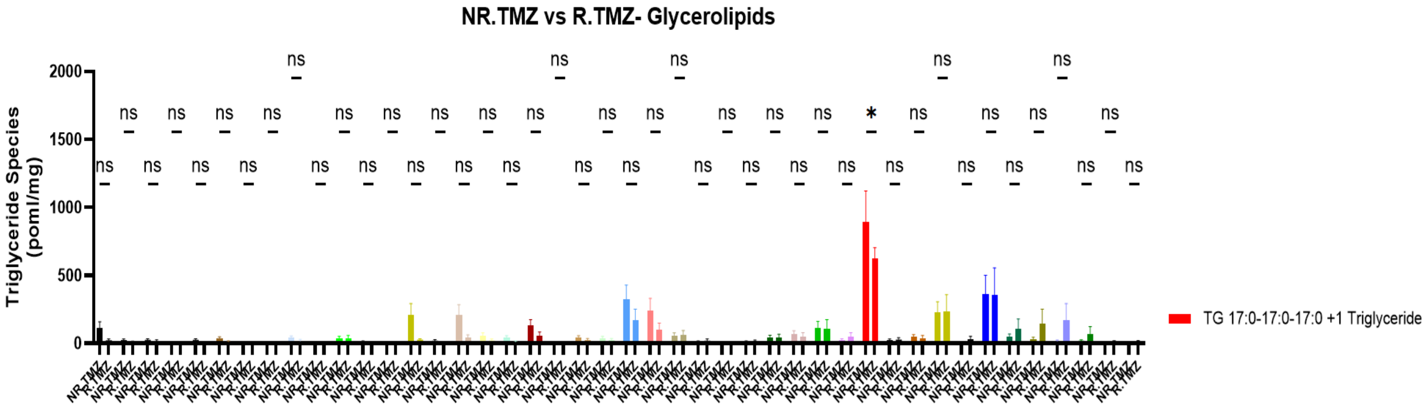
NR.TMZ vs R.TMZ- Glycerolipids
Triglyceride Species (poml/mg)
2000
1500
1000
500
0
TG 17:0-17:0-17:0 +1 Triglyceride

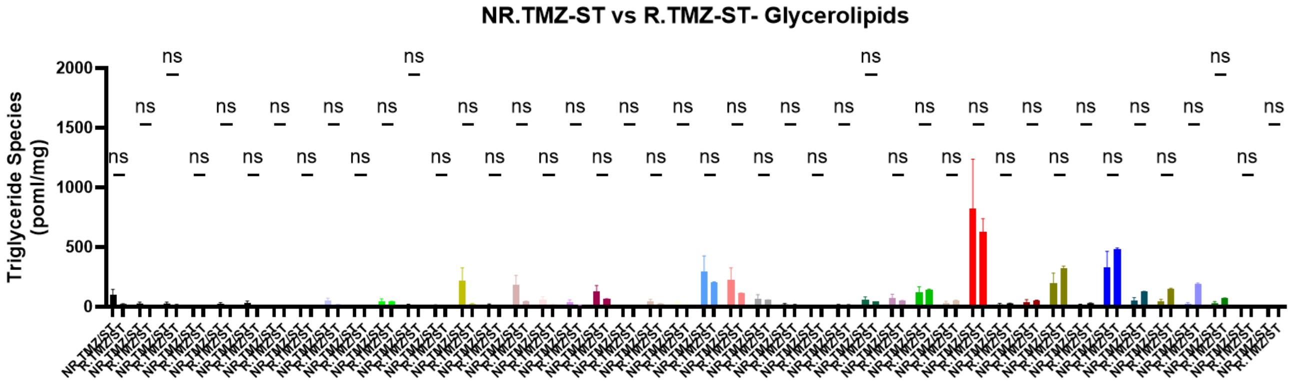
NR.TMZ-ST vs R.TMZ-ST- Glycerolipids
Triglyceride Species (poml/mg)
2000
1500
1000
500
0

**T.**

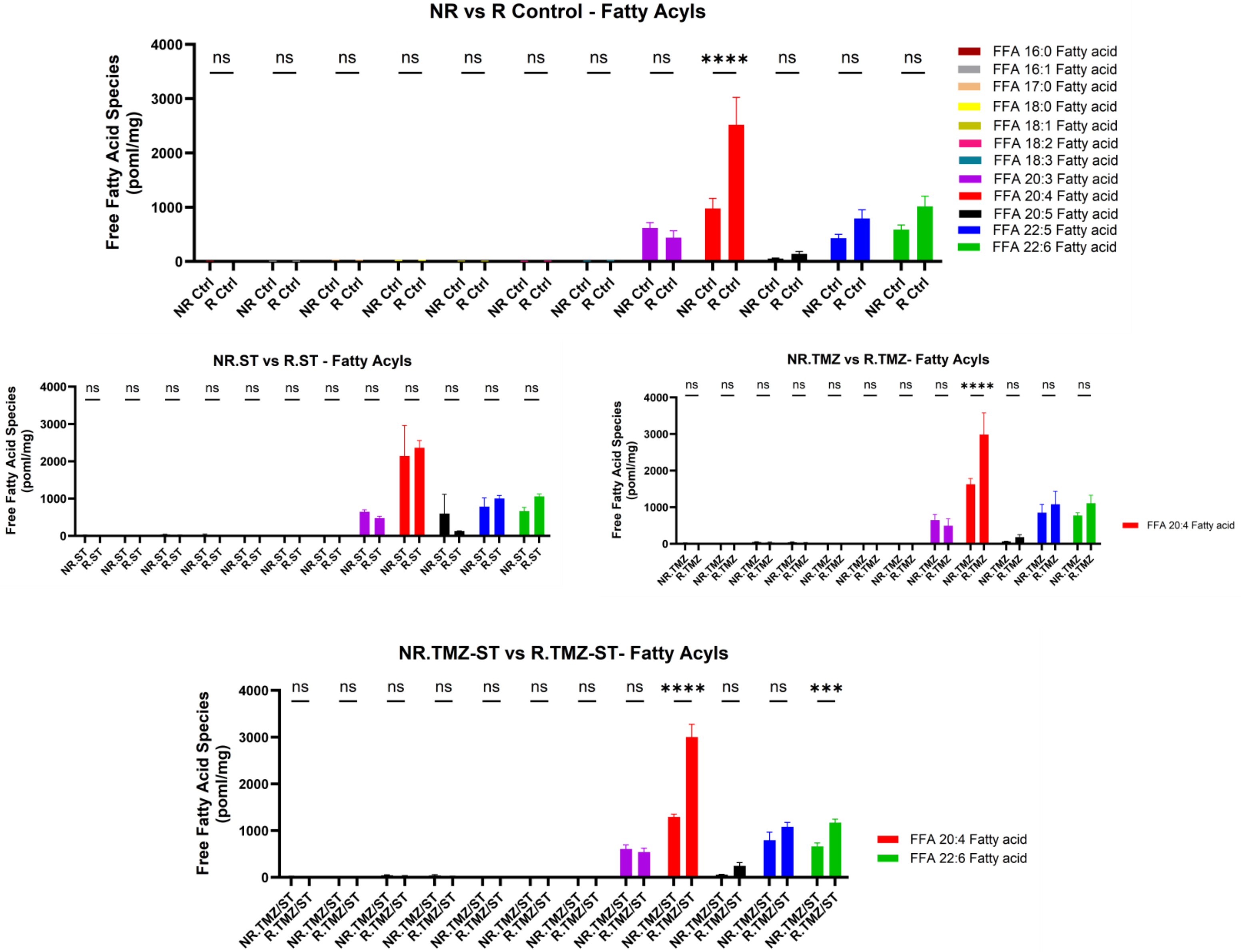
NR vs R Control - Fatty Acyls
Free Fatty Acid Species (poml/mg)
FFA 16:0 Fatty acid
FFA 16:1 Fatty acid
FFA 17:0 Fatty acid
FFA 18:0 Fatty acid
FFA 18:1 Fatty acid
FFA 18:2 Fatty acid
FFA 18:3 Fatty acid
FFA 20:3 Fatty acid
FFA 20:4 Fatty acid
FFA 20:5 Fatty acid
FFA 22:5 Fatty acid
FFA 22:6 Fatty acid
NR Ctrl
R Ctrl
NR.ST vs R.ST - Fatty Acyls
NR.ST
R.ST
NR.TMZ vs R.TMZ- Fatty Acyls
NR.TMZ
R.TMZ
FFA 20:4 Fatty acid
NR.TMZ-ST vs R.TMZ-ST- Fatty Acyls
NR.TMZ/ST
R.TMZ/ST
FFA 20:4 Fatty acid
FFA 22:6 Fatty acid

**U.**

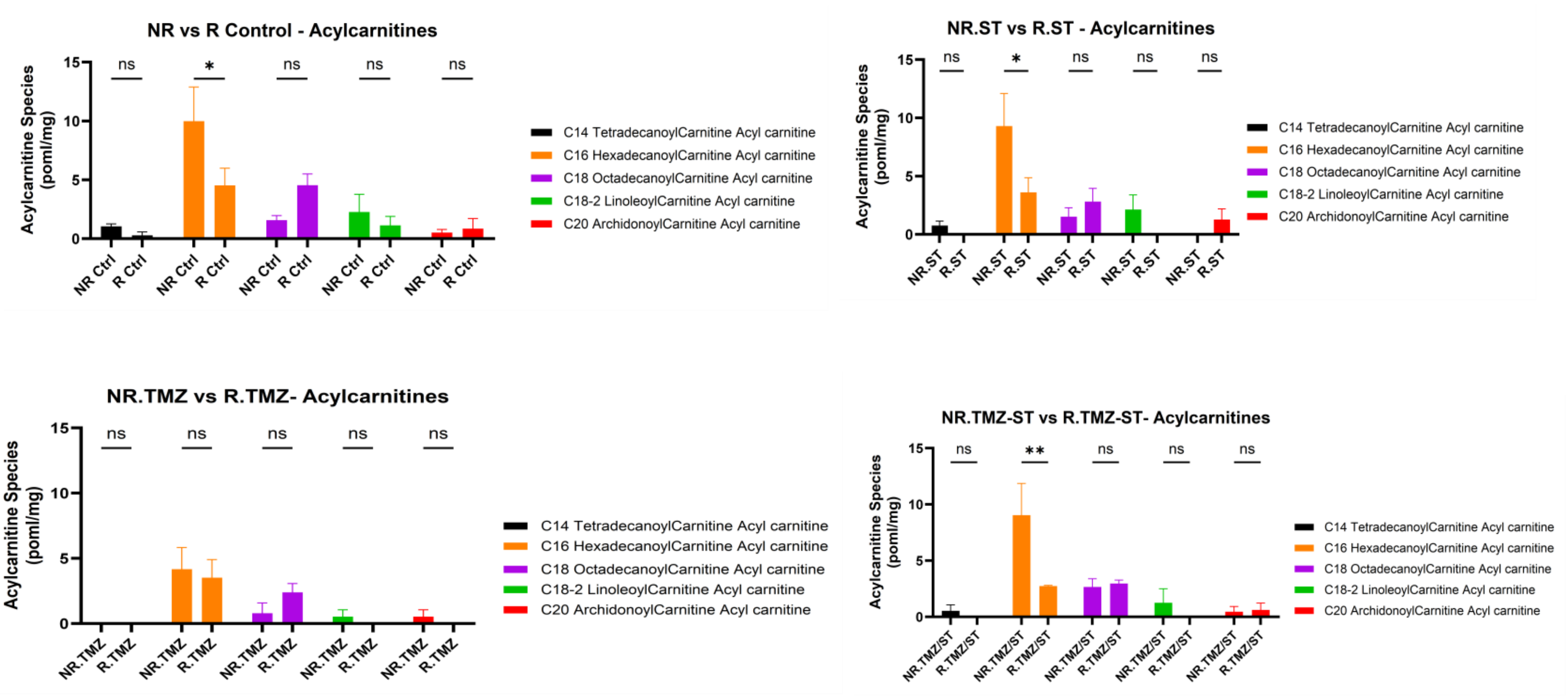

NR vs R Control - Acylcarnitines
Acylcarnitine Species (poml/mg)
ns
*
NR Ctrl
R Ctrl
NR.ST vs R.ST - Acylcarnitines
NR.ST
R.ST
NR.TMZ vs R.TMZ- Acylcarnitines
NR.TMZ
R.TMZ
NR.TMZ-ST vs R.TMZ-ST- Acylcarnitines
**
NR.TMZ/ST
R.TMZ/ST
C14 TetradecanoylCarnitine Acyl carnitine
C16 HexadecanoylCarnitine Acyl carnitine
C18 OctadecanoylCarnitine Acyl carnitine
C18-2 LinoleoylCarnitine Acyl carnitine
C20 ArchidonoylCarnitine Acyl carnitine


**V.**

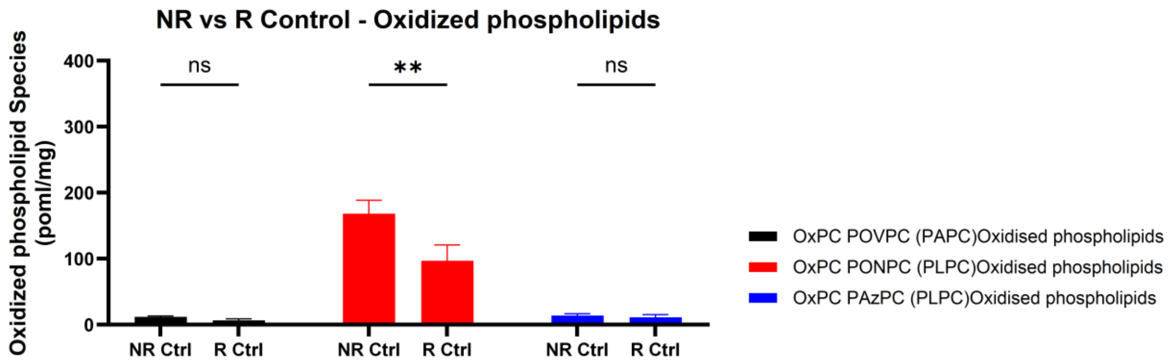

NR vs R Control - Oxidized phospholipids
Oxidized phospholipid Species (poml/mg)
400
300
200
100
0
ns
**
ns
NR Ctrl
R Ctrl
OxPC POVPC (PAPC)Oxidised phospholipids
OxPC PONPC (PLPC)Oxidised phospholipids
OxPC PAzPC (PLPC)Oxidised phospholipids

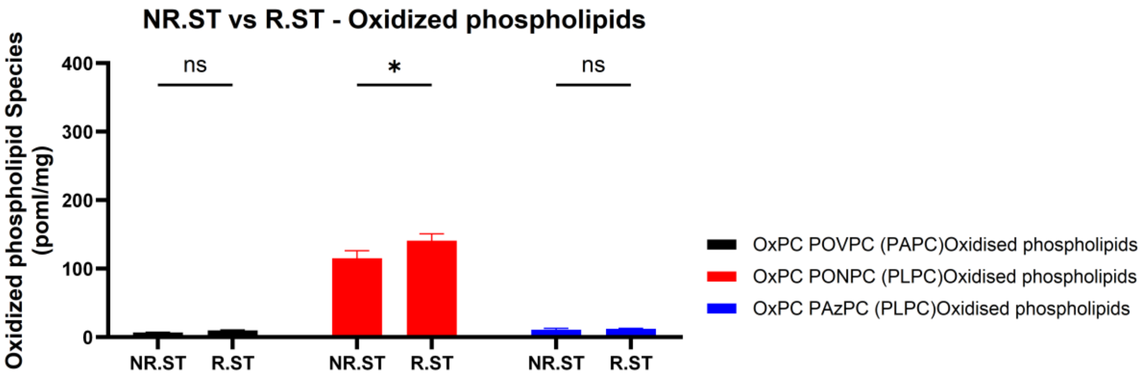


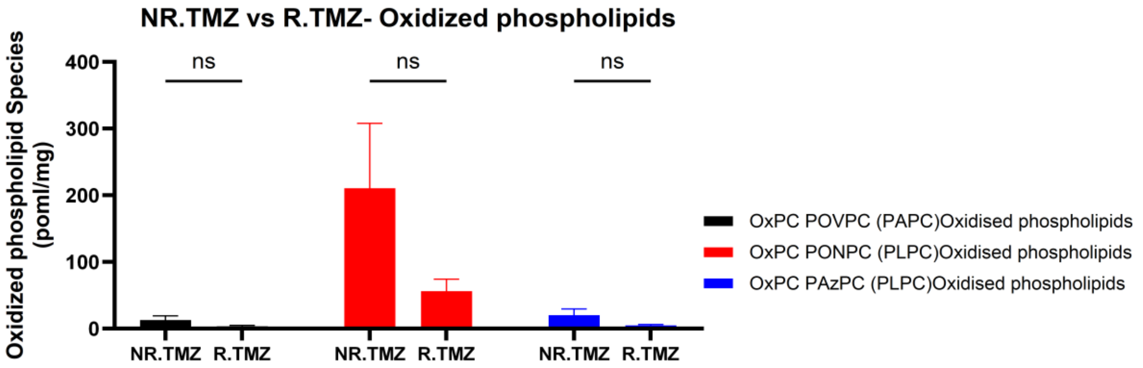


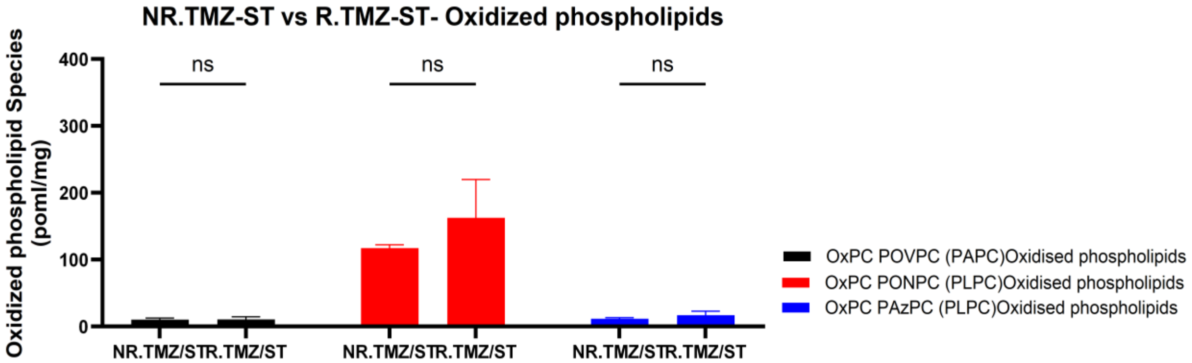


**Supplementary Figure S5.** Univariate lipidomic analysis reveals class-specific remodeling in TMZ-resistant glioblastoma cells. (A-V) Species-level comparisons between NR and R U251 glioblastoma cells under control, ST, TMZ, and TMZ-ST conditions for ceramides (A), dihydroceramides (B), dihexosylceramides (C), monohexosylceramides/cerebrosides (D), trihexosylceramides/globosides (E), gangliosides (F), sphingomyelins (G), phosphatidylcholines (H), ether/plasmalogen phosphatidylcholines (I), phosphatidylethanolamines (J), ether/plasmalogen phosphatidylethanolamines (K), phosphatidylinositols (L), phosphatidylserines (M), phosphatidylglycerols (N), lysophosphatidylcholines (O), lysophosphatidylethanolamines (P), cholesteryl esters (Q), diglycerides (R), triglycerides (S), free fatty acids (T), acylcarnitines (U), and oxidized phospholipids (V). Resistant cells show recurrent enrichment of lysophospholipid, selected sphingolipid/glycosphingolipid, and cholesteryl-ester species with treatment-dependent remodeling of glycerophospholipid, glycerolipid, fatty-acid, acylcarnitine, and oxidized-phospholipid pools. Each condition represents three independent biological replicates. Data are mean ± SD; statistical testing used one-way or two-way ANOVA with Tukey multiple-comparison testing as appropriate. *P<0.05, **P<0.01, ***P<0.001, ****P<0.0001; ns, not significant.

#### Supplementary Tables

##### Supplementary Table S1. Full-dimensional family-level separation statistics.

| Family | Features | PERMANOVA F | $R^2$ | P | Norm. centroid distance | Silhouette |
|---|---|---|---|---|---|---|
| PI | 17 | 23.91 | 0.521 | 0.0001 | 2.49 | 0.414 |
| PG | 4 | 21.28 | 0.492 | 0.0001 | 2.26 | 0.378 |
| LPC | 23 | 18.66 | 0.459 | 0.0001 | 1.96 | 0.374 |
| LPE | 7 | 18.09 | 0.451 | 0.0001 | 1.89 | 0.372 |
| Ether/plasmalogen PC | 18 | 17.74 | 0.446 | 0.0001 | 2.11 | 0.395 |
| PC | 30 | 15.37 | 0.411 | 0.0001 | 1.82 | 0.360 |
| SM | 20 | 14.28 | 0.394 | 0.0001 | 1.80 | 0.298 |
| Ether/plasmalogen PE | 19 | 14.27 | 0.393 | 0.0009 | 1.95 | 0.312 |
| PE | 18 | 10.64 | 0.326 | 0.002 | 1.57 | 0.289 |
| Globosides | 12 | 9.70 | 0.306 | 0.0001 | 1.42 | 0.274 |
| Cerebrosides | 6 | 8.57 | 0.280 | 0.002 | 1.49 | 0.295 |
| Diglycerides | 20 | 7.49 | 0.254 | 0.0003 | 1.56 | 0.349 |
| Free Fatty Acids | 12 | 5.50 | 0.200 | 0.0008 | 1.11 | 0.191 |
| Cholesteryl Esters | 20 | 4.90 | 0.182 | 0.002 | 1.38 | 0.202 |
| Triglycerides | 44 | 4.58 | 0.172 | 0.017 | 1.12 | 0.206 |
| Acylcarnitines | 5 | 4.16 | 0.159 | 0.015 | 0.94 | 0.170 |
| Phosphatidylserine | 7 | 4.12 | 0.158 | 0.049 | 1.04 | 0.173 |
| Gangliosides | 6 | 3.45 | 0.136 | 0.001 | 0.97 | 0.159 |
| Ceramides | 12 | 2.71 | 0.110 | 0.053 | 0.81 | 0.169 |

Statistics are calculated in the full log2-transformed, z-score-normalized feature space. Embedding figures are visualization tools and do not replace these full-dimensional statistics.

##### Supplementary Table S2. Family-level linear SVM performance across internal validation schemes.

| Family | Features | LOOCV MCC | LOCO MCC | LOTO MCC | LOTO balanced accuracy | LOTO MCC q |
|---|---|---|---|---|---|---|
| Globosides | 12 | 1.000 | 1.000 | 1.000 | 1.000 | 0.0353 |
| Ether/plasmalogen PE | 19 | 1.000 | 1.000 | 1.000 | 1.000 | 0.0353 |
| PE | 18 | 1.000 | 1.000 | 1.000 | 1.000 | 0.0353 |
| PC | 30 | 1.000 | 1.000 | 1.000 | 1.000 | 0.0353 |
| LPE | 7 | 1.000 | 1.000 | 1.000 | 1.000 | 0.0353 |
| LPC | 23 | 1.000 | 1.000 | 1.000 | 1.000 | 0.0353 |
| Triglycerides | 44 | 1.000 | 1.000 | 1.000 | 1.000 | 0.0353 |
| Ether/plasmalogen PC | 18 | 1.000 | 1.000 | 1.000 | 1.000 | 0.0353 |
| Cerebrosides | 6 | 1.000 | 1.000 | 1.000 | 1.000 | 0.0353 |
| Free Fatty Acids | 12 | 0.920 | 0.920 | 0.920 | 0.958 | 0.0353 |
| Ceramides | 12 | 1.000 | 0.920 | 0.920 | 0.958 | 0.0353 |
| Diglycerides | 20 | 0.920 | 0.920 | 0.920 | 0.958 | 0.0667 |
| Cholesteryl Esters | 20 | 0.920 | 0.920 | 0.920 | 0.958 | 0.0353 |

| | | | | | | |
|---|---|---|---|---|---|---|
| PG | 4 | 0.833 | 0.833 | 0.920 | 0.958 | 0.0353 |
| Phosphatidylserine | 7 | 0.920 | 0.920 | 0.920 | 0.958 | 0.0353 |
| SM | 20 | 1.000 | 0.920 | 0.920 | 0.958 | 0.0353 |
| PI | 17 | 1.000 | 0.775 | 0.775 | 0.875 | 0.0353 |
| Gangliosides | 6 | 0.775 | 0.707 | 0.602 | 0.792 | 0.0947 |
| Free Cholesterol | 1 | 0.500 | 0.585 | 0.500 | 0.750 | 0.0353 |
| Oxidized Phospholipids | 3 | 0.385 | 0.258 | 0.258 | 0.625 | 0.2857 |
| Acylcarnitines | 5 | 0.338 | 0.338 | 0.251 | 0.625 | 0.2700 |

LOOCV, leave-one-out cross-validation; LOCO, leave-one-condition-out; LOTO, leave-one-treatment-out. q values are Benjamini-Hochberg-adjusted MCC permutation P values based on 70 permutations per family. These are internal proof-of-concept analyses, not externally validated classifier estimates.

**Supplementary Table S3. Internal reduced-signature retention analysis.**

| Family | Full features | Retained k | Reduced LOTO MCC | Full-family LOTO MCC | Reduced-signature q |
|---|---|---|---|---|---|
| Cerebrosides | 6 | 2 | 1.000 | 1.000 | 0.0333 |
| Ether/plasmalogen PC | 18 | 2 | 1.000 | 1.000 | 0.0333 |
| Ether/plasmalogen PE | 19 | 1 | 1.000 | 1.000 | 0.0333 |
| Globosides | 12 | 1 | 1.000 | 1.000 | 0.0333 |
| LPC | 23 | 1 | 1.000 | 1.000 | 0.0333 |
| LPE | 7 | 3 | 1.000 | 1.000 | 0.0333 |
| PC | 30 | 1 | 1.000 | 1.000 | 0.0333 |
| PE | 18 | 1 | 1.000 | 1.000 | 0.0333 |
| Triglycerides | 44 | 4 | 1.000 | 1.000 | 0.0333 |
| Cholesteryl Esters | 20 | 2 | 1.000 | 0.920 | 0.0333 |
| PI | 17 | 1 | 1.000 | 0.775 | 0.0333 |
| Ceramides | 12 | 6 | 0.920 | 0.920 | 0.0333 |
| Diglycerides | 20 | 1 | 0.920 | 0.920 | 0.0333 |
| Free Fatty Acids | 12 | 4 | 0.920 | 0.920 | 0.0333 |
| PG | 4 | 1 | 0.920 | 0.920 | 0.0333 |
| Phosphatidylserine | 7 | 2 | 0.920 | 0.920 | 0.0333 |
| SM | 20 | 1 | 0.920 | 0.920 | 0.0333 |
| Gangliosides | 6 | 2 | 0.602 | 0.602 | 0.1579 |
| Free Cholesterol | 1 | 1 | 0.500 | 0.500 | 0.0333 |
| Oxidized Phospholipids | 3 | 1 | 0.354 | 0.258 | 0.2286 |
| Acylcarnitines | 5 | 1 | 0.333 | 0.251 | 0.2100 |

Reduced subsets were selected and evaluated within the same 24-sample dataset and therefore assess internal redundancy only. They should not be interpreted as validated species-level biomarkers. The current exported feature identifiers are analysis-pipeline identifiers and are not used here to assign biological lipid names.
